# Tissue-specific collagen hydroxylation at GEP/GDP triplets mediated by P4HA2

**DOI:** 10.1101/2023.01.25.524868

**Authors:** Dafné Wilhelm, Alison Wurtz, Hanane Abouhelfara, Guillaume Sanchez, Catherine Bui, Jean-Baptiste Vincourt

## Abstract

Collagen, the most abundant organic compound of vertebrate organisms, is a supramolecular, protein-made polymer. Details of its post-translational maturation largely determine the mechanical properties of connective tissues. Its assembly requires massive, heterogeneous prolyl-4-hydroxylation (P4H), catalyzed by Prolyl-4-hydroxylases (P4HA1-3), providing thermostability to its elemental, triple helical building block. So far, there was no evidence of tissue-specific regulation of P4H, nor of a differential substrate repertoire of P4HAs. Here, the post-translational modifications of collagen extracted from bone, skin, and tendon were compared, revealing lower hydroxylation of most GEP/GDP triplets, together with fewer other residue positions along collagen α chains, in the tendon. This regulation is mostly conserved in two distant homeotherm species, mouse and chicken. The comparison of detailed P4H patterns in both species suggests a dual mechanism of specificity. *P4ha2* expression is low in tendon and its genetic invalidation in the ATDC5 cellular model of collagen assembly specifically mimics the tendon-related P4H profile. Therefore, P4HA2 has a better ability than other P4HAs to hydroxylate the corresponding residue positions. Its local expression participates in determining the P4H profile, a novel aspect of the tissue specificities of collagen assembly.

**Data availability:** Proteomics data are available *via* ProteomeXchange with the identifier PXD039221. Reviewer account details:

login: MSV000091002_reviewer

password: P4HA2tendon.

## INTRODUCTION

Collagen is a fibrillar polymer providing a semi-rigid framework for the organization of multicellular organisms. It evolved as the most abundant organic compound in vertebrates and particularly in their connective tissues [1]. Collagen assembles from its elemental building block, a triple-helical protein [2]. Distinct types of collagen are encoded by corresponding genes, providing part of the mechanical diversity required for the specific functions of each tissue [3, 4]. However, most connective tissues are made primarily of type I collagen and yet have radically distinct mechanical properties. Along its biosynthesis, collagen undergoes an incomparably high rate of post-translational modifications (PTMs), including disulfide bridge formation and proteolytic cleavages (reviewed in [3]), as well as virtually collagen-specific PTMs targeting prolyl and lysyl residues. Oxidative deamination, hydroxylation and glycosylation of lysyl residues are critical in establishing the valency, structure, and residue position of intermolecular cross-links [5-9] and are viewed as major determinants of mechanical properties. Indeed, crosslinks differ between bone, skin, and tendon [6, 9, 10], which are considered prototypical connective tissues with radically distinct mechanical properties, though mostly made of type I collagen.

Quantitatively, however, most collagen PTMs consist in prolyl hydroxylation, about 150 prolyl residues being modified per collagen α chain [11]; they are divided into prolyl-3-hydroxylation (P3H), which targets prolyls standing on X positions of the collagen-typical GXY triplet, and prolyl-4-hydroxylation (P4H), which mostly targets those occupying the Y positions [12]. P3H, a relatively rare PTM, exhibits tissue as well as developmental specificities [13-15] and is believed to favor inter-triple helix interactions [16-18]. In contrast, P4H, the most abundant PTM of collagen and therefore, the most abundant PTM at all in vertebrates, is mostly considered non-tissue-specific, because it plays a critical, ubiquitous function in thermodynamically stabilizing the procollagen triple helix assembly [19, 20], partly by promoting inter-helix hydrogen bounding [21].

However, even when considering a single collagen type extracted from a single tissue, P4H is very heterogeneous [22], in the sense that many potential P4H-susceptible residues are found under both hydroxylated and non-hydroxylated molecular states, a point that has complicated detailed P4H description. Even though the overall P4H rate of collagen chains needs to remain relatively constant to allow adapted thermal stability, its positional heterogeneity leaves space for potential drastic residue-directed regulations, which could impact the mechanics of the resulting network but also the interactions of collagen with other molecules, including cellular receptors [23], thereby contributing to tissue-specific functions. P4H is catalyzed by 3 Prolyl-4-hydroxylases, P4HA1 to 3. P4HA1 is mostly ubiquitous, while P4HA2 expression is high in the skeleton and in the vascular endothelium but low in most other tissues [24, 25] and P4HA3 is expressed in many tissues but at very low levels [26]. Their respective functional specificities remain elusive [27-29]. In mice, the *P4ha1* knock-out is embryonically lethal [30], while that of *P4ha2* has no major phenotype [30] and that of *P4ha3* has not been reported yet. Here, we provide the first evidence that tendon collagen exhibits a specific P4H pattern, which we find is recapitulated in the ATDC5 model specifically upon *P4ha2* genetic invalidation.

## RESULTS

### Mouse tendon exhibits lower hydroxylation of a subset of collagen peptides, compared to bone and skin

To better describe the potential tissue specificities of prolyl-hydroxylation, collagen was acid-extracted from mouse bone, skin, and tendon tissues. Such crude extracts mostly contain the collagen-typical α, β and γ chains (figure 1. A). The differential hydroxylation of their tryptic peptides was investigated through a dedicated shot-gun, relative quantitative proteomics approach with adapted database interrogation methods. At this point, for the sake of automating, the method identified peptide sequences as well as the nature and number of their PTMs, but not the residue positions of PTMs. This resulted in the identification of 108 unique peptides encompassing more than 90% of the mature type I collagen α chain polypeptides. Almost each peptide was identified under several molecular states differing by their number of hydroxylations and therefore, the 108 unique peptides corresponded to 300 molecular states. Remarkably, 20% of the molecular states identified in this approach demonstrated significant quantitative changes between tissues (figure 1. B). Tendon exhibited the highest number of enriched species, 24, of which 22 were identified as under-hydroxylated versions of 16 different peptide sequences. Among these, 13 were redundantly represented among the molecular states found significantly enriched in bone and skin, (the second largest class of changes, figure 1. B), this time reflecting increased hydroxylation. Typical peptide examples are shown in figure 1. C-D to better explain the repartition of molecular states per unique peptide sequence. The peptide corresponding to residue positions 177 to 209 of type I collagen α1 chain precursor (COL1A1) is found under a total of 6 molecular states, 4 of which are significantly enriched in tendon compared to bone and skin, while the 2 others exhibit the contrary enrichment (figure 1. C). The four molecular states enriched in tendon are those with the lowest hydroxylation levels while the two states enriched in bone and skin are those with the highest hydroxylation levels, all of them together reflecting lower hydroxylation of the corresponding peptide in tendon. The same logic applies to the peptide corresponding to residue positions 106 to 138 of type I collagen α2 chain precursor (COL1A2, figure 1. D). Other peptide cases are shown in supplemental data S1. These data indicate a tendon-specific under-hydroxylation of 16 type I collagen peptides.

**Figure 1.**
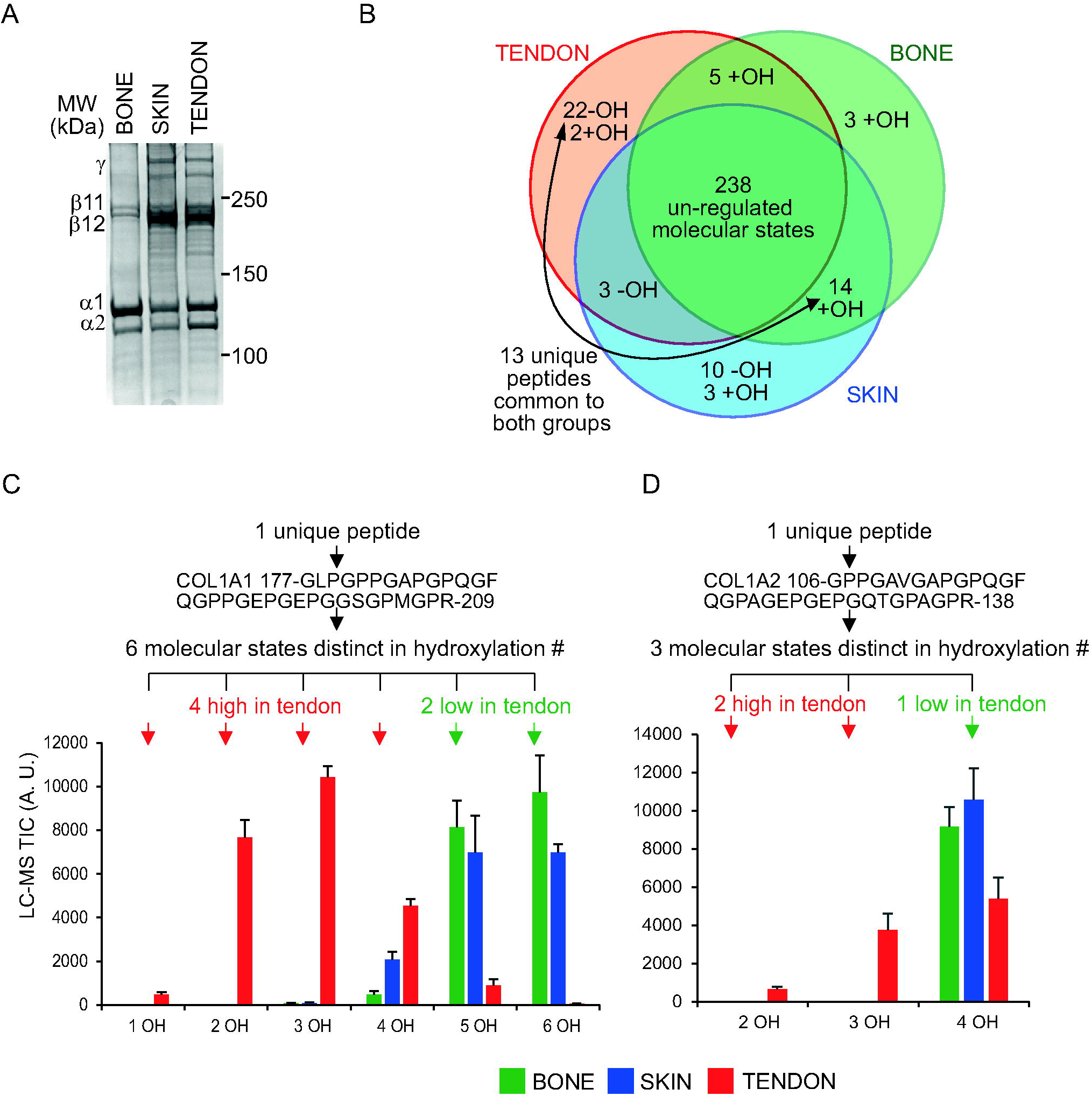
Mouse tendon exhibits lower hydroxylation of a subset of collagen peptides, compared to bone and skin. (A) collagen acid extracts from bone, skin and tendon (2 μg each) analyzed by SDS-PAGE coupled to Coomassie staining. Annotations on the right indicate the positions of α, β and γ collagen chains. (B) Venn diagram summarizing tissue-specific differences in tryptic collagen peptide hydroxylation. The number of molecular states significantly increased in each sub-group is annotated; +OH, over-hydroxylated; -OH, under-hydroxylated. (C-D) Typical examples of hydroxylation profiles of two peptides originating from COL1A1 and COL1A2, the two precursor chains of type I collagen, respectively. Each peptide, of which the sequence and boundaries are given, is represented by distinct molecular states distinguished by their number of post-translationally added hydroxyl groups, as indicated on the x-axis, and the histograms show the respective total ion chromatogram values, + SD. In (C), t-test p-values for tendon versus bone and skin together: 1.2E-07, 8.4E-11, 2.6E-14, 3.7E-05, 5.7E-06, 5.3E-06, respectively for the states with 1, 2, 3, 4, 5, or 6 hydroxylations. In (D), t-test p-values for tendon versus bone and skin together: 4.7E-08, 1.3E-07, 3.7E-4, respectively for the states with 2, 3, or 4 hydroxylations. Throughout this figure, n=4 per tissue.

### Recurrent under-hydroxylation of GEP/GDP triplets in mouse tendon collagen

In order to determine at which residue positions hydroxylation is reduced in tendon, compared to bone and skin, MS/MS spectra of all hydroxylation states of the peptides of interest were inspected in detail (supplemental data S1. A). In addition, crude quantitative data obtained in each tissue and for each hydroxylation variant of each peptide were combined per peptide (supplemental data S1. B) to calculate the average peptide hydroxylation number, per tissue. Both aspects are summarized in table 1. All tendon down-regulated hydroxylations occur on prolyl residues in Y position of GXY triplets, consistent with the notion that P4H, rather than P3H, is involved [12]. Remarkably, for 13 out of 16 peptides, the tendon-down-regulated hydroxylations occur at either a GEP or GDP triplet (table 1). More specifically, the single GDP occurring within the whole type I collagen molecule, as well as 14 out of 20 GEPs within both COL1A1 and COL1A2, exhibit low hydroxylation in tendon. This is the first evidence of a tissue-regulated P4H profile of collagen.

**Table 1.**
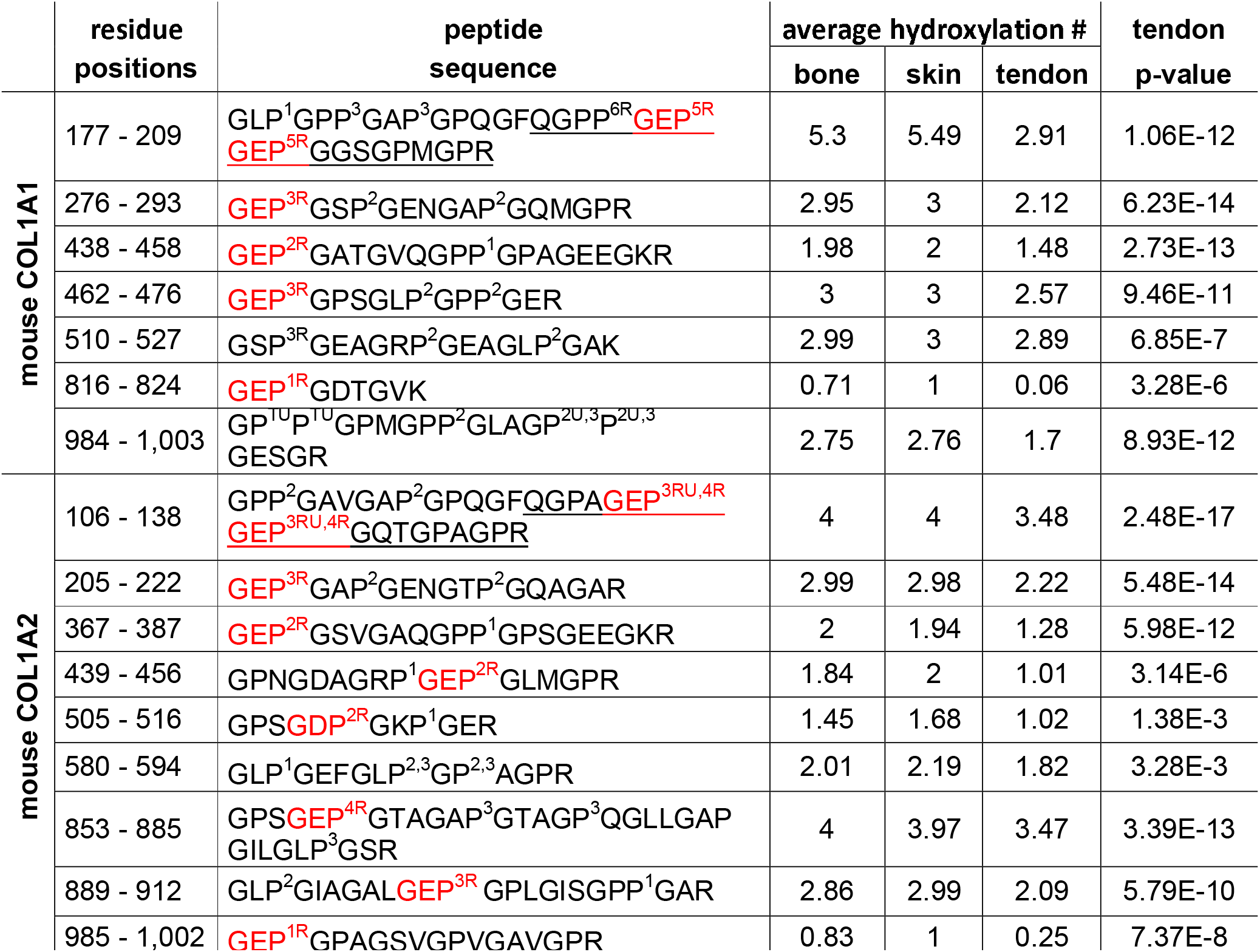
Mouse tendon exhibits low hydroxylation at most collagen GEP/GDP triplets. Table summarizing positions and fold regulations of hydroxylation. Peptides are classified per their amino acid boundaries, referring to their positions in the α1 (COL1A1) or α2 chain (COL1A2) of type I mouse collagen precursor. The amino acid sequence is given, with numbers indicating the order of preferential hydroxylation according to MS/MS fragmentation of the distinct molecular states: ^T^tendon-specific hydroxylated position. ^U^ : the exact residue position of hydroxylation could not be determined between similarly annotated residues. ^R^ : positions at which hydroxylation is found lower in tendon collagen. Underlined: sequences within which hydroxylated positions were determined from a chymotrypsin + trypsin digest. GEP/GDP triplets are highlighted in red. The average hydroxylation number of each peptide in each tissue is calculated from total ion chromatograms. t-test p-value is given for tendon versus others (n=4 per tissue).

### Comparison of the tendon-related P4H profiles of mouse and chicken suggests a dual mechanism of specificity

To address the evolutive representativity of the decreased GEP/GDP hydroxylation found in mouse tendons, the P4H profiles of chicken leg tendon and skin were compared to one another using the same methods (details are shown in supplemental data S2). Overall, findings in the two species correlate positively (figure 2. A, Pearson r = 0.64). In details, some of the GEP/GDPs are not conserved between the two species and in most such cases, the peptide version possessing a GEP/GDP exhibits tendon-related under-hydroxylation while that lacking a GEP/GDP does not (figure 2. A), re-enforcing the notion that GEP/GDPs determine the underlying process. However, table 1 also shows peptides that lack a GEP/GDP and yet are under-hydroxylated in tendon, indicating additional determinants. The peptide with boundaries 510-527 from COL1A1 is remarkable in this respect: it contains a GEP in chicken and the corresponding P residue is its 4^th^ potential hydroxylation site (figure 2. B, top part), being less hydroxylated in tendon. In mouse, the same peptide is fully conserved except for the GEP, which is substituted with GEA and therefore the peptide has only 3 potential hydroxylation sites, but one of them is still less hydroxylated in tendon (figure 2. B, bottom part). Therefore, the P4H profile of tendon relies on partial specificity towards GEP/GDP triplets, combined with other mechanisms targeting, at a larger scale, particular regions of procollagen.

**Figure 2.**
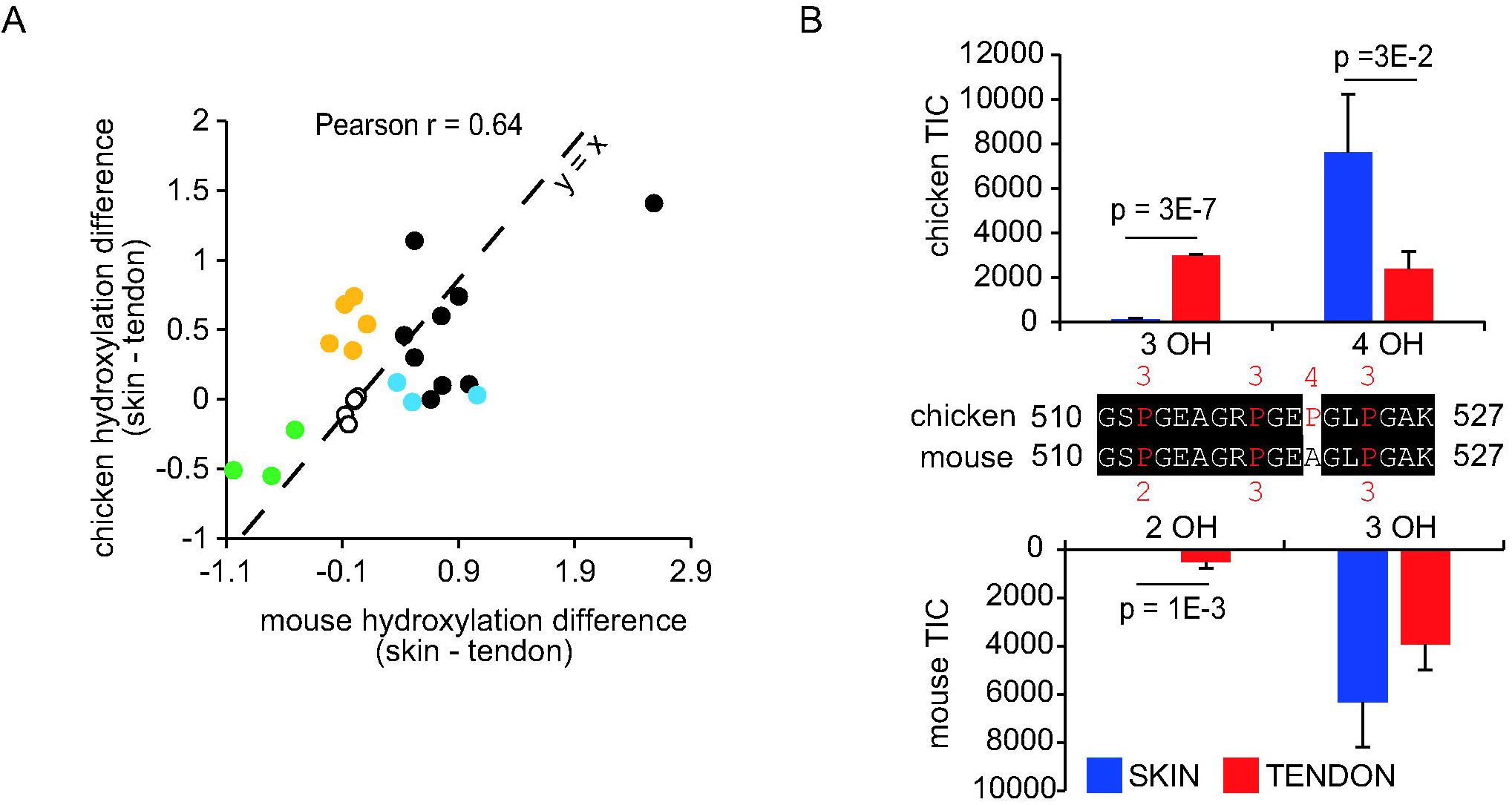
Partial conservation of the P4H profile of tendon collagen between mouse and chicken. (A) Comparison of the influence of tendon synthesis, as compared to skin, between mouse and chicken. Black dots: peptides found under-hydroxylated both in mouse and chicken tendon, independently, compared to skin; blue dots: peptides found under-hydroxylated in mouse tendon but lacking a GEP in chicken; orange dots: peptides found under-hydroxylated in chicken tendon but lacking a GEP in mouse. Empty dots: peptides exhibiting no hydroxylation regulation between tissues; green dot: peptide found under-hydroxylated in skin compared to tendon, in both species. (B) Hydroxylation states of peptide 510-527 from COL1A1 in mouse and chicken tendon and skin. The amino acid sequences are shown in the middle, with hydroxylation positions highlighted in red and attached numbers indicating the preferential order of residue hydroxylation. Histograms show the respective total ion chromatogram average values of each peptide version, + SD, versions differing in their number of hydroxylated residues (in mouse, n=4; in chicken, n=3). Details for each chicken peptide are shown in supplemental data S2.

### *P4ha2* exhibits low mRNA expression in tendon, suggesting its involvement in GEP/GDP hydroxylation

To assess which P4HA isoenzyme [24-26] might be responsible for GEP/GDP hydroxylation, the expression levels of *P4ha1-3* mRNA were explored in mouse reference tissues (figure 3. A). The skin expresses all three *P4ha* genes at their lowest levels among the three tissues, suggesting that this tissue requires low overall P4HA expression. The tendon expresses the highest level of *P4ha1* and the second, close to the highest, level of *P4ha3*, disfavoring the idea that any of these enzymes are involved in GEP/GDP hydroxylation. In contrast, *P4ha2* is mostly expressed in the bone and its level is insignificantly different between the tendon and the skin. Therefore, P4HA2 was hypothesized to exert GEP/GDP hydroxylation.

**Figure 3.**
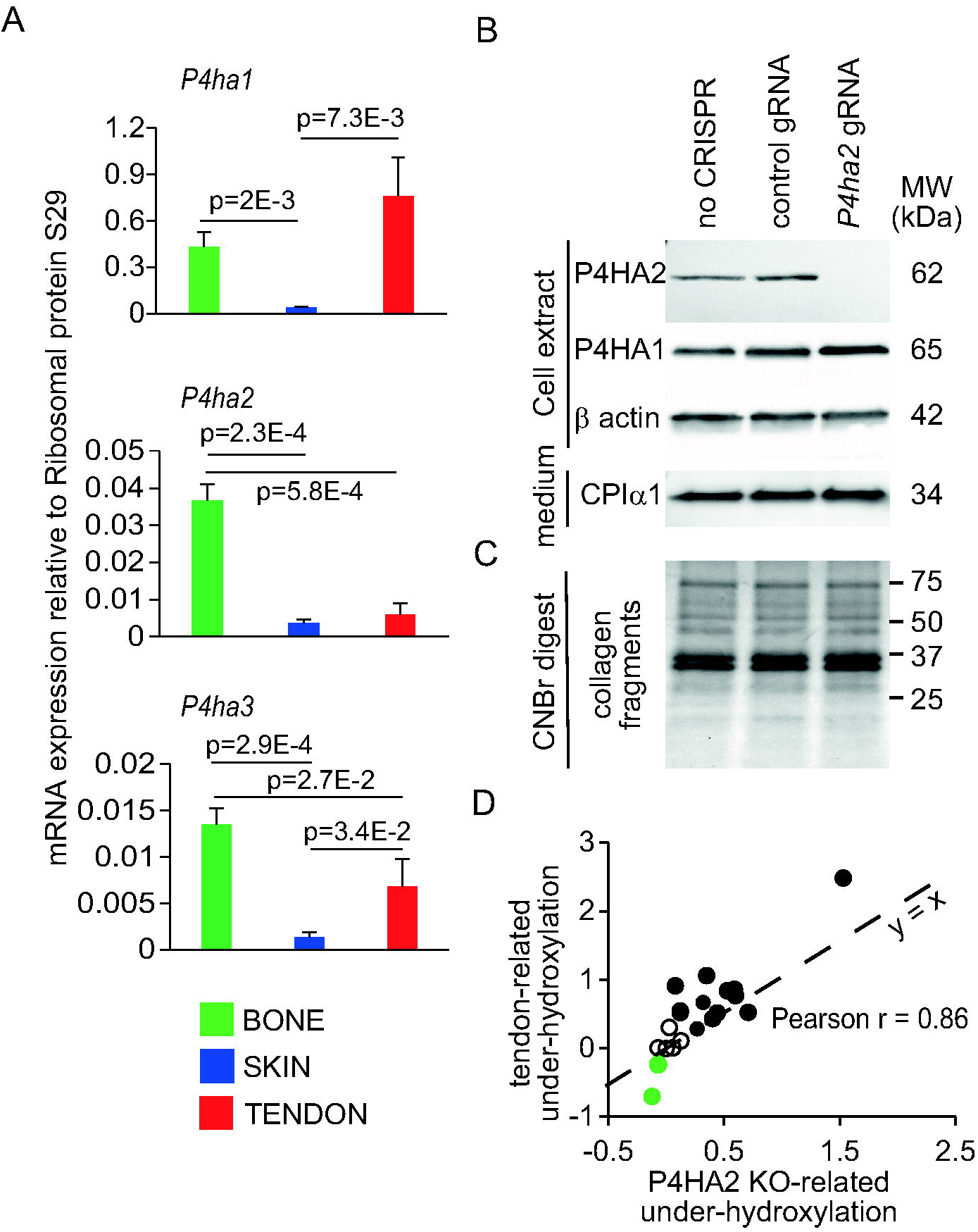
Invalidation of *P4ha2* in the ATDC5 model mimics the tendon-related P4H profile. (A) Average tissue expression of *P4ha1-3*, reported to that of the reference ribosomal protein S29, + SD. (B) Western blotting analysis of distinct cell fractions extracted from control versus *P4ha2* knock-out cells. CP I α1: C-propeptide of type I collagen α1 chain. (C) P4HA2 ablation does not affect the collagen content or assembly, as reflected by the electrophoretic profile of CNBr digests, analyzed by SDS-PAGE coupled to Coomassie blue staining. (D) Correlation analysis between the influence of tendon synthesis (using bone and skin as mixed reference) and that of P4HA2 deficiency in ATDC5 cells (versus control ATDC5 cells), over the average hydroxylation number of distinct collagen peptides. Black dots: peptides found under-hydroxylated in tendon. Empty dots: peptides found unregulated between tissues. Green dots: peptides found under-hydroxylated in skin. Throughout this figure, n=3.

### Invalidation of *P4ha2* in the ATDC5 model mimics the P4H profile of tendon collagen

To question the specific function of P4HA2, the corresponding gene was invalidated in the widely used chondrogenesis model relying on ATDC5 cells [31, 32], which we found recapitulate many aspects of collagen biogenesis and assembles massive type I collagen [33]. mRNA studies show robust expression of *P4ha1-3* in control ATDC5 cells (compared to mouse tissues) and near extinction of *P4ha2* upon dedicated CRISPR/Cas9-mediated invalidation, without changes of *P4ha1* or *P4ha3* expression, cell growth or survival (supplemental data S3). Western blot analysis confirms effective loss of P4HA2 expression without changes in either P4HA1 expression or type I collagen secretion (figure 3. B). Culture supernatant analysis does not reveal any changes in the secreted protein levels (not shown). Quantitatively, crosslinked collagen production is not affected by P4HA2 depletion (figure 3. C). However, the collagen produced by these cells exhibits significant hydroxylation reduction of most peptides found under-hydroxylated in tendon (supplemental data S4), including GEP/GDP-containing and -lacking ones. This is illustrated by a comparative analysis, for distinct peptides, of the influence of tendon biosynthesis (compared to bone and skin) to that of P4HA2 depletion in ATDC5 cells (figure 3. D, Pearson r = 0.86). These results support the idea that the low expression of P4HA2 in the tendon determines the specific collagen P4H profile in this tissue, implying that P4HA2 is, better than P4HA1 and P4HA3, capable of hydroxylating the corresponding positions. However, one could also hypothesize that the loss of P4HA2 decreases overall P4H activity, thereby resulting in low hydroxylation of these residues if these are the least favorable substrates by all P4HAs.

### P4HA2 deficiency is specific in mimicking the tendon-related P4H profile

To address this issue, the ATDC5 model was treated with sub-lethal doses of nickel, which, among other effects, depletes ascorbic acid, thereby affecting all enzymatic hydroxylation reactions [34], including P3H and P4H. At the dose used here, nickel results in minimal (yet, significant) growth reduction (figure 4. A), does not perturb procollagen secretion (figure 4. B), but virtually abolishes mature collagen assembly (figure 4. C), consistent with insufficient overall P4H. Indeed, all collagen peptides exhibit decreased hydroxylation to variable degrees upon nickel treatment (supplemental data S5). Peptides found under-hydroxylated in *P4ha2*-invalidated cultures are not less hydroxylated upon nickel exposure than others (figure 4. D). Furthermore, several of them are less, or no more, affected by nickel than by P4HA2 deficiency (figure 4. D, compare on which side of the dashed line the coordinates are located). Finally, when highly influenced by nickel (such as peptide 889-912 from COL1A2, figure 4; D), their corresponding hydroxylation profile is very heterogeneous (figure 4. E, top histograms), indicating that multiple hydroxylation sites are under-used, while *P4ha2* invalidation affects isolated residues (figure 4. E, bottom part). Overall, the P4H profile in *P4ha2*-invalidated cells does not reflect global P4H inhibition, but rather, the better ability of P4HA2 (compared to P4HA1 and 3) at hydroxylating specific residue positions that define the tendon-related P4H profile of collagen.

**Figure 4.**
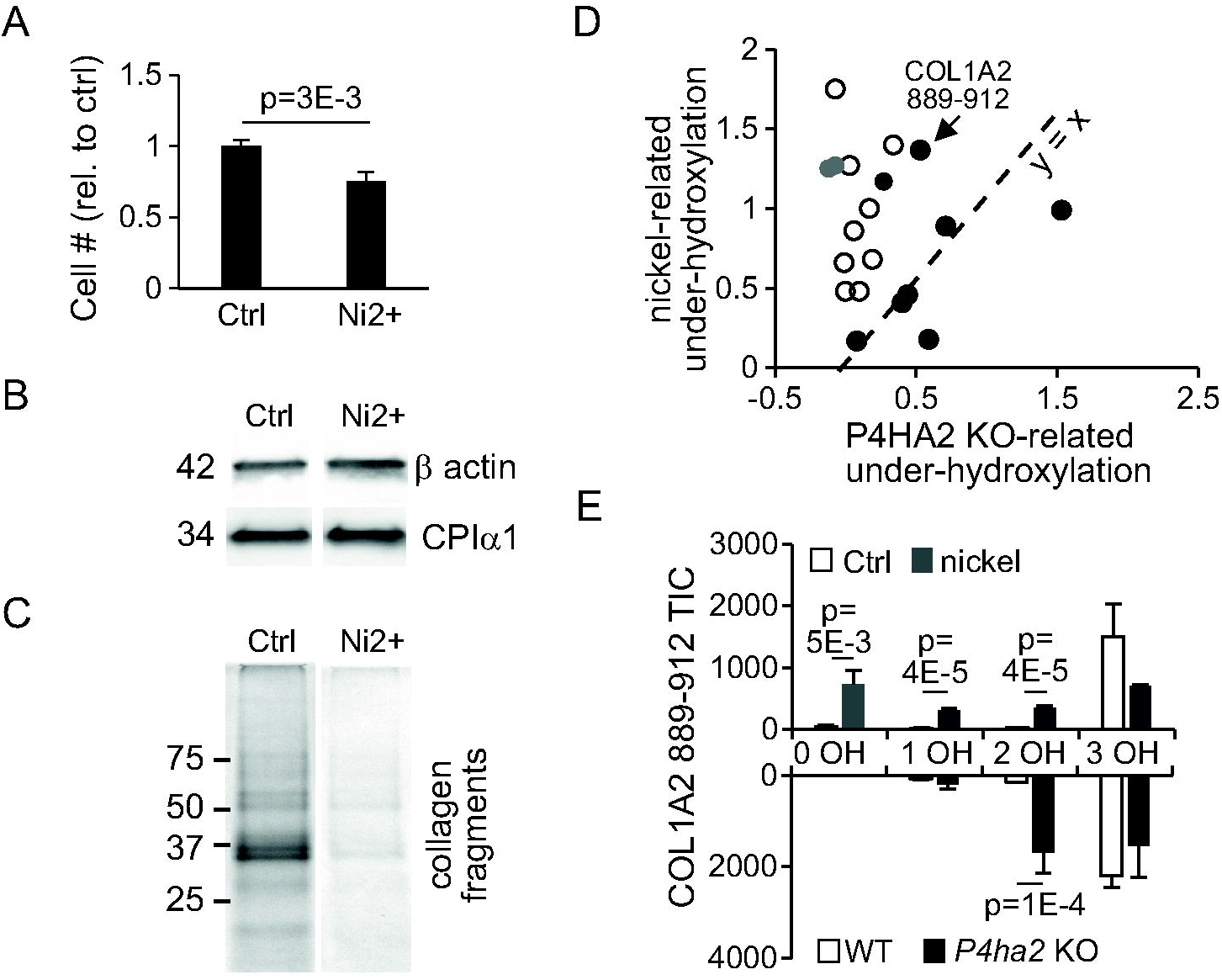
General inhibition of P4HA 1 to 3 does not specifically affect GEP/GDP hydroxylation. (A) Influence of nickel (100 μM) over the cell number, expressed as average, relative to control, +SD. (B) Western blotting analysis of actin in the intracellular fraction and of CPI α1 (the C-propeptide of type I α1 procollagen chain), in the secreted fraction. (C) Effect of nickel over collagen assembly, as revealed by SDS-PAGE coupled to Coomassie staining of the CNBr digest. (D) Comparison of the influence of nickel administration to that of P4HA2 deficiency, both in ATDC5 cells, over the average hydroxylation number of distinct collagen peptides (measured from culture supernatant samples). Black dots: peptides found under-hydroxylated in tendon. Empty dots: peptides found unregulated between tissues. Gray dots: peptides found under-hydroxylated in skin. (E) Total Ion Chromatogram values, expressed as average + SD, for each hydroxylation state of peptide 889-912 from COL1A2 measured in nickel-treated and *P4ha2* knock-out ATDC5 cells, versus controls, respectively. Details of other collagen peptides are in supplemental data S5. Throughout this figure, n =3.

## DISCUSSION

Bone, skin, and tendon, are made mostly of type I collagen and yet have radically distinct mechanical properties, inherent to their differential functions. A strong part of the distinction between them resides in their collagen crosslinking [5-9] and more modestly in their collagen P3H [35]. P4H, despite its quantitatively major contribution to collagen PTMs, has not, so far, been considered a potential contributor to the regulation of collagen biomechanics, because its main, ubiquitous function, stabilizing the triple helix assembly, relies mostly on its high rate of occurrence along α chains. In the present study, we report that 20 % of the molecular states (corresponding to about 15 % of the unique peptides) collected after trypsin digestion of type I collagen exhibit significant quantitative differences between bone, skin and tendon, reflecting changes in P4H profiles (figure 1 & table 1). Of note, these numbers possibly remain under-estimated, due to technical limitations: LC-MS approaches have difficulties in separating distinct molecular states corresponding to a same peptide harboring the same number of hydroxyl groups at distinct positions, because they have exactly the same mass and very close chemical properties. Also, the present study relies on shot-gun analysis and therefore, does not address the relationship between P4H events occurring at residues contained in distinct tryptic peptides. Future analytical developments should aim at better describing the details of P4H to fully understand the subtleties of collagen maturation and assembly.

The largest and most robust class of tissue specificities identified here reflects a lower degree of hydroxylation in tendon than in skin and bone, affecting specific collagen peptides (figure 1). The vast majority of hydroxylations being down-regulated in tendon occur at GEP/GDP motifs (table 1 & supplemental data 2. C). This is conserved between two distant homeotherms (figure 2. A). In this sense, the tissue specificity of P4H is more conserved through the evolution of vertebrates, than that of P3H [14]. Of note, one of the peptides carrying a GEP, that undergoes less hydroxylation in tendon, was already reported to do so in an earlier study conducted in rat [15], but the finding was not discussed. Interestingly, hydroxylation at this position was found more independent than P3H on the age of animals [15]. Therefore, the regulation of P4H is not only better conserved but also more stable than P3H through the lives of homeotherms, suggesting that it may play a more critical role in establishing tissue mechanical properties.

Investigations in the ATDC5 model demonstrate that P4HA2, one of three P4HA isoenzymes, is required to optimally hydroxylate the corresponding residues (figure 3). We could not evidence any other phenotypic consequence of *P4ha2* invalidation in ATDC5 cells (figure 3, supplemental data 3 and further interpretable from the available proteomics data). The general inhibition of all three P4HAs results in a very distinct hydroxylation profile (figure 4), suggesting that P4HA2 is better able than P4HA1 and P4HA3 of hydroxylating GEP/GDPs. P4HA3 is poorly characterized yet, but P4HA1 and P4HA2 were so far believed to possess no intrinsic GXP substrate specificity [27, 29]. Indeed, most GEP/GDPs exhibit significant, but not binary regulation of hydroxylation, in *P4ha2*-invalidated versus control cells (figure 3. F and supplemental data S3). Therefore, P4HA1&3 retain some ability in hydroxylating these positions, though with a lower efficacy. Furthermore, most, but not all GEP/GDPs exhibit hydroxylation regulation (summarized in supplemental data S6). Finally, hydroxylation at positions others than GEP/GDPs are also found modulated (table 1, figure 2. B, supplemental data S4). Overall, the nature of the X residue within the GXP typical motif is insufficient to determine the specificity of P4HA2. P4HAs possess a Peptide Substrate Binding (PSB) domain, N-terminal to the catalytic domain, which largely influences their interaction with collagen peptides [36]. The PSB of P4HA2 preferentially binds PXGP motifs, where X is not a proline [28], while that of P4HA1 targets PPG repeats [36]. Therefore, the PSBs of both isoenzymes bind to distinct regions of procollagen. Recently, the structure of a P4HA2 truncation combined to its functional Protein Disulfide Isomerase partner was solved and suggested that the binding of the PSB to a given region of the procollagen molecule could influence the interaction of the catalytic domain with other, spatially related regions, [37]. Still, the substrate acceptability is likely to depend on its own affinity towards the catalytic domain, which, in the light of the present study, in the case of P4HA2, favors GEP/GDP motifs more than in the case of P4HA1 and 3. A mechanism combining the influence of the PSB and the better acceptability of GEP/GDP as substrates by the catalytic domain would explain both the bias towards GEP/GDPs and its non-exhaustivity. The present findings will facilitate further differential, mechanistic characterization of P4HAs, critical contributors to tissue elaboration [38] also involved in cancer progression (reviewed in [39]).

*P4ha2* mutant mice exhibit no remarkable pathology [27]. Therefore, GEP/GDP hydroxylation is not critical. However, independent *P4ha2* mutations were found linked to human myopia [40, 41], suggesting that it may exert subtle functions. While P4H promotes triple helix stability in general [19-21], in the vicinity of acidic residues, it might have other consequences, yet to be characterized, that are tissue-regulated based on the expression of P4HA2, which is relatively specific [24, 25]. Beyond, the present, unexpected finding, that P4H is subject to positional regulation in specific tissues of homeotherms highlights the possible fundamental function of P4H positioning in tuning the mechanical and functional properties of the collagen network. Its full range of action in this regard deserves further investigation.

## METHODS

### Collagen extraction and normalization

C57BL/6N mice were bred in the animal facility of JANVIER Labs. Tissues (femur, skin and tail tendon) were dissected from 4, 7-weeks old, male mice and stored at - 80°C prior to processing. Collagen was extracted from crude tissues. Chicken legs were obtained from a local supermarket and directly processed. Extraction steps were carried out at 4°C unless stated otherwise. For mouse bone, one femur was cut into three equal pieces and the medial part was kept for extraction. For skin, pieces of about 0.1 cm^2^ were cut on the shaved back of mice or on chicken legs. For chicken tendon, the posterior tendon of the quadriceps was dissected, and its medial part was used. As for mouse tendon, in the original study, a whole single tail tendon was used per extraction. The original experiment described in figure 1 was performed with 4 replicate extractions from each tissue. The experiment on chicken tissues was performed with 3 replicates per group. However, in the light of results, another triplicate extraction was performed from the posterior tendon of mouse quadriceps and the hydroxylation profile was verified to resemble that of tail tendon (not shown). In the follow-up, all tissues were processed similarly. Samples were transferred to 2 ml Eppendorf tubes, washed 5 times for 30 min each in 1.5 ml PBS, 1% triton and complete protease inhibitors (Roche), demineralized three times, for several days each in 0.5 M EDTA, defatted in chloroform/methanol (75:25, v/v), rinsed 3 times in water, 5 minutes each, speed vac-dried for 1 hour, transferred to new tubes and weighted. Collagen was extracted in 0.5 M acetic acid, 1 ml/50 mg for 3 days at 4 °C on a roller and then heat-denatured at 75°C for 25 min, immediately cooled on ice, and centrifuged for 15 min at 15,000 g at 4°C. The supernatant was collected and aliquoted. The collagen-dedicated Bradford assay [42] was used to measure extracted amounts using a standard curve prepared from commercial rat tendon collagen and at the time of use, aliquots were diluted in 0.5 M acetic acid to normalize concentrations, speed vac-dried, and resuspended in dedicated buffers for downstream applications. 2 μg of each extract were loaded for SDS-PAGE/Coomassie control.

### Digestion and fractionation by nano-HPLC

10 μg of each extract was resuspended in 20 μl Urea buffer (6 M Urea, 50 mM Tris, pH 8.0) and sonicated for 2 min in a water bath sonicator. Samples were then diluted 10-fold in (Tris, 100 mM, pH 8.0, CaCl_2_ 10 mM) containing sequencing grade trypsin and digested at 37°C overnight; occasionally chymotrypsin was added for further, 6 hour digestion (both enzymes from Promega, 200 ng each). Samples were equilibrated to 0.1 % TFA, 2.5 % acetonitrile (ACN) and purified on a C18 column (C18 spin columns, Thermofischer) according to the supplier’s recommendations, dried, resuspended in 10 μl 2 % ACN, 0.1 % TFA and then transferred to nano-HPLC injection tubes. The peptides were fractionated by nano-HPLC an Ultimate3000 system equipped with a 20 μl sample loop, a C18 pepMap 100 desalting precolumn, and a 15 cm C18 pepMap RSLC fractionation column (all from Thermo). Samples (6.4 μl) were injected using μlpickup mode and eluted by a gradient of 2-45% ACN over 60 min at 300 nl/min. Fractions (340, 9 seconds each) were collected on a ProteineerFcII (Bruker) and eluted fractions were directly mixed on MTP-1536 BC target spots with α-cyano-4-hydroxycinnamic acid (both from Bruker).

### Generation of label-free relative quantitative data by LC-MALDI

LC-MALDI runs dedicated to quantitation were processed for all replicates using dedicated automatic methods piloted by WARP-LC software on an Autoflex speed MALDI-TOF/TOF mass spectrometer (Bruker) in MS mode in the 700-4500 mass range, using next-neighbor external calibration for all MALDI spots. WARP-LC MS runs were then exported to BAF files, which were used in ProfileAnalysis (Bruker) for alignment of MS ion chromatograms, using quantile normalization with the following filters: 50 ppm mass deviation; 2 min retention time deviation; minimum 3 counts within one biological group, generating relative quantitation bucket tables for all precursors passing the above criteria.

### MS/MS fragmentation

The above bucket tables were used in ProfileAnalysis to identify molecular states enriched (>1.5x, t-test p-value < 0.05) in either biological group, compared to the sum of the two other groups. The corresponding (retention time; mass) coordinates were used to generate scheduled precursor lists (SPL), used for targeted MS/MS in one or two runs of the corresponding biological groups. In addition, all masses with S/N > 20 were selected for untargeted MS/MS from 1 random run of each biological group. All MS/MS analyses were performed using a dedicated LIFT method on the Autoflex speed, from the same, already spotted LC runs.

### Peptide identification

The above LC-MS/MS data were exported to the Proteinscape server, merged, and identified through interrogation of the Swissprot mouse database, allowing 50 ppm deviation of MS data and 0.8 Da deviation of MS/MS data, with trypsin cut (or semi-trypsin cut when considering samples digested with chymotrypsin plus trypsin), one allowed miss-cleavage and considering proline hydroxylation, lysine hydroxylation, lysine deamination, lysine galactosylation, and lysine galactosylation plus glucosylation, together with methionine oxidation, requiring Mascot scores above 20. This resulted in a peptide identity FDR < 2 % using the random decoy strategy. At this point, residue positions of evidenced modifications were not taken into account.

### Identification of tissue-regulated peptide molecular states

Quantitative data generated in ProfileAnalysis were exported to proteinscape and attached to the identified peptides above, providing lists of identified peptides with relative quantitative calculations of each tissue versus the average of other tissues ratio and the corresponding t-test p-values, highlighting molecular states (without the notion of PTM residue position at this point) exhibiting tissue-specific regulations.

### Curation of PTM residue positions

Crude MS/MS spectra corresponding to the molecular states of interest were manually annotated, based on the selected fragments automatically selected during Mascot interrogation, to clarify whether the residue positions provided by Mascot were truly ascertained based on the fragments or not. This procedure was considered required as in several cases, as Mascot tended to select random positions among several possibilities of similar probabilities.

### Relative quantification of individual molecular states between tissues

The bucket tables generated as described above were exported to an excel file and total ion chromatogram values corresponding to all molecular states of a given peptide of interest in each sample data were collected based on their (retention time; mass) and used as crude relative quantification data. For each molecular state, averages and standard deviations were calculated per experimental group. For mouse samples, unpaired t-tests were calculated comparing all samples within each tissue to the sum of all samples from the two others.

### Calculation of the average hydroxylation number per peptide in each tissue

From the tables mentioned in the previous section, the average hydroxylation number of each peptide was calculated for each sample, using the formula: Σ(Qx*X)/Σ(Qx), where X is the number of hydroxylations of a given molecular state, ranging from 0 to the maximum observed for the given peptide and Qx is the corresponding total ion chromatogram (TIC) value. Results were averaged per group. Unpaired t-tests were calculated comparing all tendon samples to the sum of all bone and skin samples together.

### Cell culture and genome edition

The ATDC5 model was used strictly as described in [33], except that for genome editing, 48 hours after plating, cells were inoculated overnight with 5.10^7^ adenoviral constructs purchased from Vectorbuilder, transducing expression of a Cas9-GFP fusion protein together with selected gRNA, as follows: Controls were inoculated with a single-guided adenoviral CRISPR/Cas9 construct targeting human *Hexose-6-Phosphate Dehydrogenase* (gRNA sequence: CACCTGCGAAGGTCGGCGTC). Genetic invalidation of *P4ha2* was induced similarly using the following gRNA sequence: AGATCAGCTGCCGACCCCGA. Cells were allowed to reach confluency, the medium being changed every day, before insulin stimulation. 3-4 independent infections were used per experimental group as specified. For nickel exposure, NiCl_2_ was administered at 100 μM during the whole insulin stimulation. Differentiation was stopped on day 14 in all experiments. These experiments were repeated 3 times independently.

### Gene expression analysis

From the same mouse tissues as those used for collagen studies, total RNA was extracted using TRIzol Plus RNA Purification Kit (Thermo Fisher Scientific). From ATDC5 cells, total RNA was isolated using RNeasy mini kit (Qiagen). In both cases, purified RNA was quantified using a NanoDrop One spectrophotometer (Thermo Fisher). 500 ng of total RNA were reverse-transcribed using M-MLV reverse transcriptase (Invitrogen) with random hexamer primers, according to the manufacturer’s instructions. Real-time quantitative PCR was performed using the following primers: S29_For, GGAGTCACCCACGGAAGTT; S29_Rev, GCCTATGTCCTTCGCGTACT; P4HA1_FOR, AAACAACAGGCTGGAAATGG; P4HA1_REV, GCTTTGGTAGTGGGAATGGA; P4HA2c_FOR, TTACGCAGAGAAGGACCTGG; P4HA2c_REV, CAGGCCAGTCTGTGTTCAAC, P4HA3_FOR: GACCAATTCCAGCCCCTACC; P4HA3_REV: GCTGATGCGGTACTCCACTT, with SYBRGreen (Qiagen) on a StepOneplus instrument (ABSciex) and analysis was based on the ΔΔCT method, using *S29* as reference.

### Protein fractionation and sample preparation from ATDC5 cells

These aspects were managed mostly as described in [33] except that the intracellular fraction protein content was measured using the Bradford assay [43].

### SDS-PAGE and western blotting analysis

Intracellular and soluble-secreted proteins were Deoxycholate/Trichloroacetic acid– precipitated and separated by SDS-PAGE on gradient 4-20% PROTEAN TGX Stain-Free Gels (Bio-rad) and proteins were transferred to polyvinylidene difluoride membranes within Trans-Blot Turbo Transfer Packs (Bio-rad) with Trans-Blot Turbo transfer system. Antibodies against the C-propeptides of type I procollagen as well as the corresponding HRP-conjugated secondary antibodies were described elsewhere [44]. Anti-P4HA1: HPA026593, anti-P4HA2: HPA027824, anti-βactin: A3854, all from Sigma. Blots were developed with the enhanced chemiluminescence system (Bio-rad) and acquired on a Chemidoc XRS+ imager (Bio-rad). Acetic acid extracts and aliquoted CNBr-digests were speed-vac-dried and separated on SDS-PAGE as explained above. Gels were stained with Brilliant blue Coomassie overnight, washed in pure water several times, and scanned on a Chemidoc XRS+ imager (Bio-rad).

## ABBREVIATIONS

COL1A1: (α1 chain of type I collagen)
COL1A2: (α2 chain of type I collagen)
CNBr: (cyanogen bromide)
P3H: (Prolyl-3-Hydroxylation)
P4H: (Prolyl-4-Hydroxylation)
P4HA: (Prolyl-4-Hydroxylase Alpha subunit)
PSB: (Peptide Substrate-Binding)
PTM: (post-translational modification)
TIC: (total Ion Chromatogram)

## ACKNOWLEDGMENTS

The proteomics core facility of UMS 2008 UL-CNRS-INSERM IBSLor acknowledges the funding from CPER IT2MP_BIOS.

## DECLARATION OF INTEREST

none.

### Formatting of funding sources

This research did not receive any specific grant from funding agencies in the public, commercial, or not-for-profit sectors.

