## supplemental data S1 for "Tissue-specific collagen hydroxylation at GEP/GDP triplets mediated by P4HA2"

**Supplemental data S1. Detailed identification and relative quantitative data for peptides found underhydroxylated in mouse tendon.** (A) MS/MS spectra allowing determination of the residue positions of hydroxylations from mouse collagen. Crude MS/MS spectra of interest were mannually annotated, based on automated annotations obtained from the proteinscape server following Swissprot interrogation, in order to clarify the determinability of each proposed position. The locations of b and y ions are indicated on the spectra in red. The corresponding breaks are shown at the top and bottom of the corresponding peptide sequence, respectively. * indicates the positions of hydroxylations in the peptide sequence. (B) Detailed relative quantification data for peptides found under-hydroxylated in tendon collagen. Each panel shows the data obtained for a given peptide, the sequence and boundaries of which are given at the top. Graphs report on the absolute MS signal obtained for all identified versions of a given peptide. Statistics are not reported here due to their complexity but they are summarized in table 1.

A

COL1A1 191 – 209    F.QGPPGEP*GEP*GGSGPMGPR.G

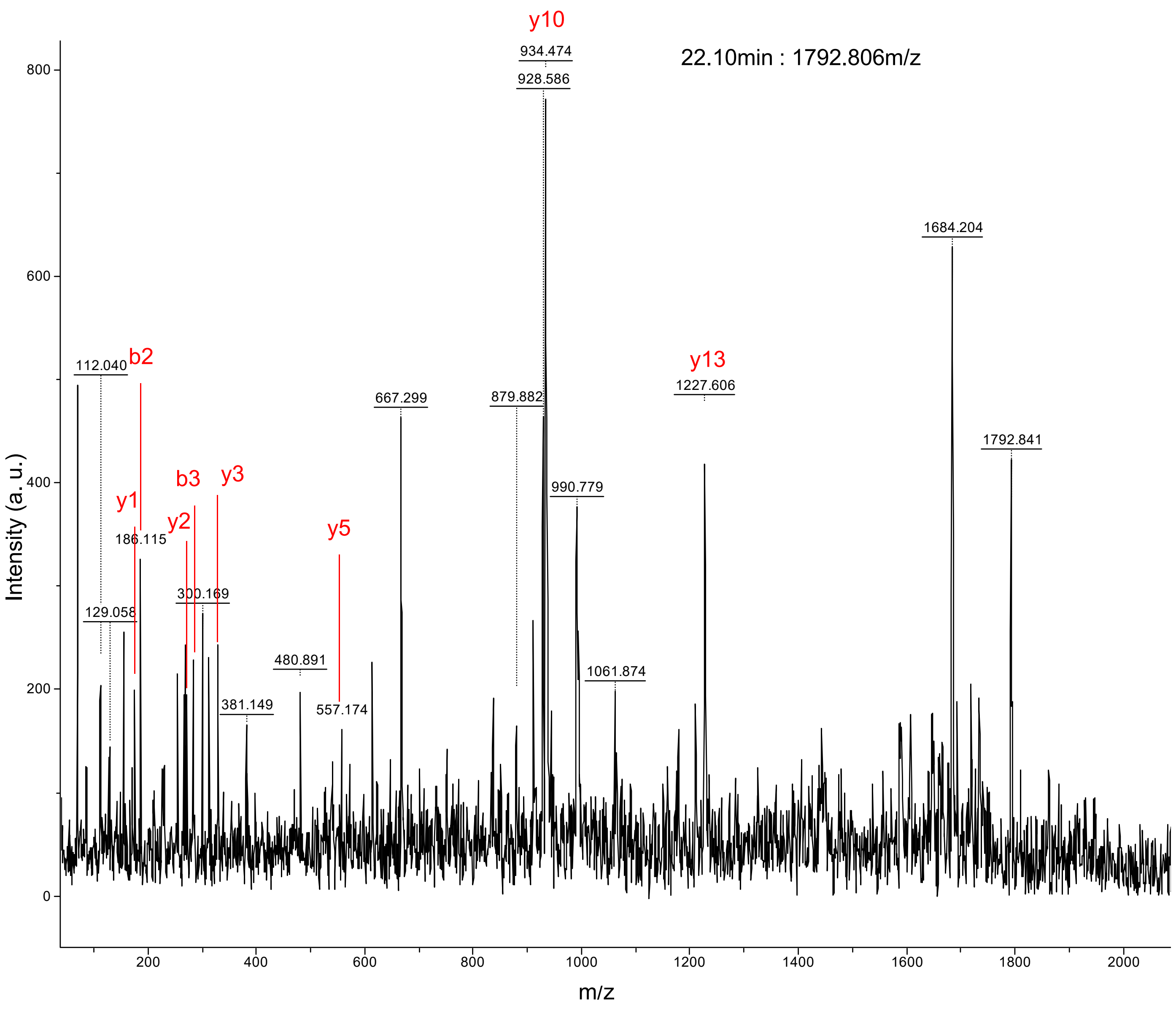

A (continued)

COL1A1 191 – 209 F.QGPP*GEP*GEP*GGSGPMGPR.G

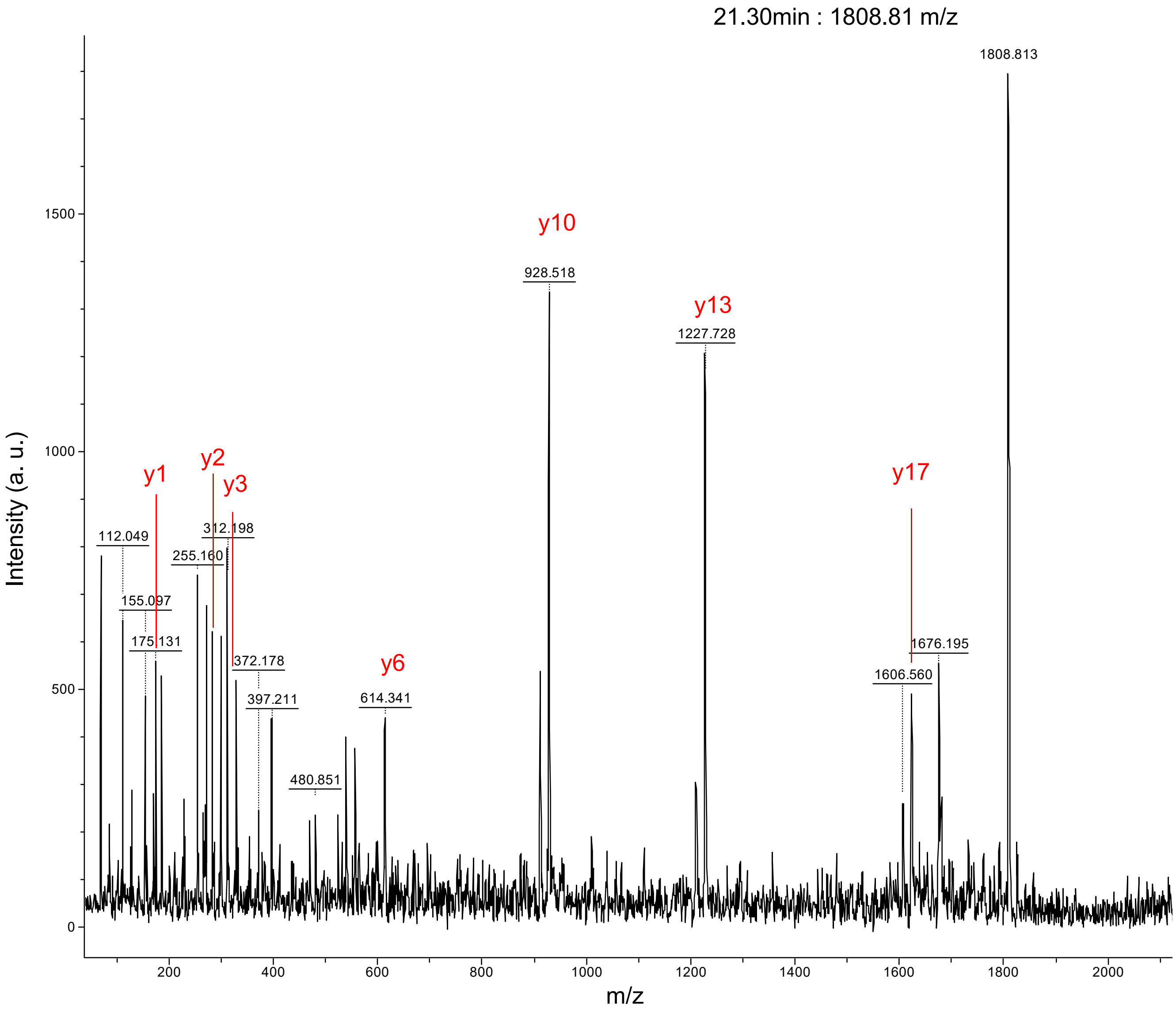

A (continued)

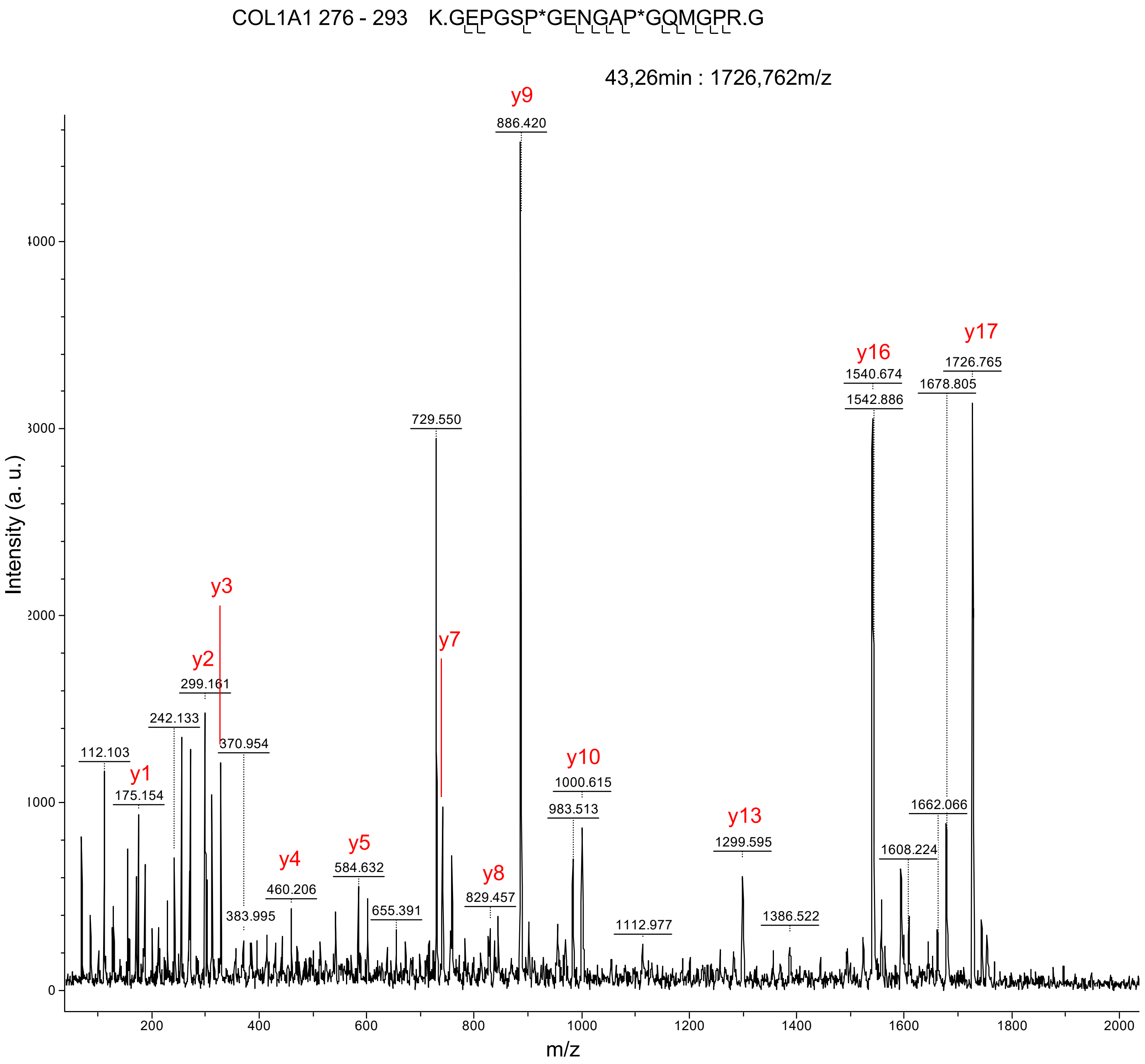
COL1A1 276 - 293 K.GEPGSP*GENGAP*GQMGPR.G
43,26min : 1726,762m/z
Intensity (a. u.)
m/z
y9
886.420
729.550
y16
1540.674
1542.886
y17
1726.765
1678.805
y3
y7
y2
299.161
242.133
370.954
112.103
y1
175.154
y10
1000.615
983.513
1662.066
y13
1299.595
1608.224
y5
584.632
y4
460.206
y8
829.457
383.995
655.391
1112.977
1386.522
0
1000
2000
3000
4000
200
400
600
800
1000
1200
1400
1600
1800
2000

A (continued)

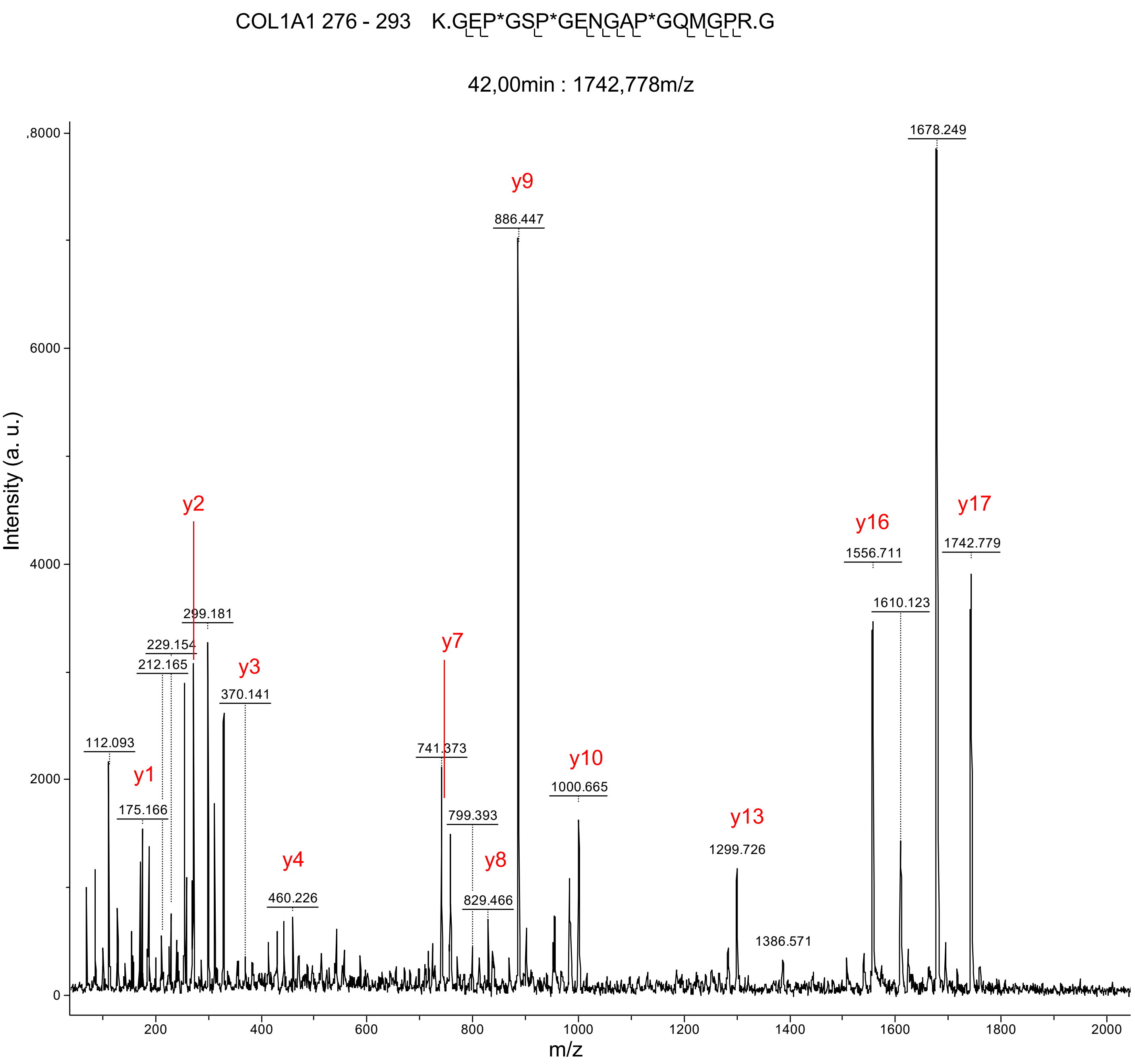

COL1A1 276 - 293 K.GEP*GSP*GENGAP*GQMGPR.G
42,00min : 1742,778m/z
Intensity (a. u.)
m/z
1678.249
y9
886.447
y2
y16
y17
1556.711
1742.779
1610.123
299.181
229.154
212.165
y3
370.141
y7
112.093
741.373
y1
y10
1000.665
175.166
799.393
y13
1299.726
y4
y8
460.226
829.466
1386.571

A (continued)

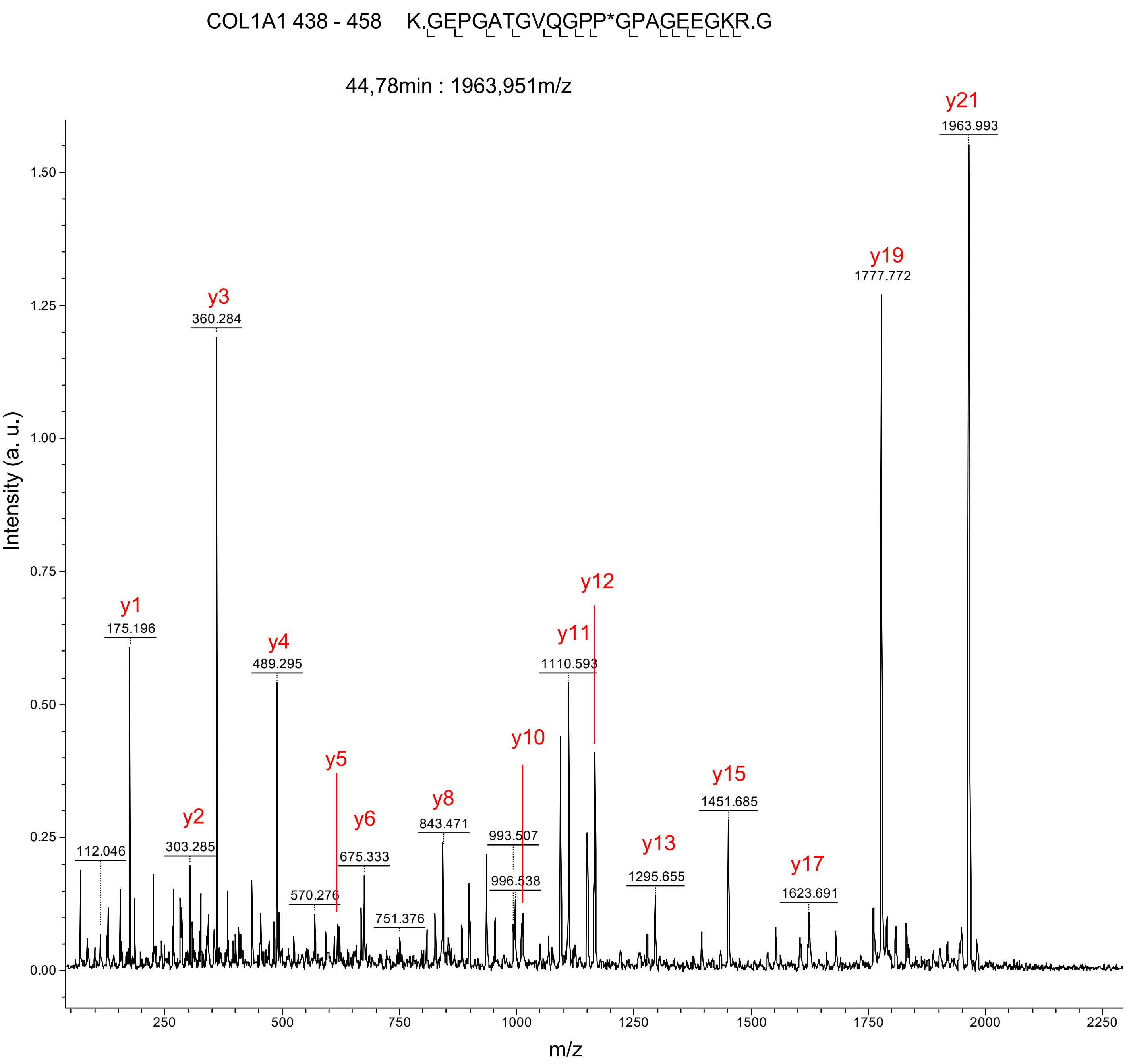

COL1A1 438 - 458 K.GEPGATGVQGPP*GPAGEEGKR.G
44,78min : 1963,951m/z
Intensity (a. u.)
m/z
y1
175.196
y2
303.285
y3
360.284
y4
489.295
y5
570.276
y6
675.333
751.376
y8
843.471
y10
993.507
996.538
y11
1110.593
y12
y13
1295.655
y15
1451.685
y17
1623.691
y19
1777.772
y21
1963.993
112.046

A (continued)

COL1A1 438 - 458    K.GEP*GATGVQGPP*GPAGEEGKR.G

43,55min : 1979,973m/z

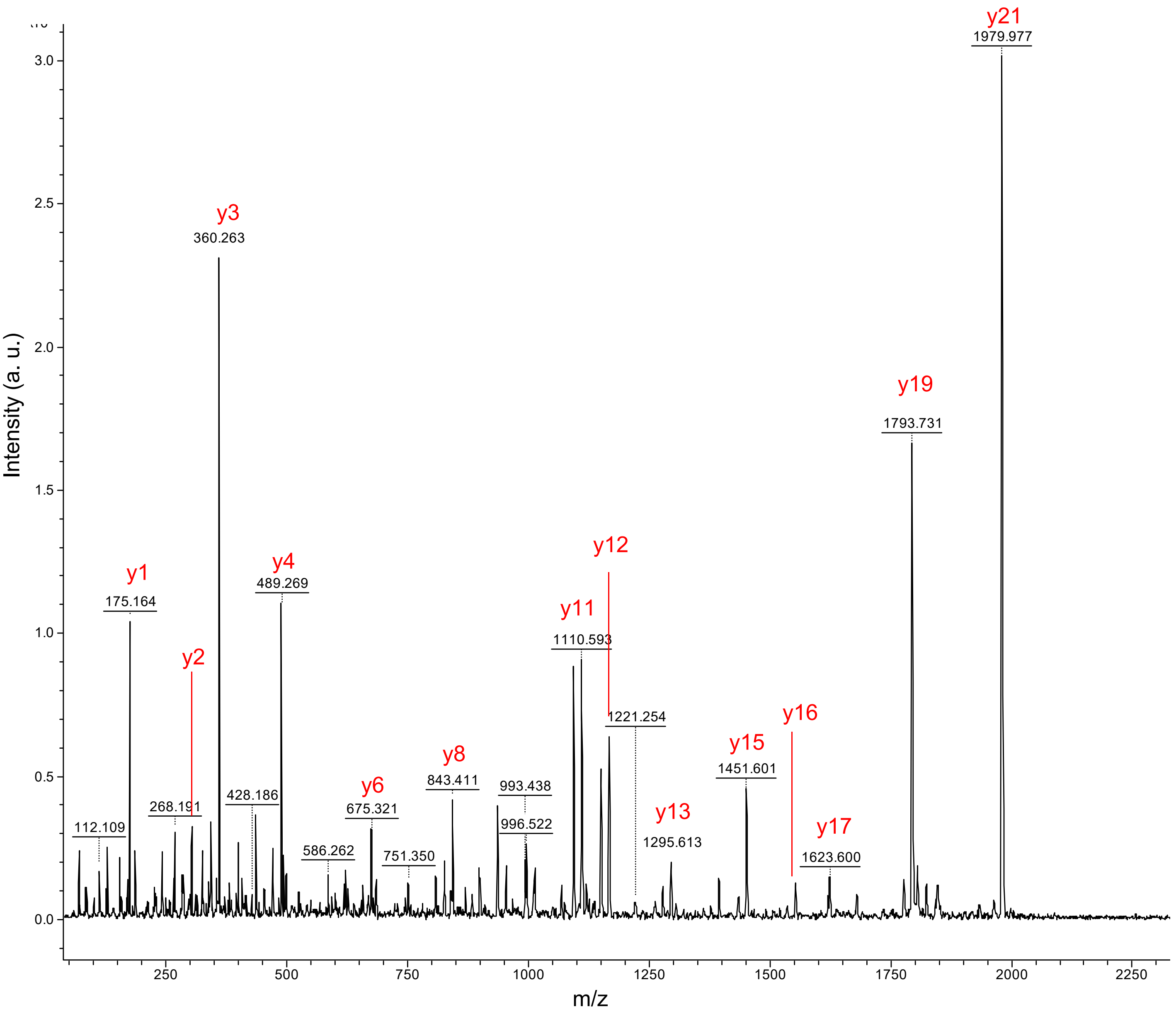

A (continued)

COL1A1 462 - 476    R.GEPGPSGLP*GPP*GER.G

47,95min : 1435,694m/z

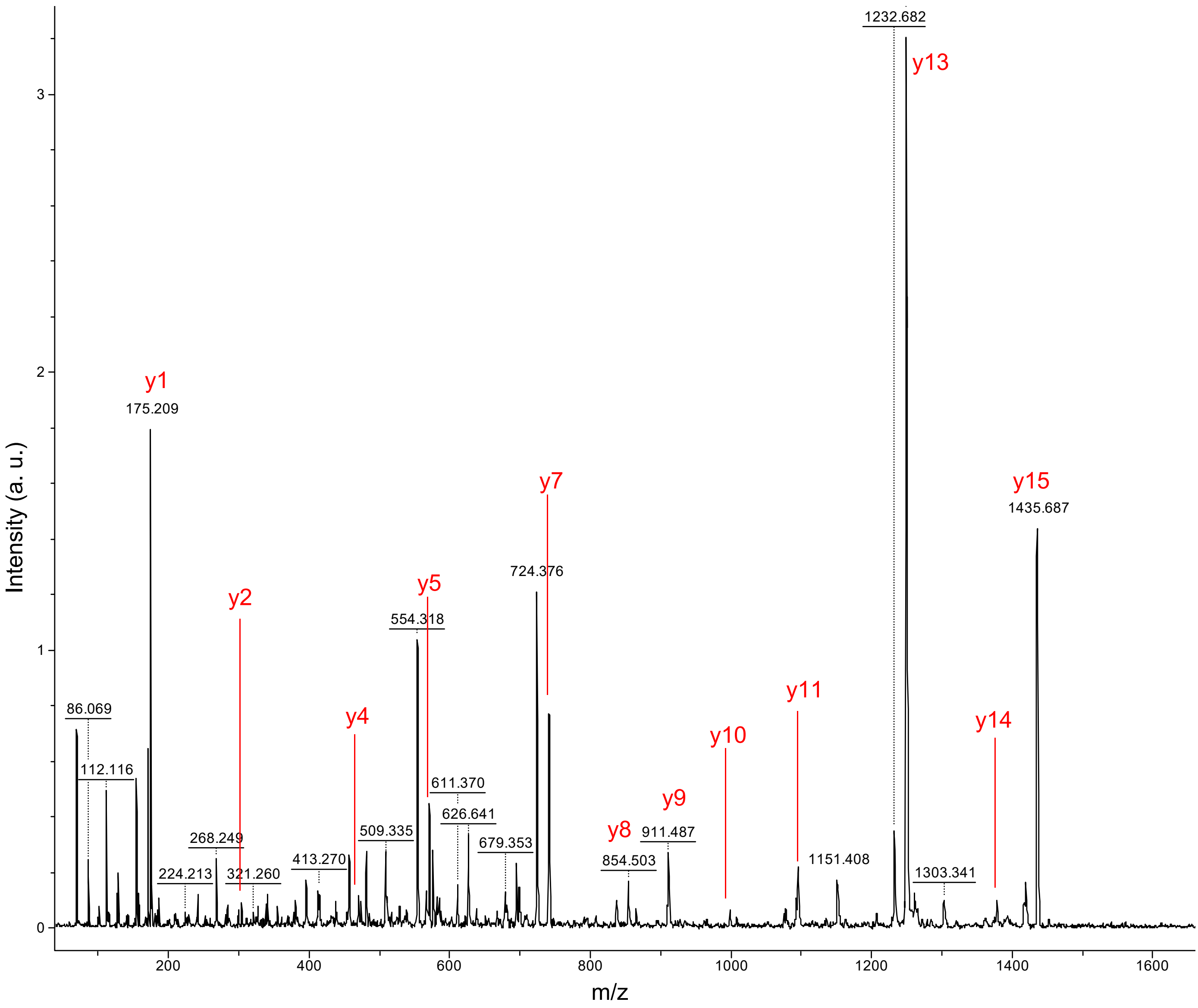

A (continued)

COL1A1 462 - 476    R.GEP*GPSGLP*GPP*GER.G

46,79min : 1451,681m/z

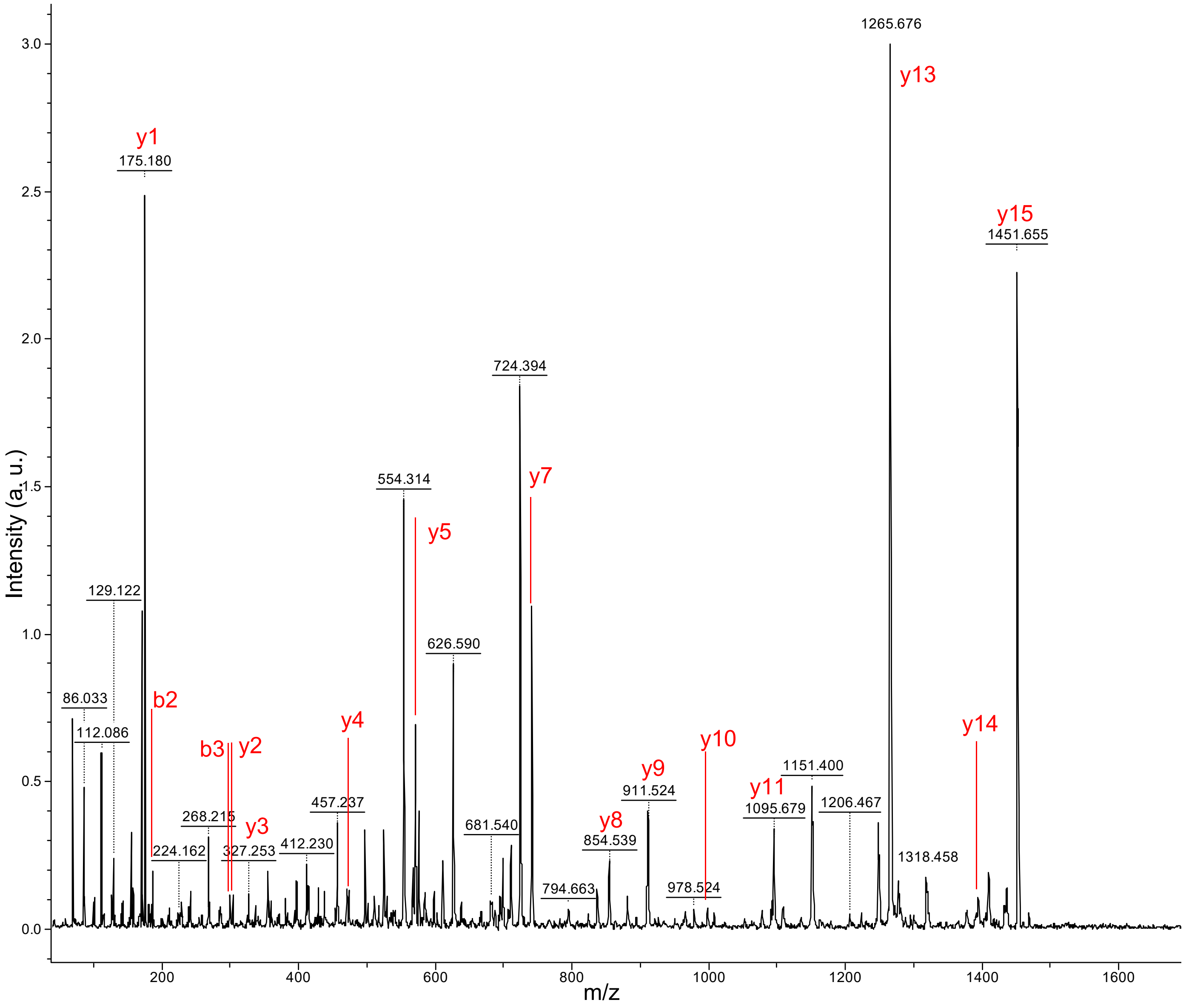

A (continued)

COL1A1 510 - 527    K.GSPGEAGRP*GEAGLP*GAK.G

44.30in : 1639.83m/z

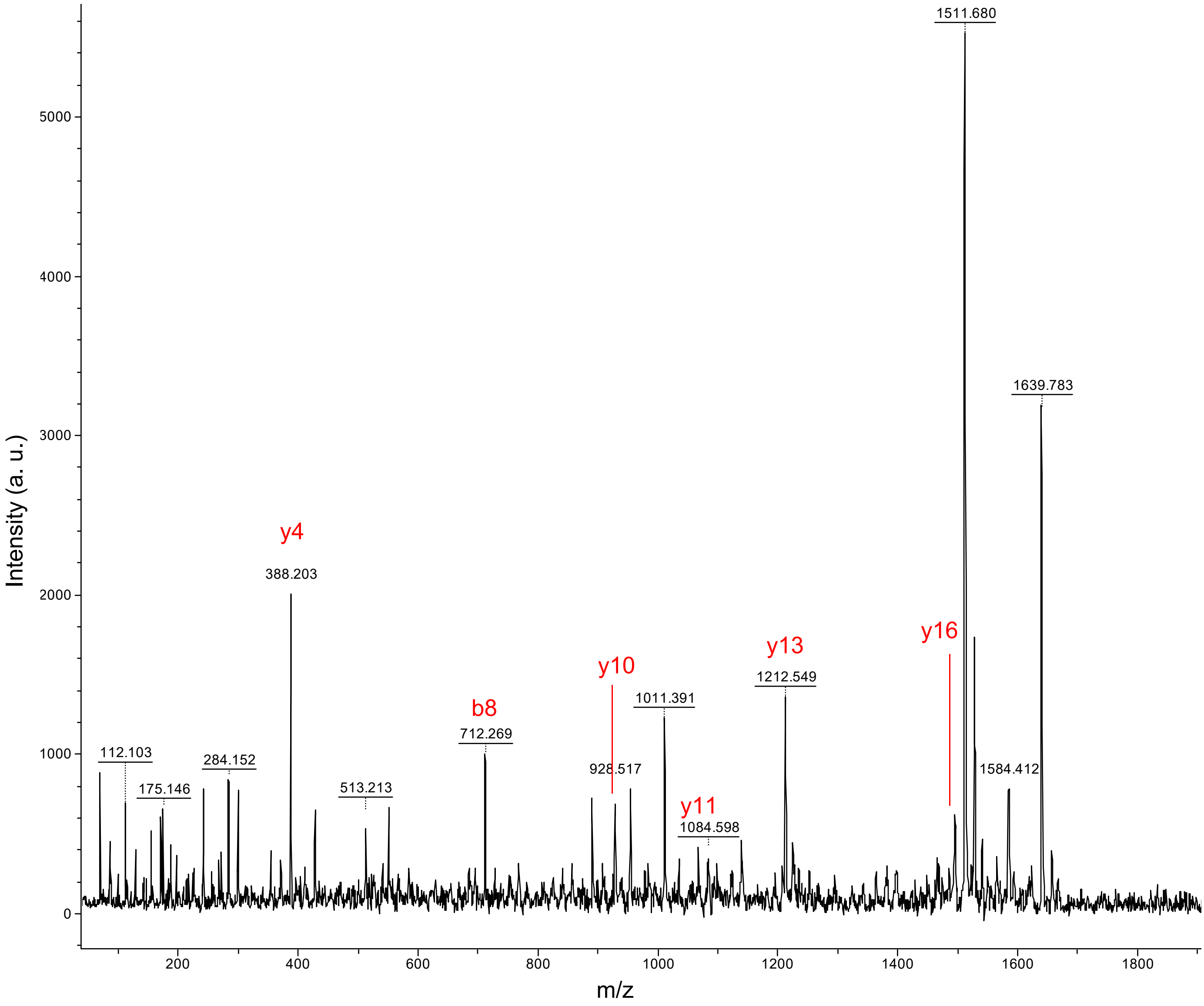

A (continued)

COL1A1 510 - 527    K.GSP*GEAGRP*GEAGLP*GAK.G

43.25min : 1655.83m/z

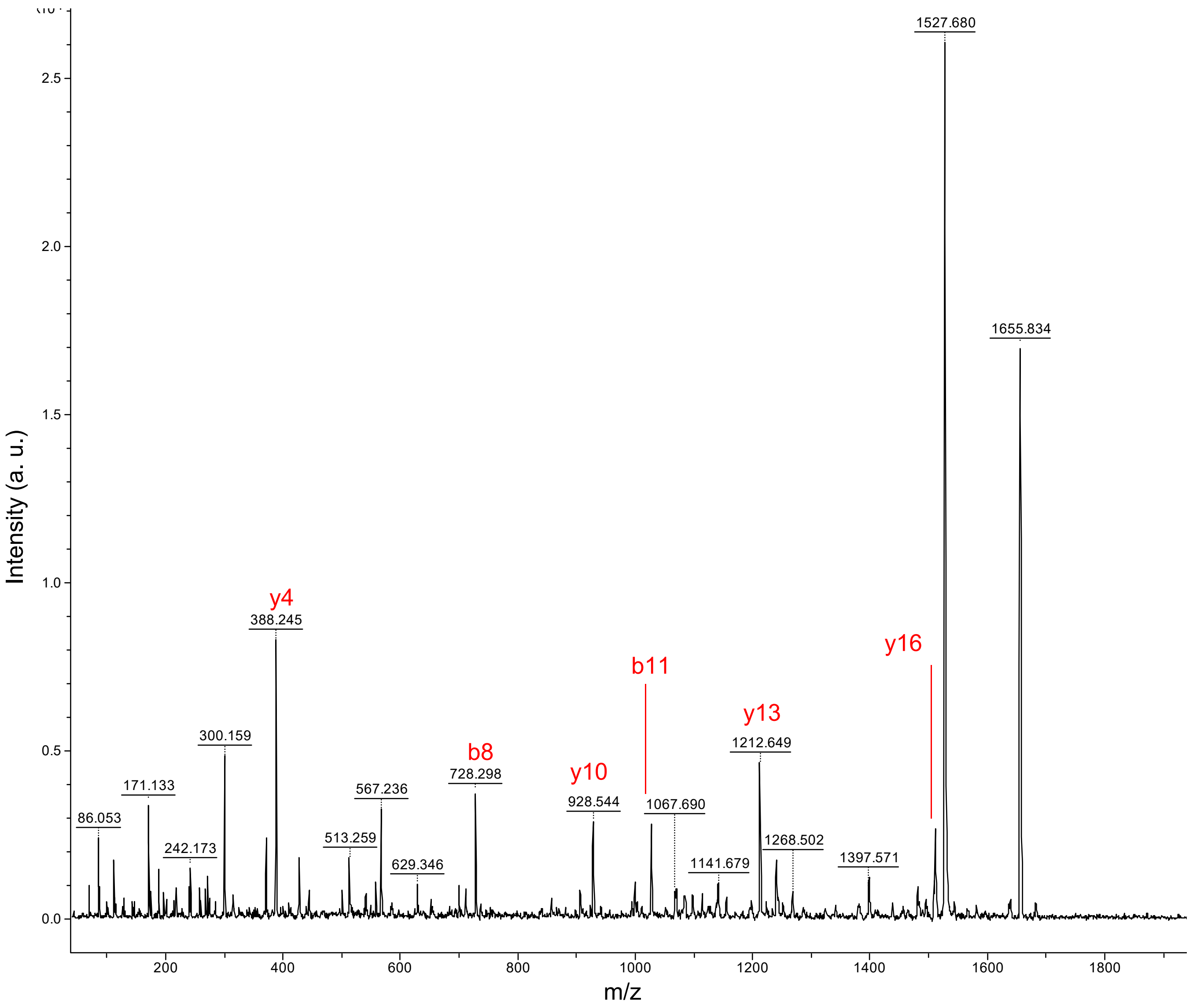

A (continued)

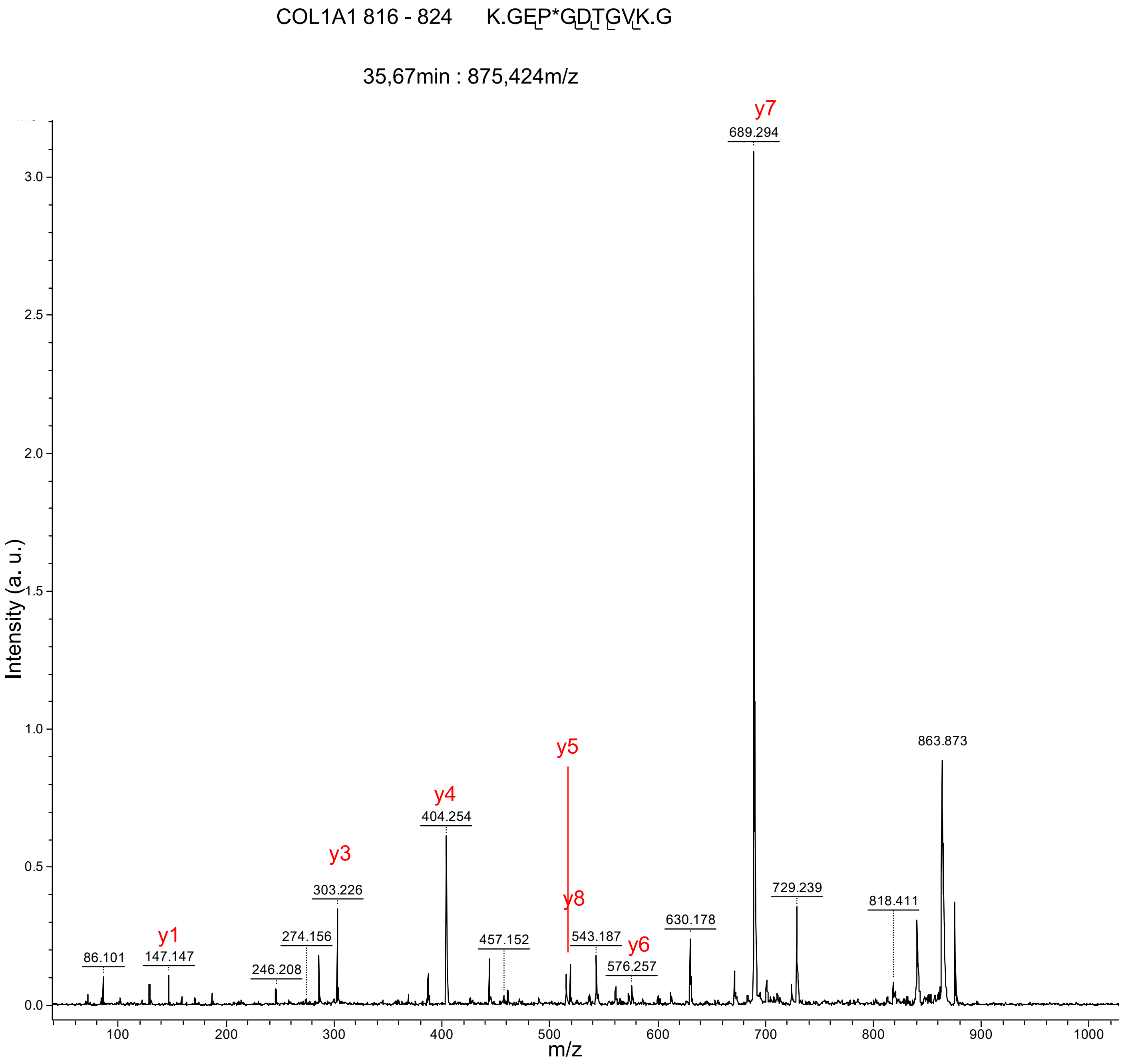

COL1A1 816 - 824 K.GEP*GDTGVK.G
35,67min : 875,424m/z
Intensity (a. u.)
m/z
y1
y3
y4
y5
y8
y6
y7
86.101
147.147
246.208
274.156
303.226
404.254
457.152
543.187
576.257
630.178
689.294
729.239
818.411
863.873

A (continued)

COL1A2 120 – 138 F.QGPAGEP*GEP*GQTGPAGPR.G

19.95min : 1791.869m/z

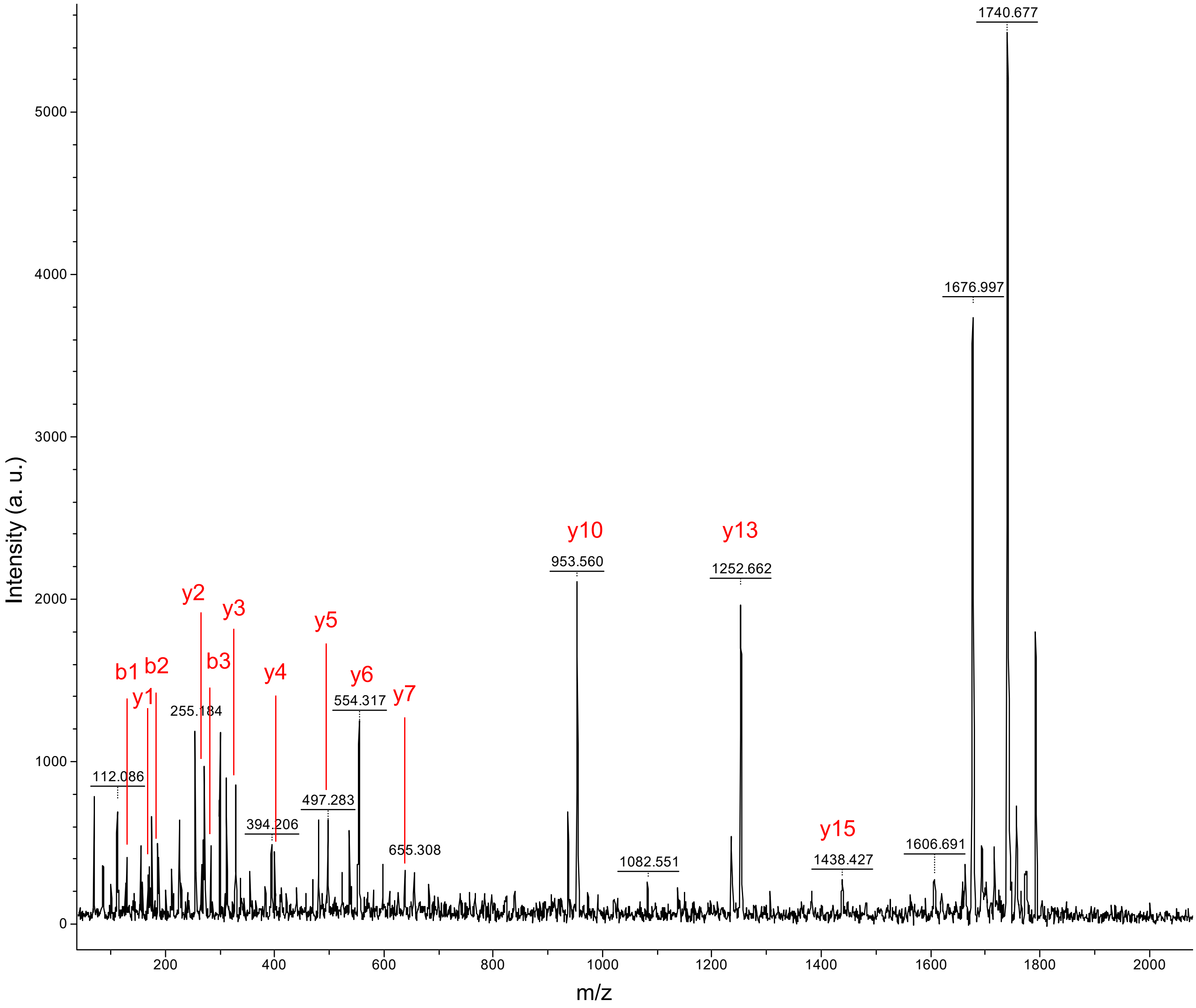

A (continued)

COL1A2 205 – 222 K.GEPGAP*GENGTP*GQAGAR.G

38,32min : 1654,762m/z

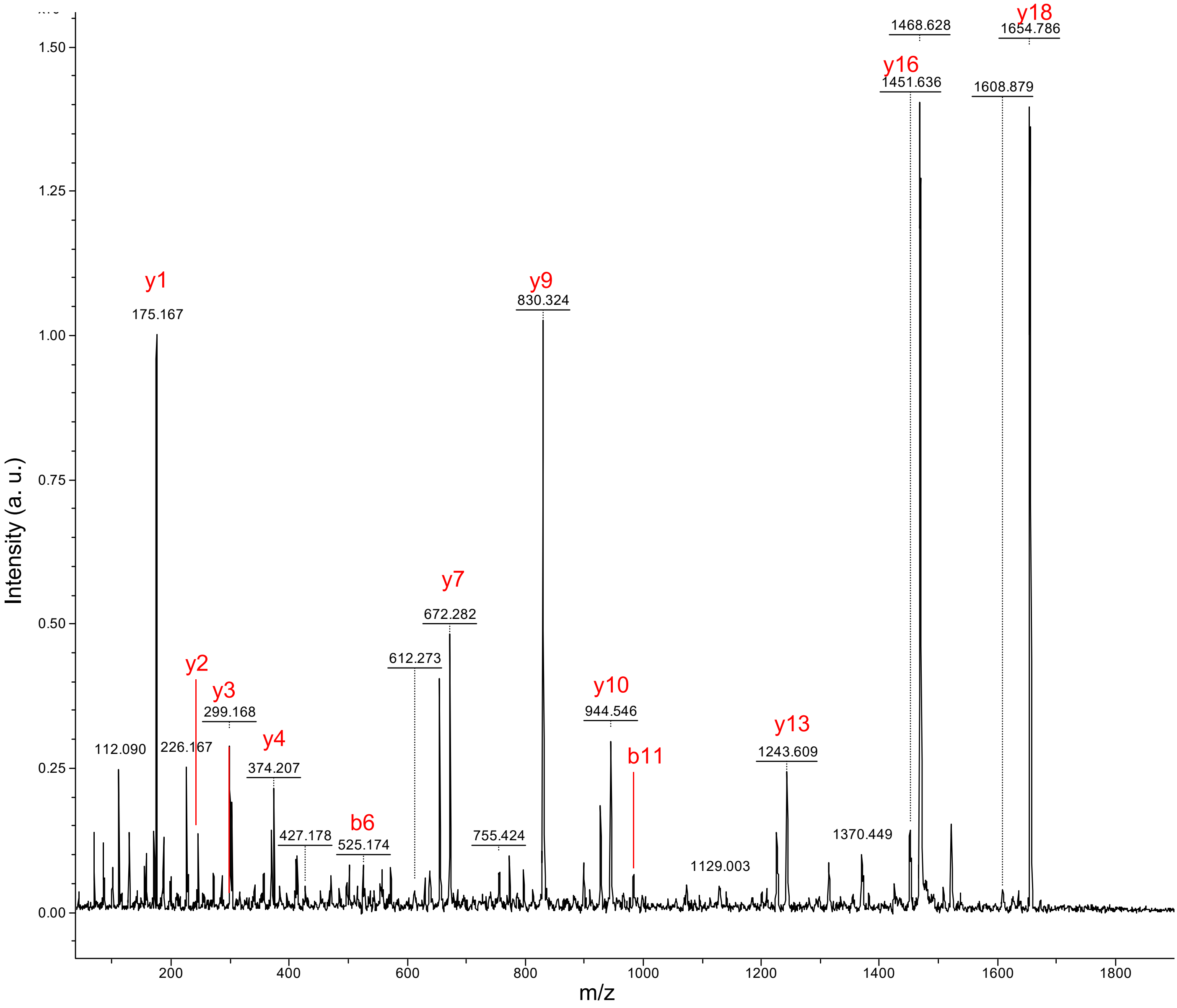

A (continued)

COL1A2 205 – 222 K.GEP*GAP*GENGTP*GQAGAR.G

36,66min : 1670,771m/z

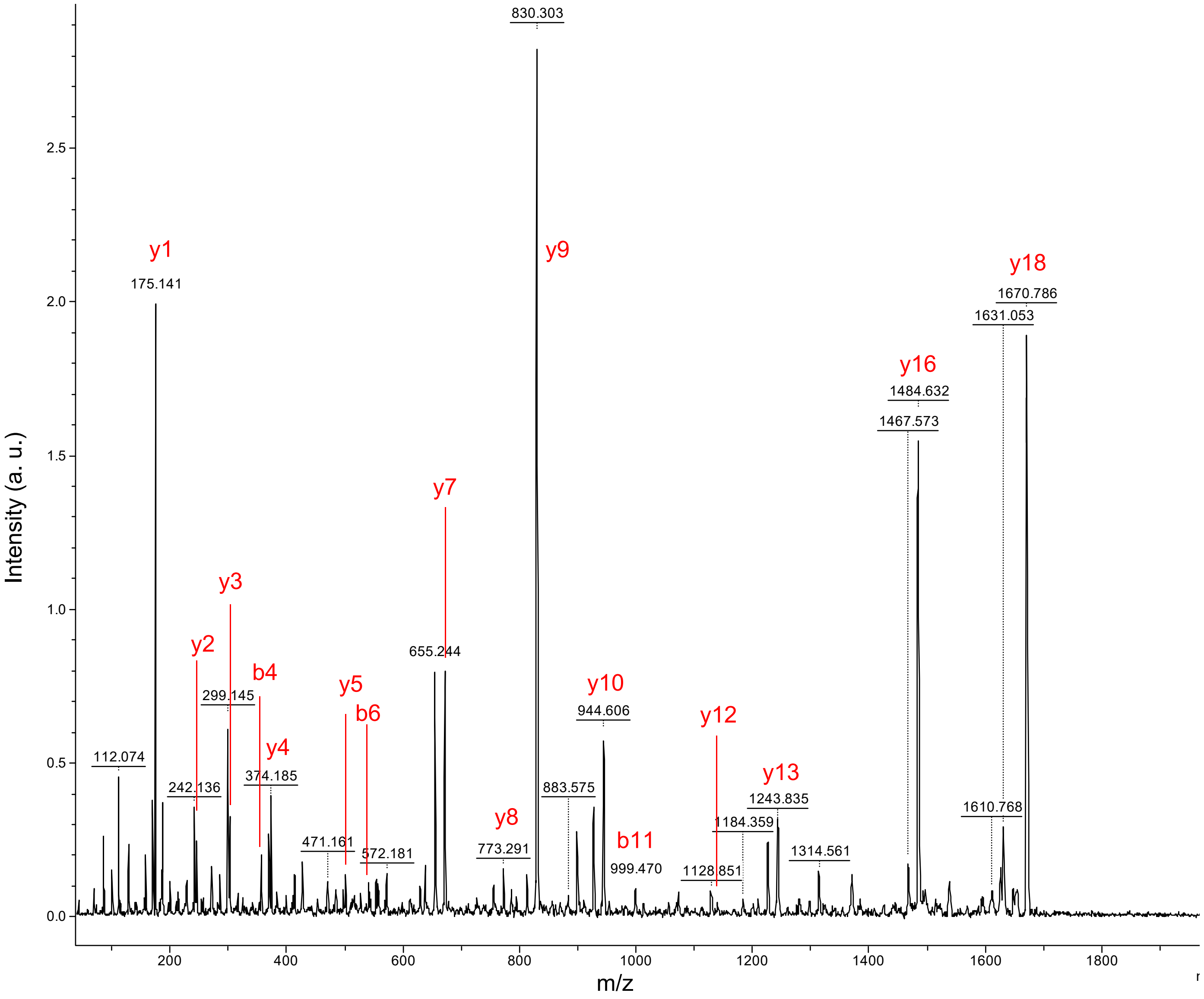

A (continued)

COL1A2 367 - 387 K.GEPGSVGAQGPP*GPSGEEGKR.G

43,03min : 1965,961m/z

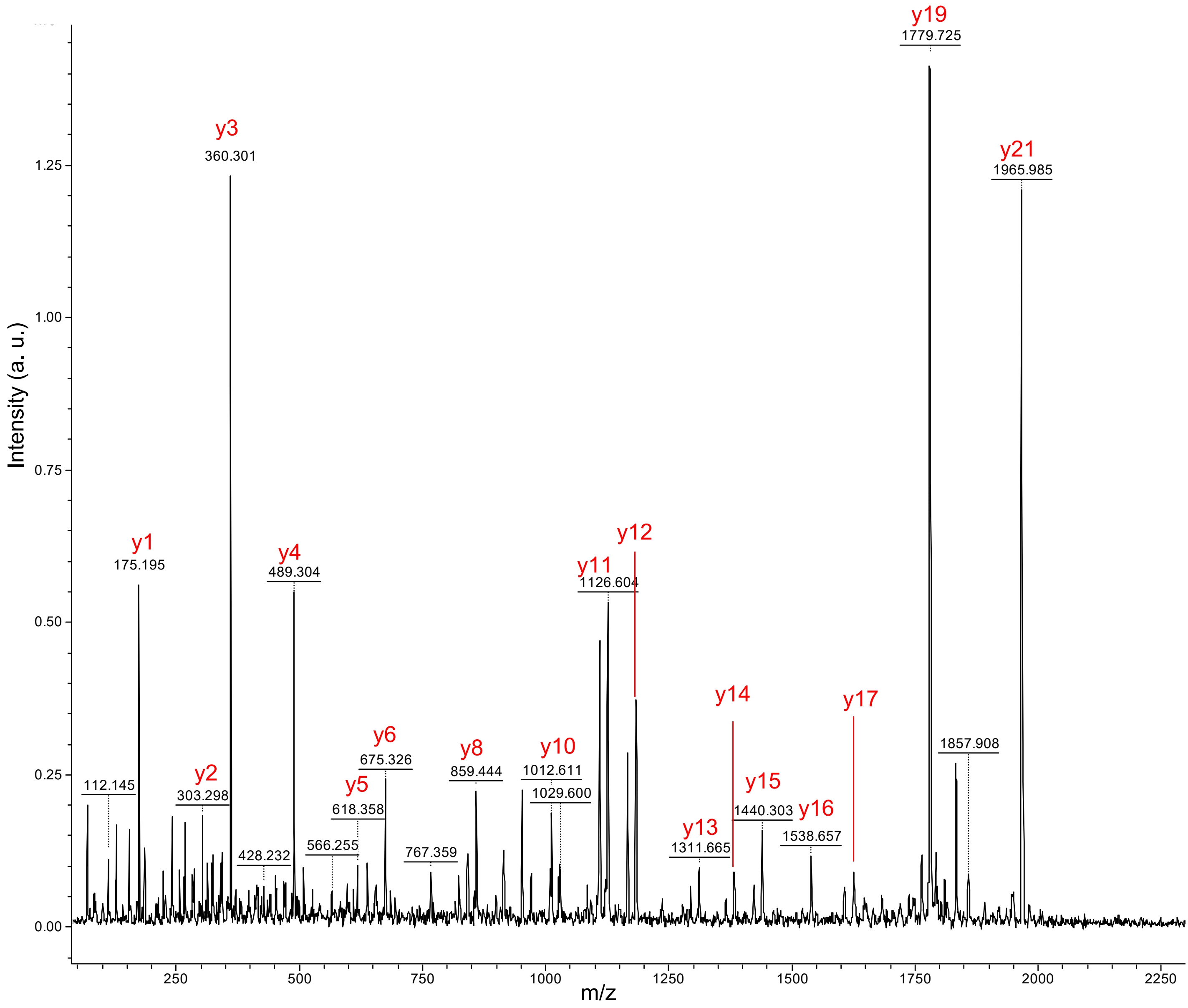

A (continued)

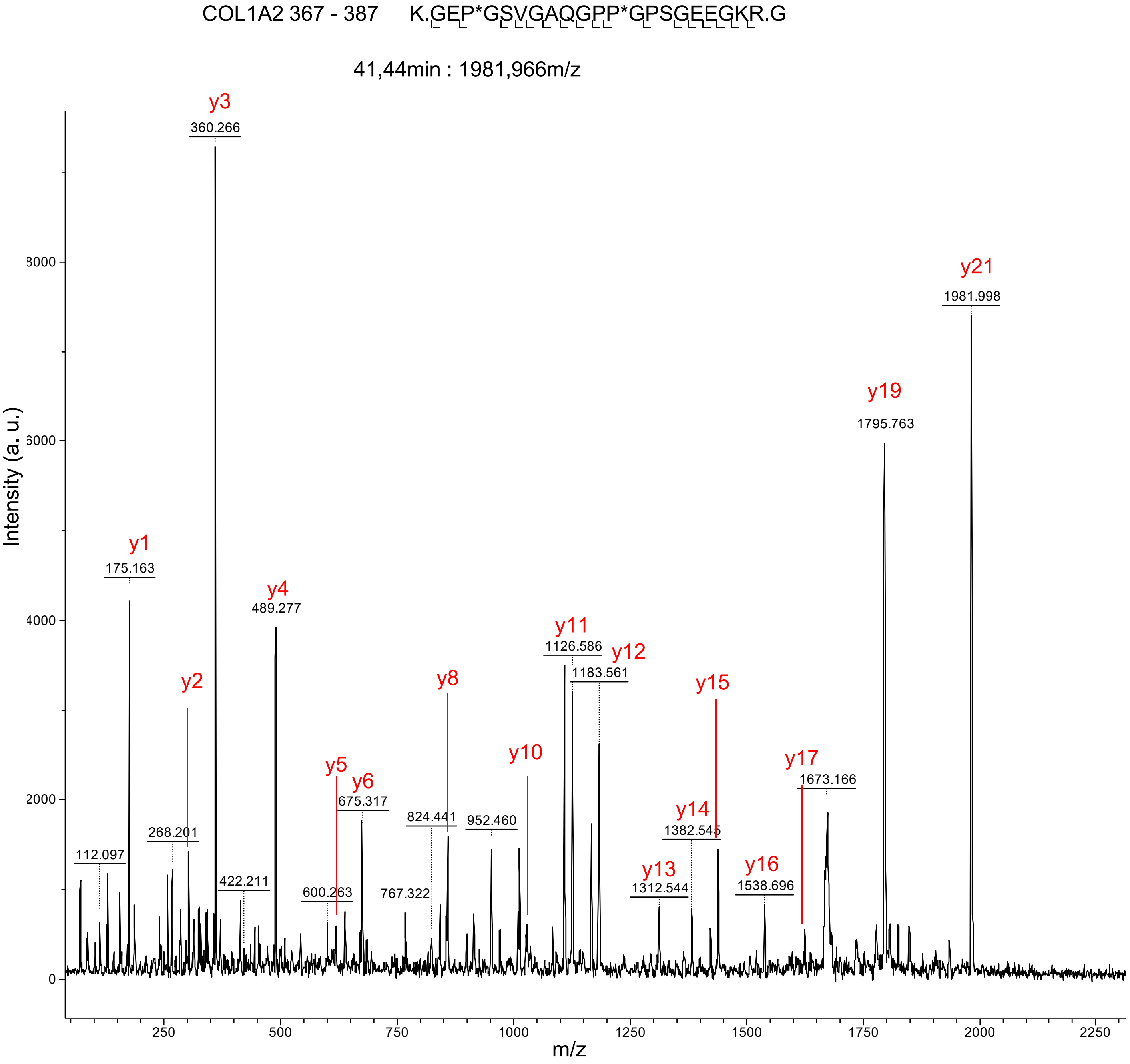

COL1A2 367 - 387 K.GEP*GSVGAQGPP*GPSGEEGKR.G
41,44min : 1981,966m/z
Intensity (a. u.)
m/z
y1
175.163
y2
268.201
y3
360.266
y4
489.277
y5
y6
675.317
y8
824.441
y10
y11
1126.586
y12
1183.561
y13
1312.544
y14
1382.545
y15
y16
1538.696
y17
1673.166
y19
1795.763
y21
1981.998
112.097
422.211
600.263
767.322
952.460

A (continued)

COL1A2 439 - 456 R.GPNGDAGRP*GEPGLMGPR.G

50,79min : 1750,844m/z

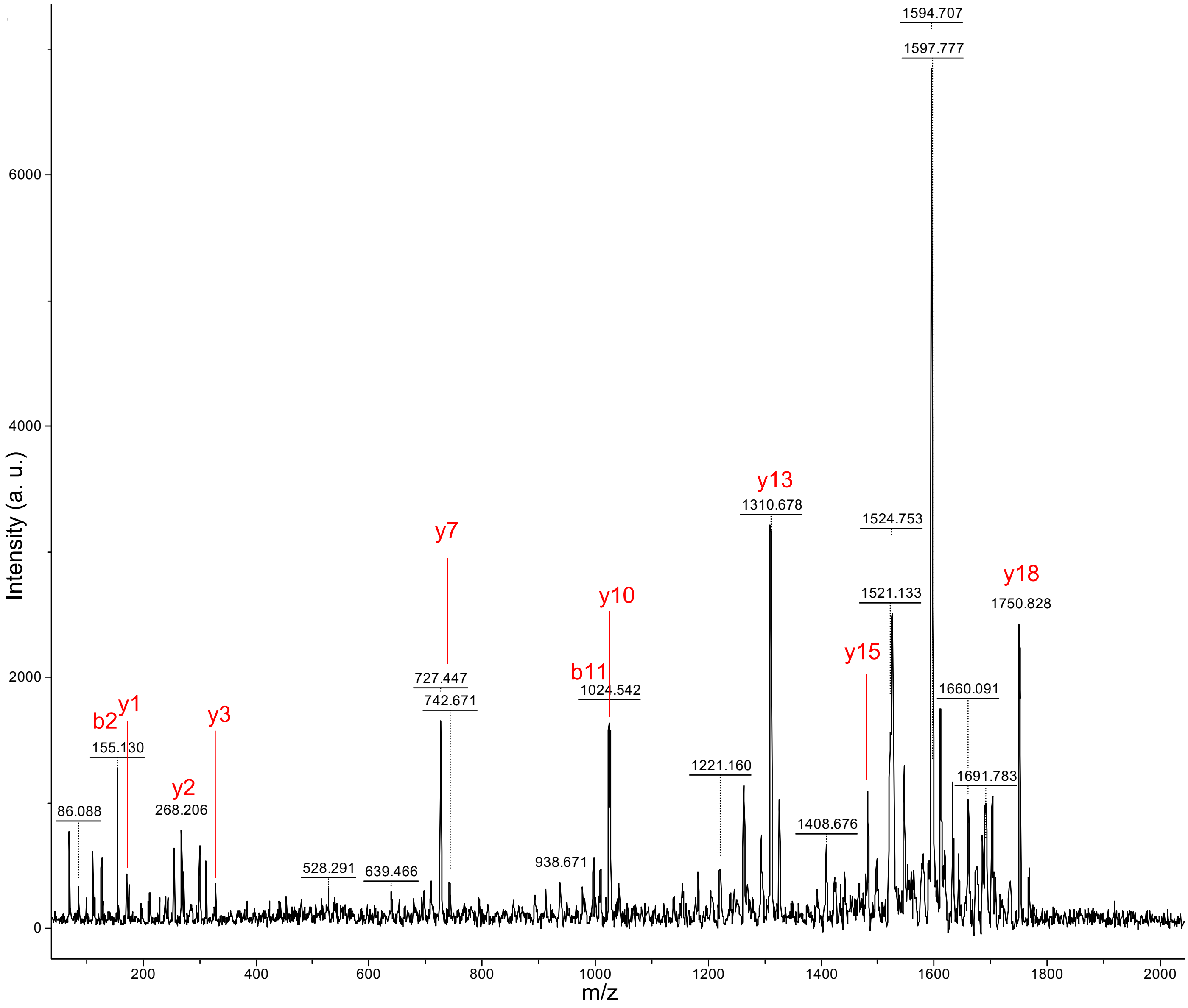

A (continued)

COL1A2 439 - 456    R.GPNGDAGRP*GEP*GLMGPR.G

50,20min : 1765,834m/z

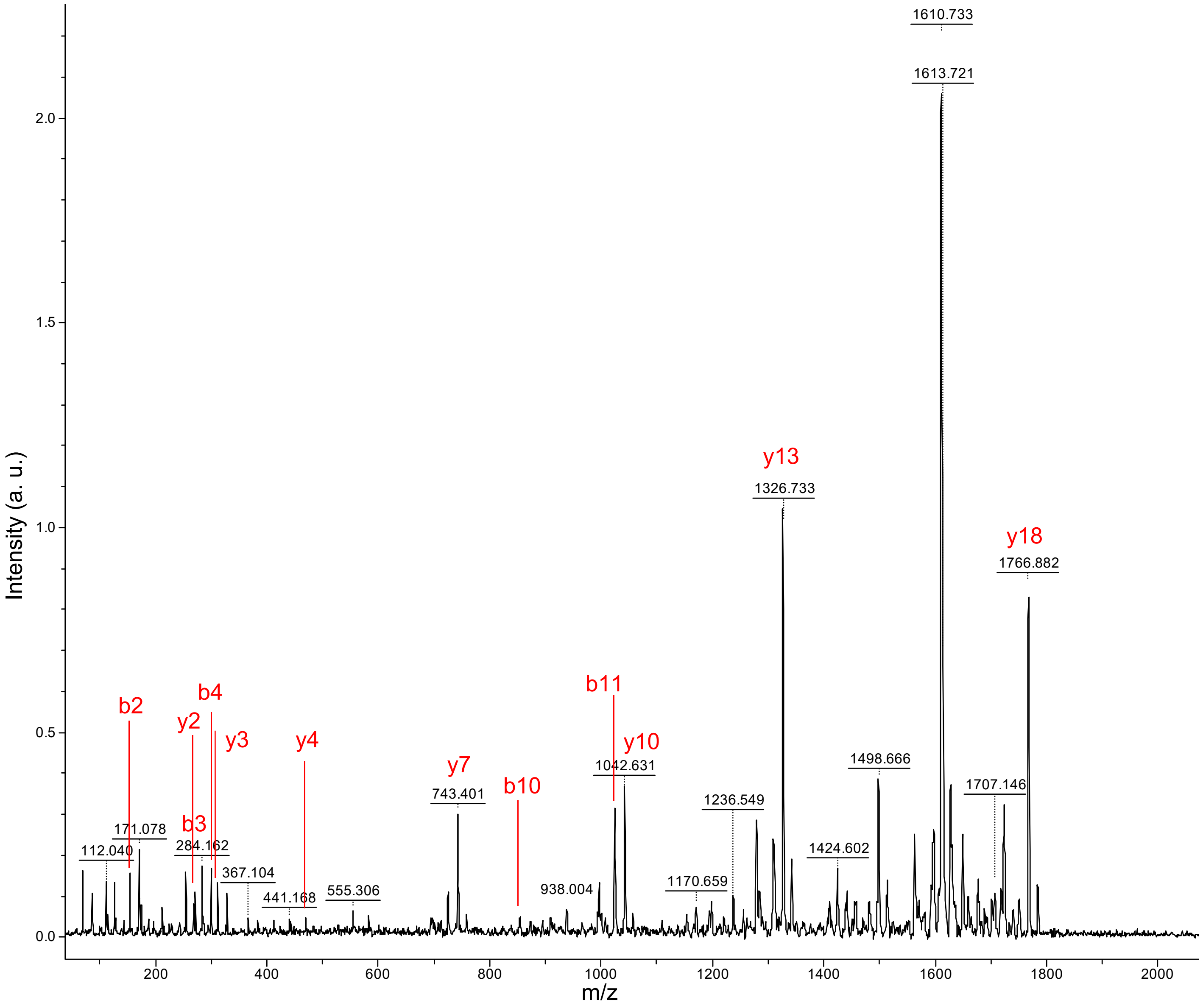

A (continued)

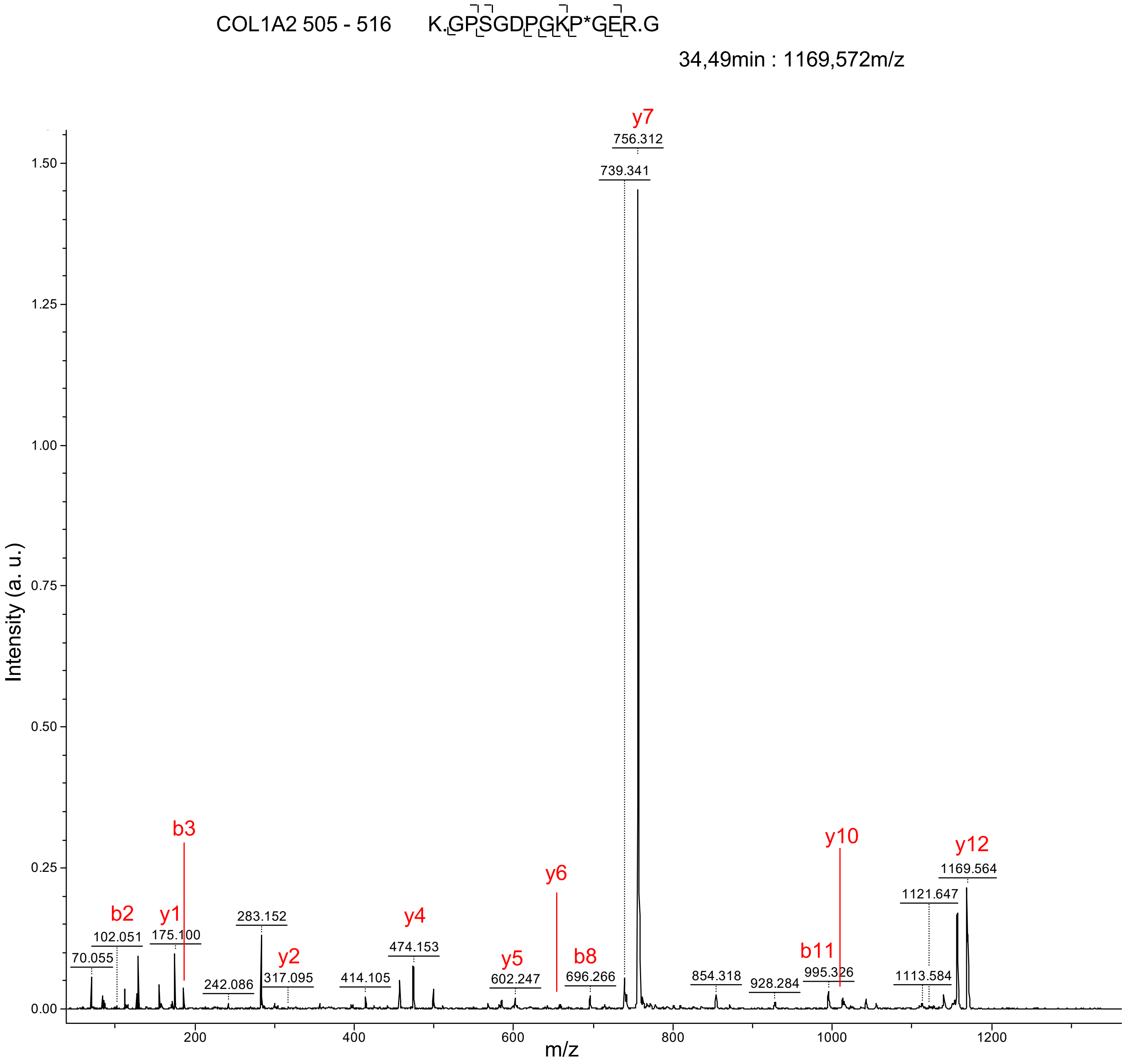
COL1A2 505 - 516
K.GPSGDPGKP*GER.G
34,49min : 1169,572m/z
Intensity (a. u.)
m/z
y7
756.312
739.341
b3
y10
y12
1169.564
1121.647
y6
b2
y1
102.051
175.100
283.152
y4
474.153
70.055
y2
317.095
y5
602.247
b8
696.266
b11
995.326
854.318
928.284
1113.584
242.086
414.105
1.50
1.25
1.00
0.75
0.50
0.25
0.00
200
400
600
800
1000
1200

A (continued)

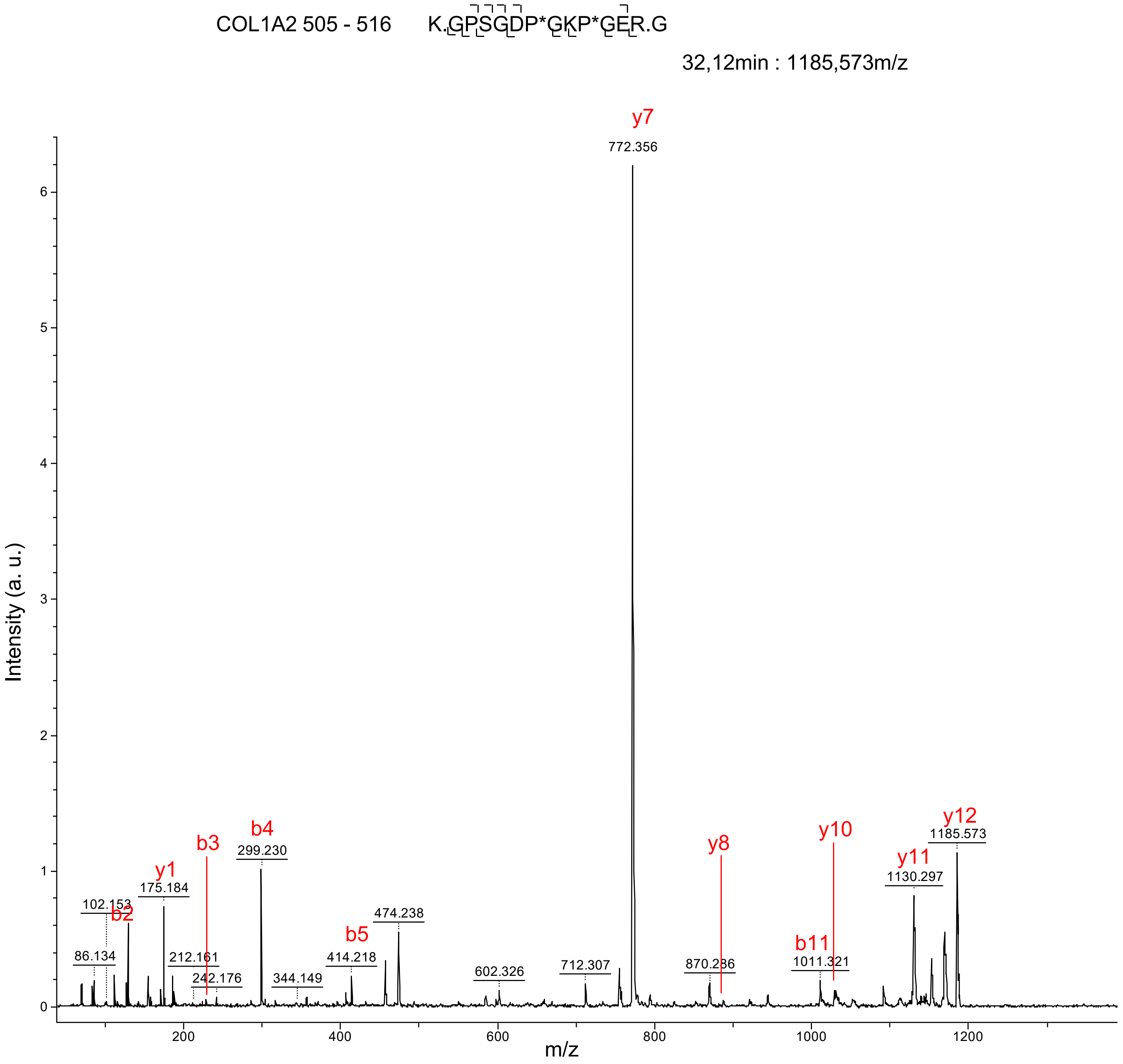

COL1A2 505 - 516
K.GPSGDP*GKP*GER.G
32,12min : 1185,573m/z
y7
772.356
Intensity (a. u.)
m/z
b2
102.153
86.134
y1
175.184
b3
212.161
242.176
b4
299.230
344.149
b5
414.218
474.238
602.326
712.307
y8
870.286
b11
1011.321
y10
y11
1130.297
y12
1185.573

A (continued)

COL1A2 853 - 885 K.GPSGEPGTAGAP*GTAGP*QGLLGAPGILGLP*GSR.G

68,04min : 2915,460m/z

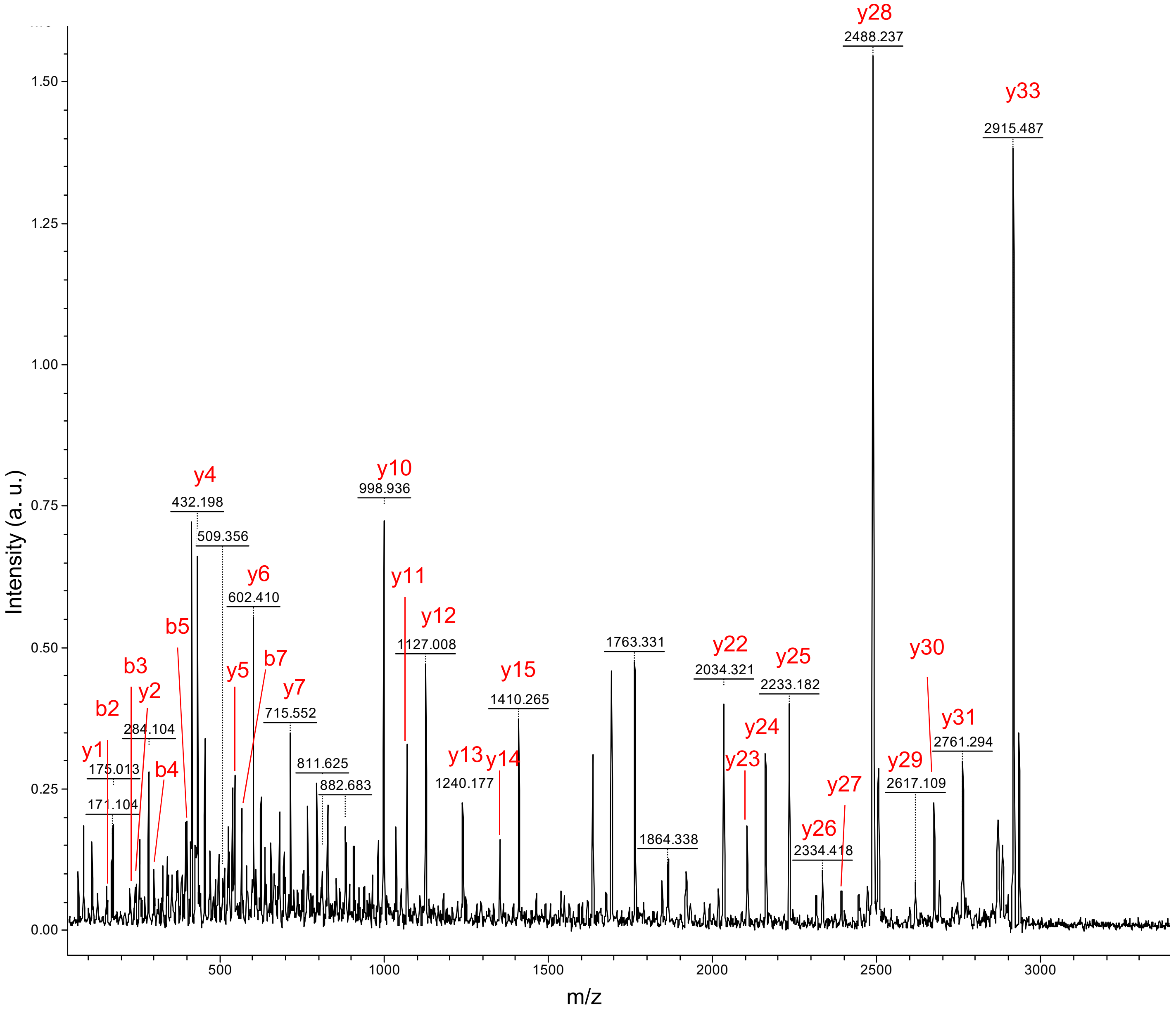

A (continued)

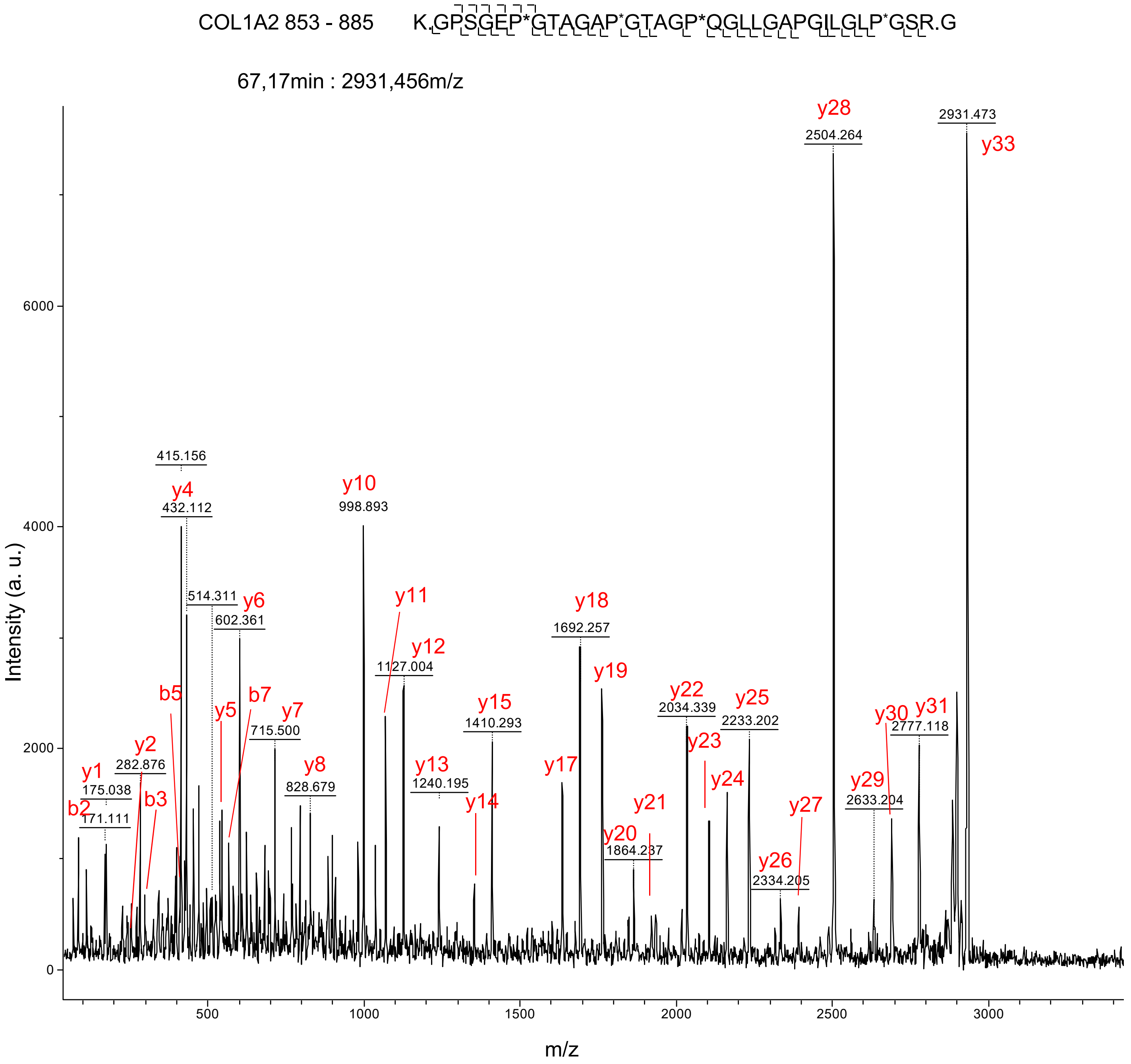

COL1A2 853 - 885
K.GPSGEP*GTAGAP*GTAGP*QGLLGAPGILGLP*GSR.G
67,17min : 2931,456m/z
Intensity (a. u.)
m/z
y28
2504.264
2931.473
y33
415.156
y4
432.112
y10
998.893
514.311
y6
602.361
y11
y12
1127.004
y18
1692.257
y19
b5
y5
b7
y7
715.500
y15
1410.293
y22
2034.339
y25
2233.202
y30
y31
2777.118
y23
y2
282.876
y1
175.038
b3
b2
171.111
y8
828.679
y13
1240.195
y14
y17
y24
y21
y29
2633.204
y27
y20
1864.237
y26
2334.205

A (continued)

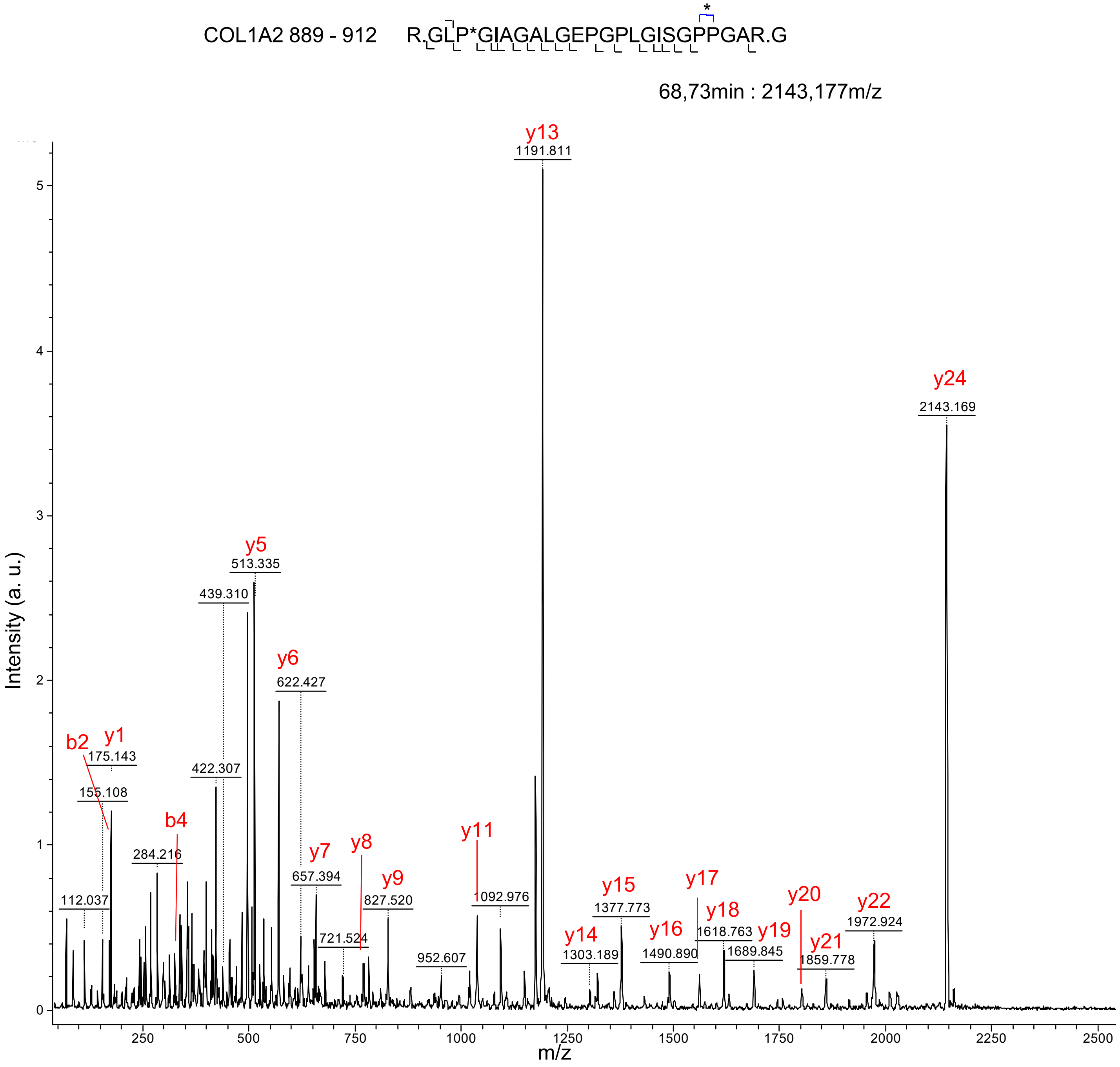

COL1A2 889 - 912
R.GLP*GIAGALGEPGPLGISGPPGAR.G
*
68,73min : 2143,177m/z
Intensity (a. u.)
m/z
y13
1191.811
y24
2143.169
y5
513.335
439.310
y6
622.427
b2
y1
175.143
155.108
422.307
b4
284.216
112.037
y7
657.394
y8
721.524
y9
827.520
952.607
y11
1092.976
y14
1303.189
y15
1377.773
y16
1490.890
y17
y18
1618.763
y19
1689.845
y20
y21
1859.778
y22
1972.924

A (continued)

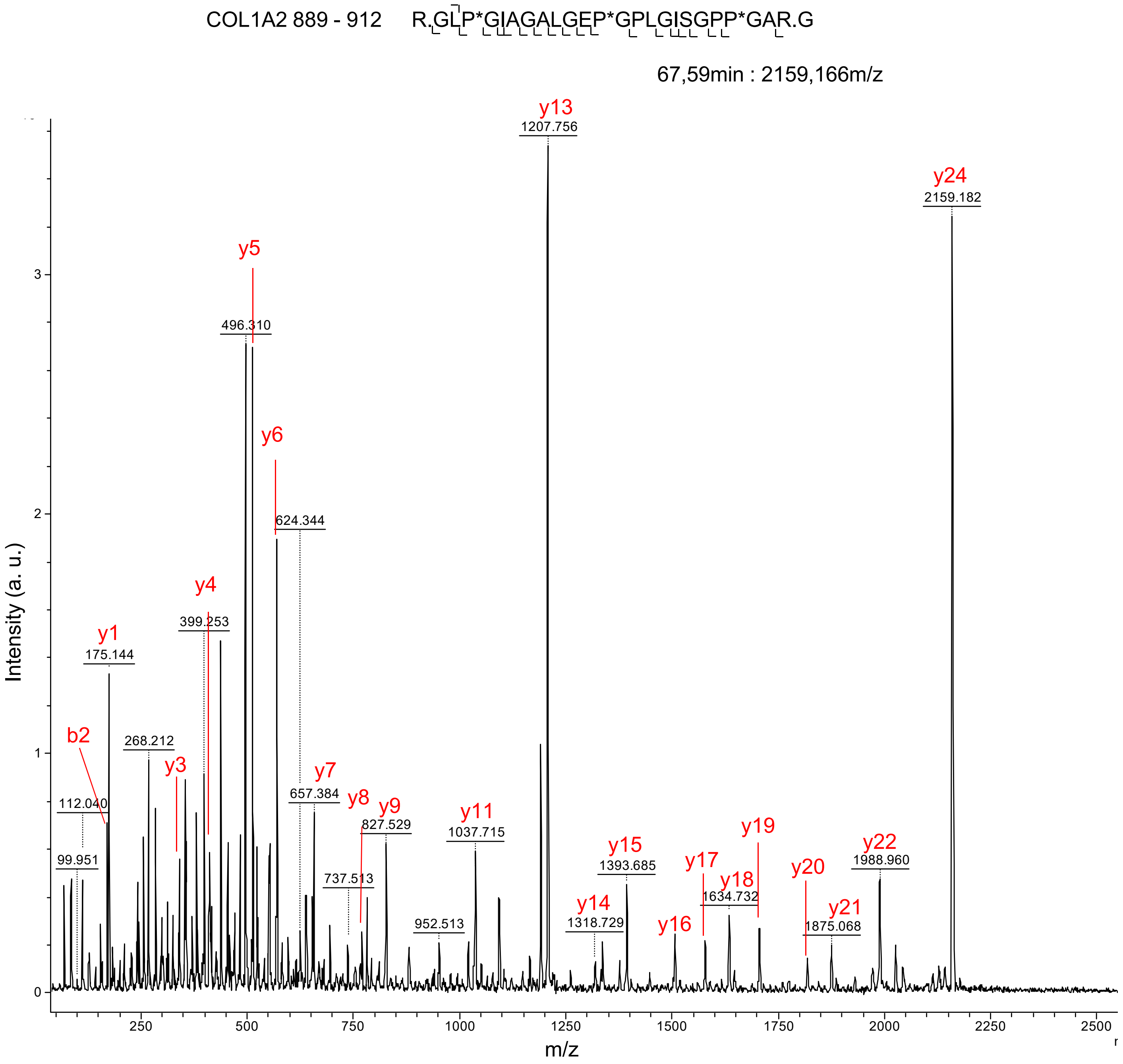

COL1A2 889 - 912
R.GLP*GIAGALGEP*GPLGISGPP*GAR.G
67,59min : 2159,166m/z
Intensity (a. u.)
m/z
y1
y3
y4
y5
y6
y7
y8
y9
y11
y13
y14
y15
y16
y17
y18
y19
y20
y21
y22
y24
b2
99.951
112.040
175.144
268.212
399.253
496.310
624.344
657.384
737.513
827.529
952.513
1037.715
1207.756
1318.729
1393.685
1634.732
1875.068
1988.960
2159.182

A (continued)

COL1A2 985 - 1,002 R.GEP*GPAGSVGPVGAVGPR.G

53,31min : 1576,811m/z

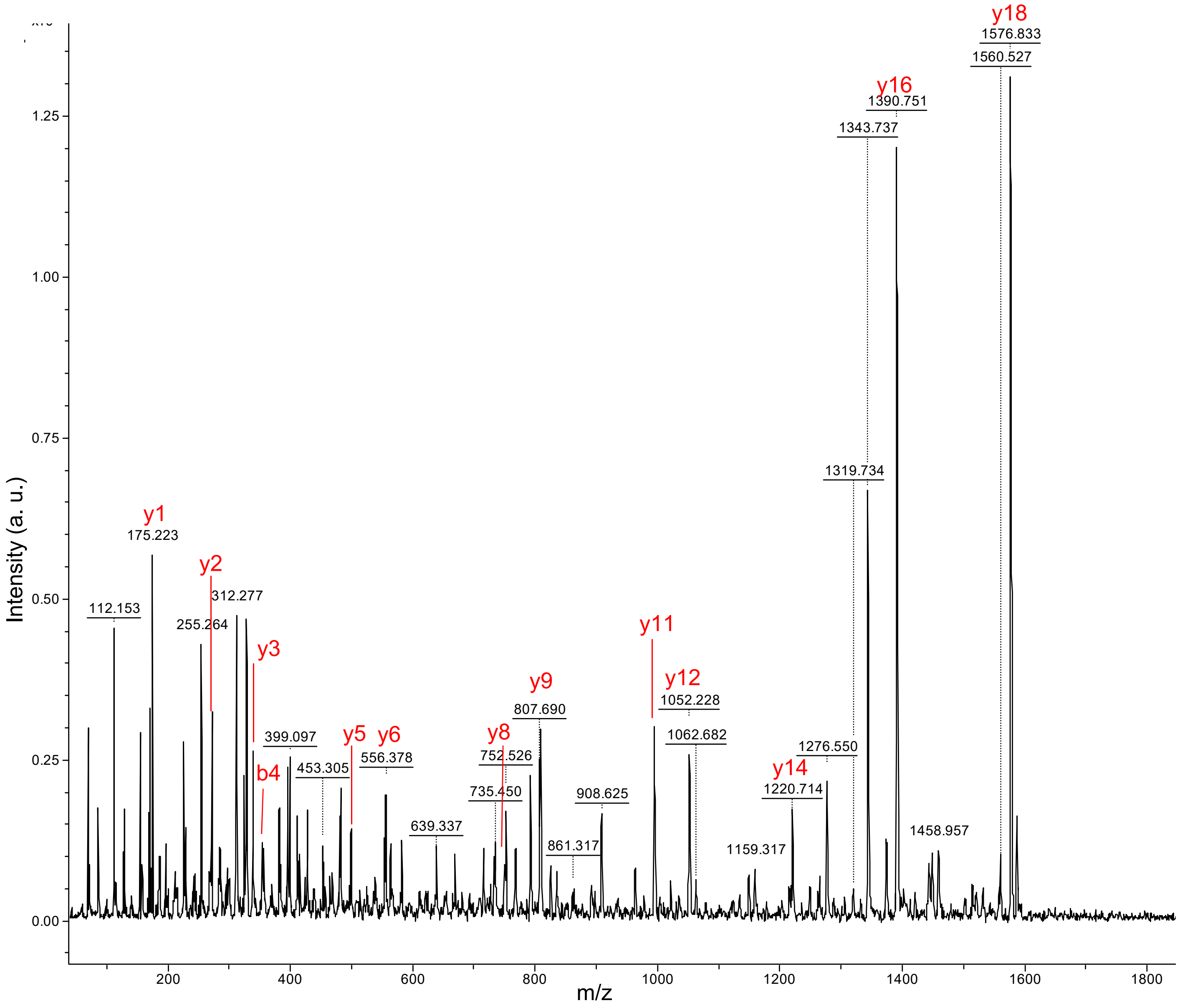

B

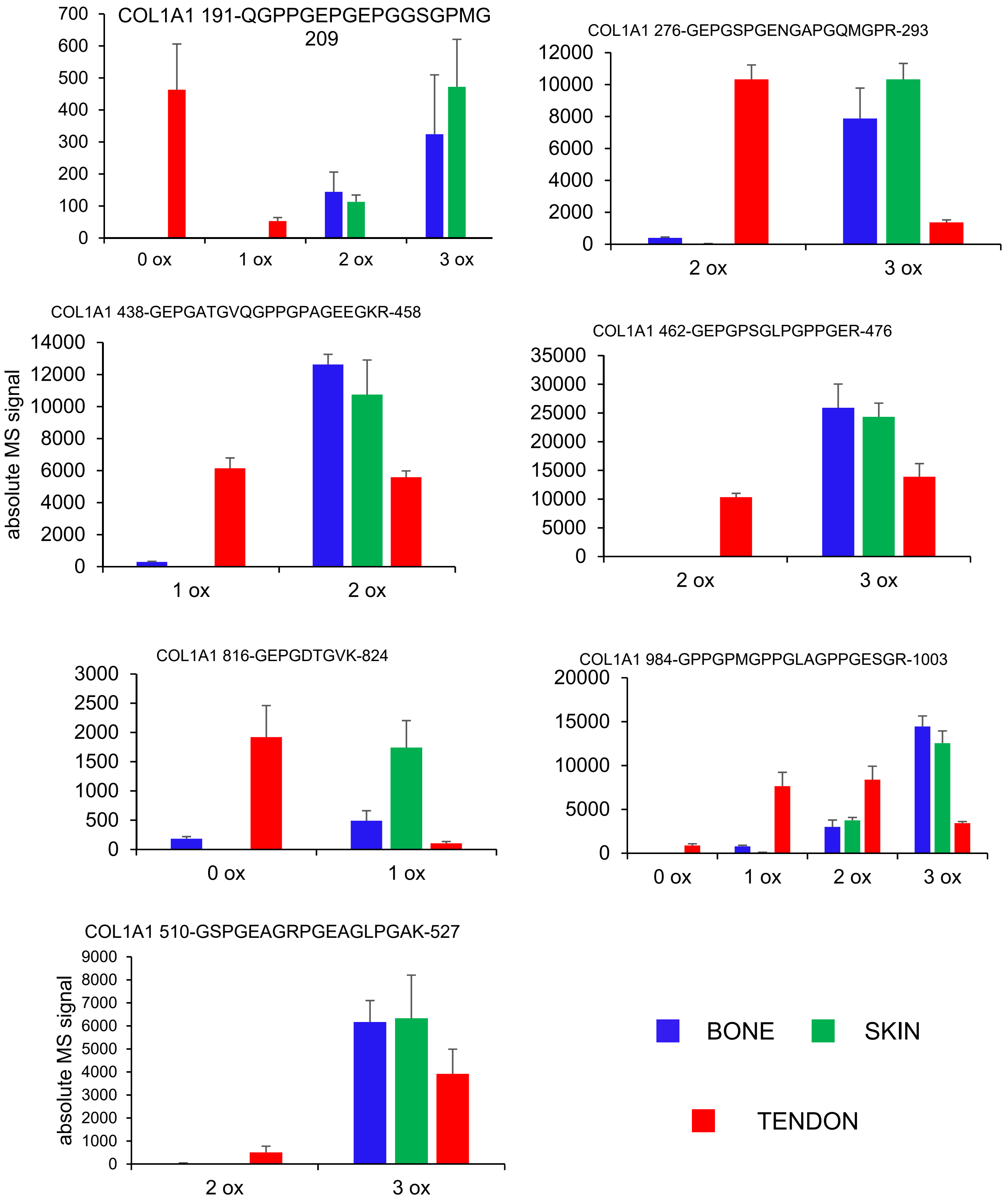

COL1A1 191-QGPPGEPGEPGGSGPMG 209
700
600
500
400
300
200
100
0
0 ox
1 ox
2 ox
3 ox
COL1A1 276-GEPGSPGENGAPGQMGPR-293
12000
10000
8000
6000
4000
2000
0
2 ox
3 ox
COL1A1 438-GEPGATGVQGPPGPAGEEGKR-458
absolute MS signal
14000
12000
10000
8000
6000
4000
2000
0
1 ox
2 ox
COL1A1 462-GEPGPSGLPGPPGER-476
35000
30000
25000
20000
15000
10000
5000
0
2 ox
3 ox
COL1A1 816-GEPGDTGVK-824
3000
2500
2000
1500
1000
500
0
0 ox
1 ox
COL1A1 984-GPPGPMGPPGLAGPPGESGR-1003
20000
15000
10000
5000
0
0 ox
1 ox
2 ox
3 ox
COL1A1 510-GSPGEAGRPGEAGLPGAK-527
absolute MS signal
9000
8000
7000
6000
5000
4000
3000
2000
1000
0
2 ox
3 ox
BONE
SKIN
TENDON

B (continued)

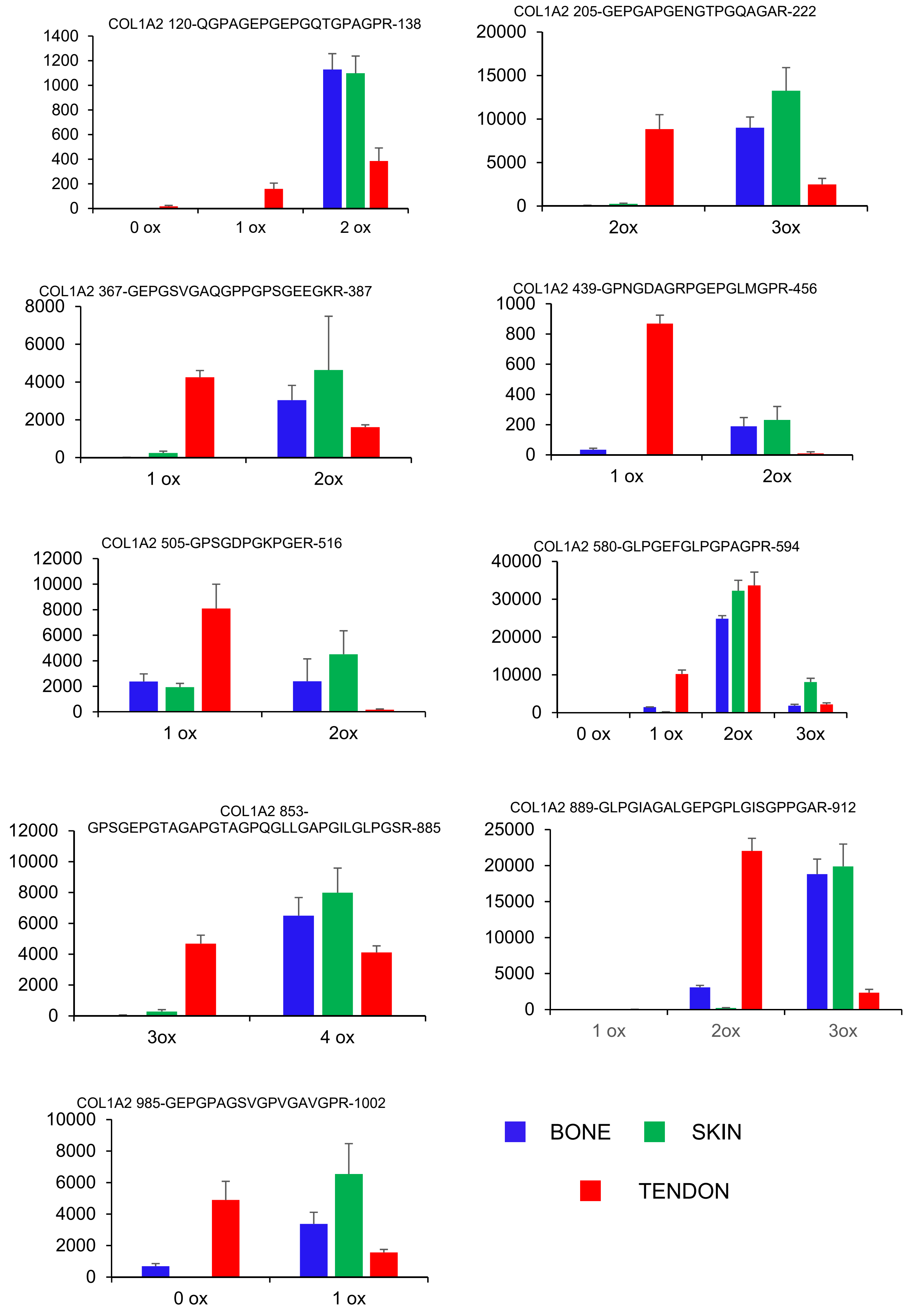
COL1A2 120-QGPAGEPGEPGQTGPAGPR-138
0
200
400
600
800
1000
1200
1400
0 ox
1 ox
2 ox
COL1A2 205-GEPGAPGENGTPGQAGAR-222
0
5000
10000
15000
20000
2ox
3ox
COL1A2 367-GEPGSVGAQGPPGPSGEEGKR-387
0
2000
4000
6000
8000
1 ox
2ox
COL1A2 439-GPNGDAGRPGEPGLMGPR-456
0
200
400
600
800
1000
1 ox
2ox
COL1A2 505-GPSGDPGKPGER-516
0
2000
4000
6000
8000
10000
12000
1 ox
2ox
COL1A2 580-GLPGEFGLPGPAGPR-594
0
10000
20000
30000
40000
0 ox
1 ox
2ox
3ox
COL1A2 853-GPSGEPGTAGAPGTAGPQGLLGAPGILGLPGSR-885
0
2000
4000
6000
8000
10000
12000
3ox
4 ox
COL1A2 889-GLPGIAGALGEPGPLGISGPPGAR-912
0
5000
10000
15000
20000
25000
1 ox
2ox
3ox
COL1A2 985-GEPGPAGSVGPVGAVGPR-1002
0
2000
4000
6000
8000
10000
0 ox
1 ox
BONE
SKIN
TENDON
