## Supplemental data S2 for "Tissue-specific collagen hydroxylation at GEP/GDP triplets mediated by P4HA2"

**Supplemental data S2. Detailed identification and relative quantitative data for peptides found underhydroxylated in chicken tendon.** (A) MS/MS spectra allowing determination of the residue positions of hydroxylations from chicken collagen. Crude MS/MS spectra of interest were mannually annotated, based on automated annotations obtained from the proteinscape server following Swissprot interrogation, in order to clarify the determinability of each proposed position. The locations of b and y ions are indicated on the spectra in red. The corresponding breaks are shown at the top and bottom of the corresponding peptide sequence, respectively. * indicates the positions of hydroxylations in the peptide sequence. (B) Detailed relative quantification data for peptides found under-hydroxylated in tendon collagen. Each panel shows the data obtained for a given peptide, the sequence and boundaries of which are given at the top. Graphs report on the absolute MS signal obtained for all identified versions of a given peptide. Statistics are not reported here due to their complexity but they are summarized in panel C. (C) Table summarizing positions and fold regulations of hydroxylation. Peptides are classified per their amino acid boundaries, referring to their positions in the $\alpha$1 (COL1A1) or $\alpha$2 chain (COL1A2) of type I chicken collagen precursor. Those found modulated in mouse but not in chicken are shown for comparison. The amino acid sequence is given, with numbers indicating the order of preferential hydroxylation according to MS/MS fragmentation of the distinct molecular states, $^{U}$ indicates that the exact residue position of hydroxylation could not be distinguished between similarly annotated residues. $^{R}$ indicates positions at which hydroxylation is found lower in tendon collagen. Underlined: sequences within which hydroxylated positions were determined from a chymotrypsin plus trypsin digest. GEP/GDP triplets are highlighted in red. Blue background: peptides found regulated only in the chicken; the corresponding peptides in mouse all lack a GEP; grey background: peptides found regulated in mouse but lacking a GEP in chicken. The average hydroxylation number of each peptide in skin and tendon, respectively, is calculated from total ion chromatograms. ND: no data available; NI: quantitative data non-interpretable. * the state with 3 hydroxylations was not considered for calculations. t-test p-value is given for tendon versus skin (n=3 per tissue).

A

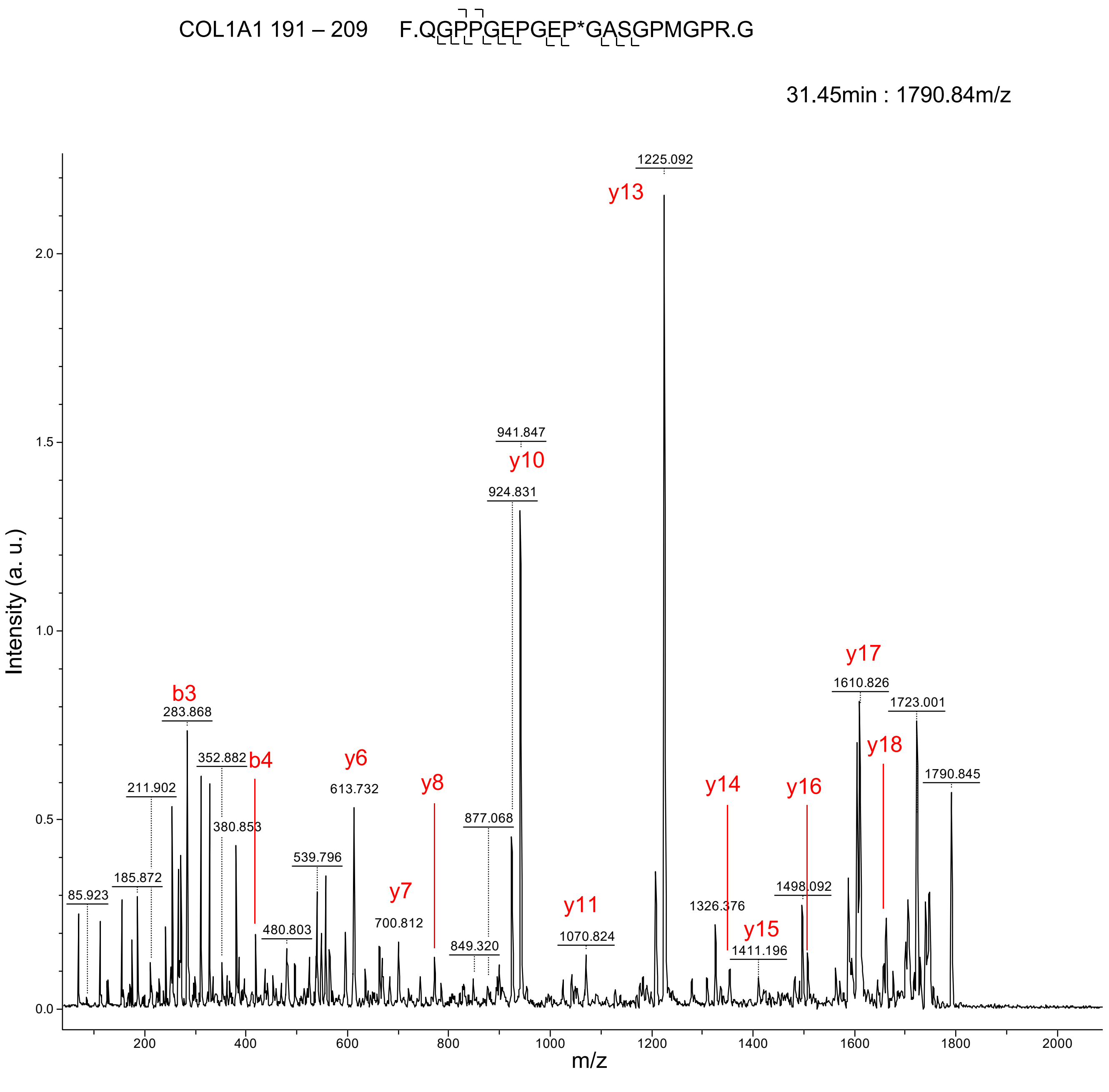
COL1A1 191 – 209
F.QGPPGEPGEP*GASGPMGPR.G
31.45min : 1790.84m/z
Intensity (a. u.)
m/z
1225.092
y13
941.847
y10
924.831
877.068
y17
1610.826
1723.001
y18
1790.845
b3
283.868
352.882
b4
y6
613.732
211.902
380.853
539.796
185.872
85.923
480.803
y7
700.812
y8
849.320
y11
1070.824
y14
1326.376
y15
1411.196
y16
1498.092

A (continued)

COL1A1 191 – 209 F.QGPP*GEPGEP*GASGPMGPR.G

30.10min : 1806.81m/z

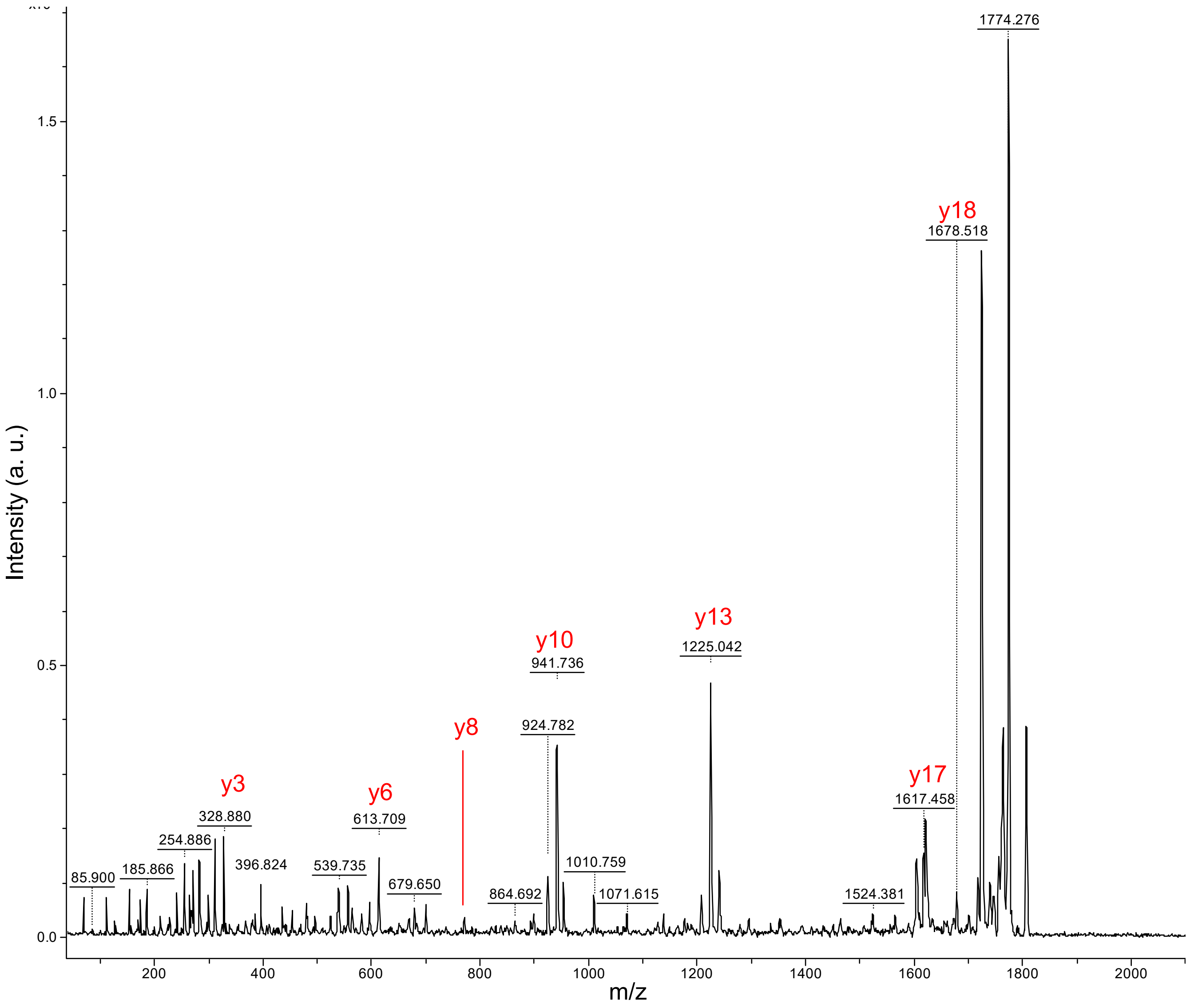

A (continued)

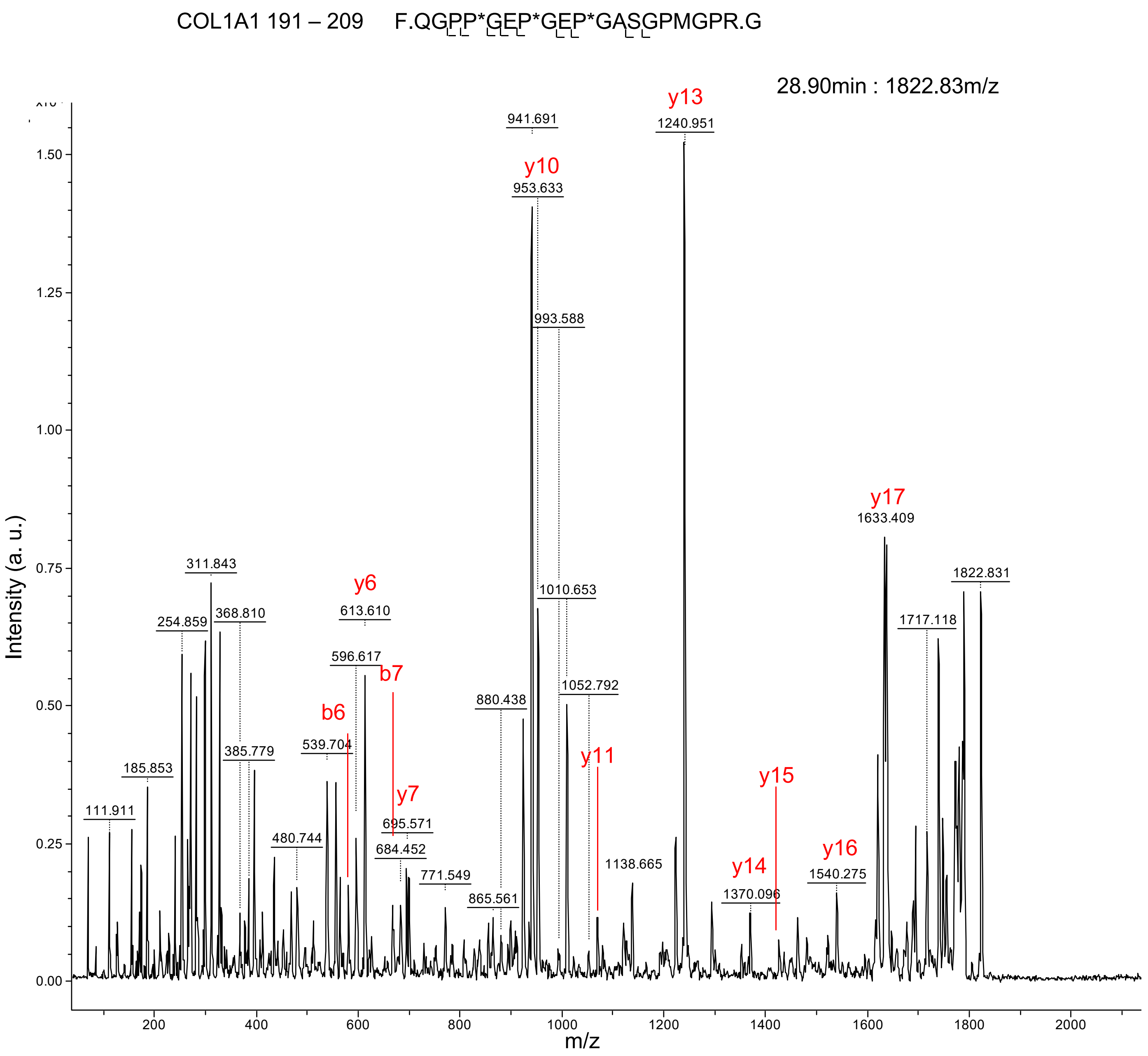
COL1A1 191 – 209 F.QGPP*GEP*GEP*GASGPMGPR.G
28.90min : 1822.83m/z
Intensity (a. u.)
m/z
y13
1240.951
941.691
y10
953.633
993.588
1010.653
1052.792
y17
1633.409
1822.831
1717.118
311.843
254.859
368.810
y6
613.610
596.617
b7
b6
385.779
539.704
185.853
111.911
480.744
y7
695.571
684.452
771.549
880.438
865.561
y11
1138.665
y15
y14
1370.096
y16
1540.275

A (continued)

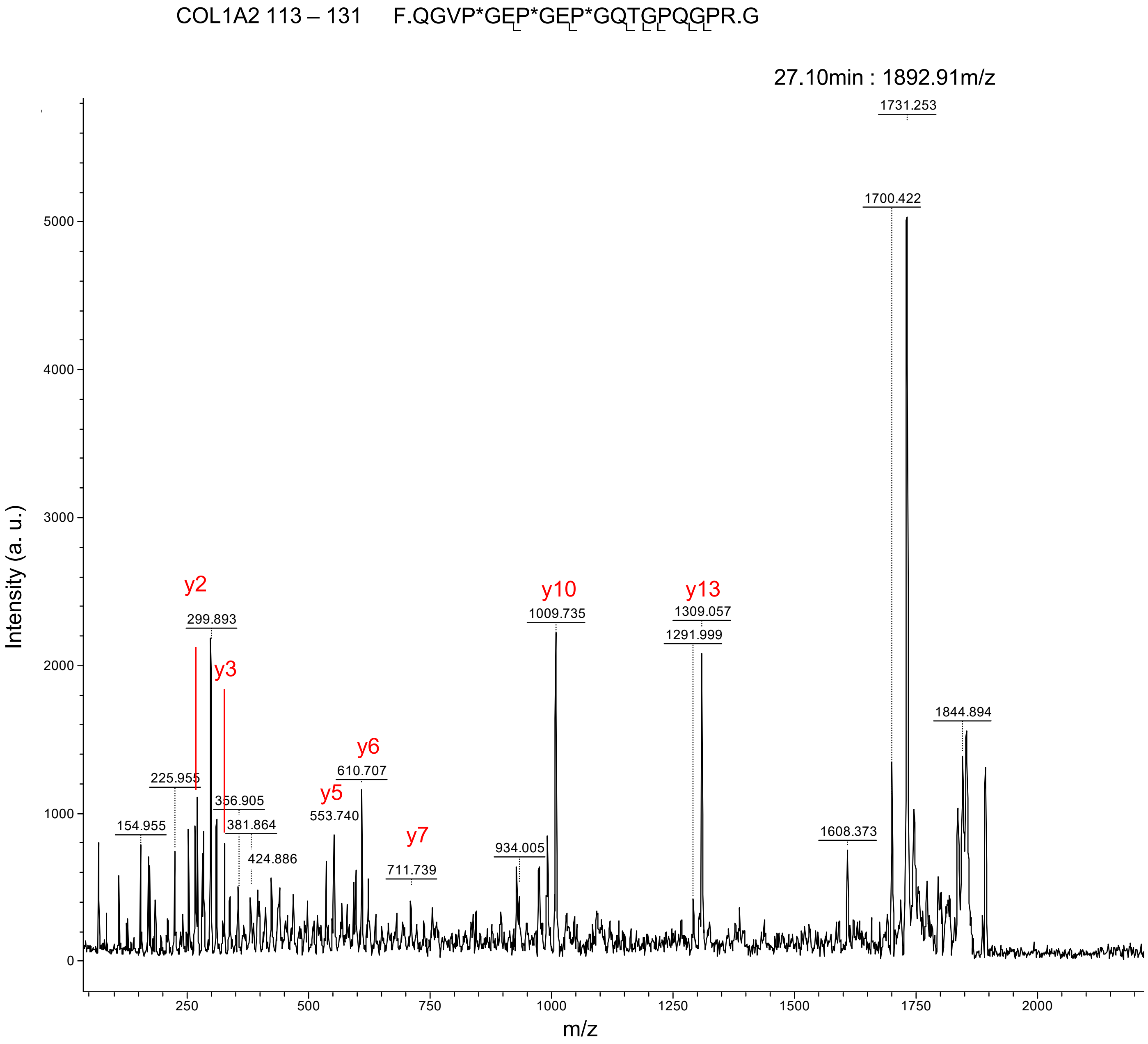

COL1A2 113 – 131 F.QGVP*GEP*GEP*GQTGPQGPR.G
27.10min : 1892.91m/z
1731.253
1700.422
1844.894
1608.373
y13
1309.057
1291.999
y10
1009.735
934.005
y7
711.739
y6
610.707
y5
553.740
y2
299.893
y3
225.955
356.905
381.864
424.886
154.955
Intensity (a. u.)
m/z
0
1000
2000
3000
4000
5000
250
500
750
1000
1250
1500
1750
2000

A (continued)

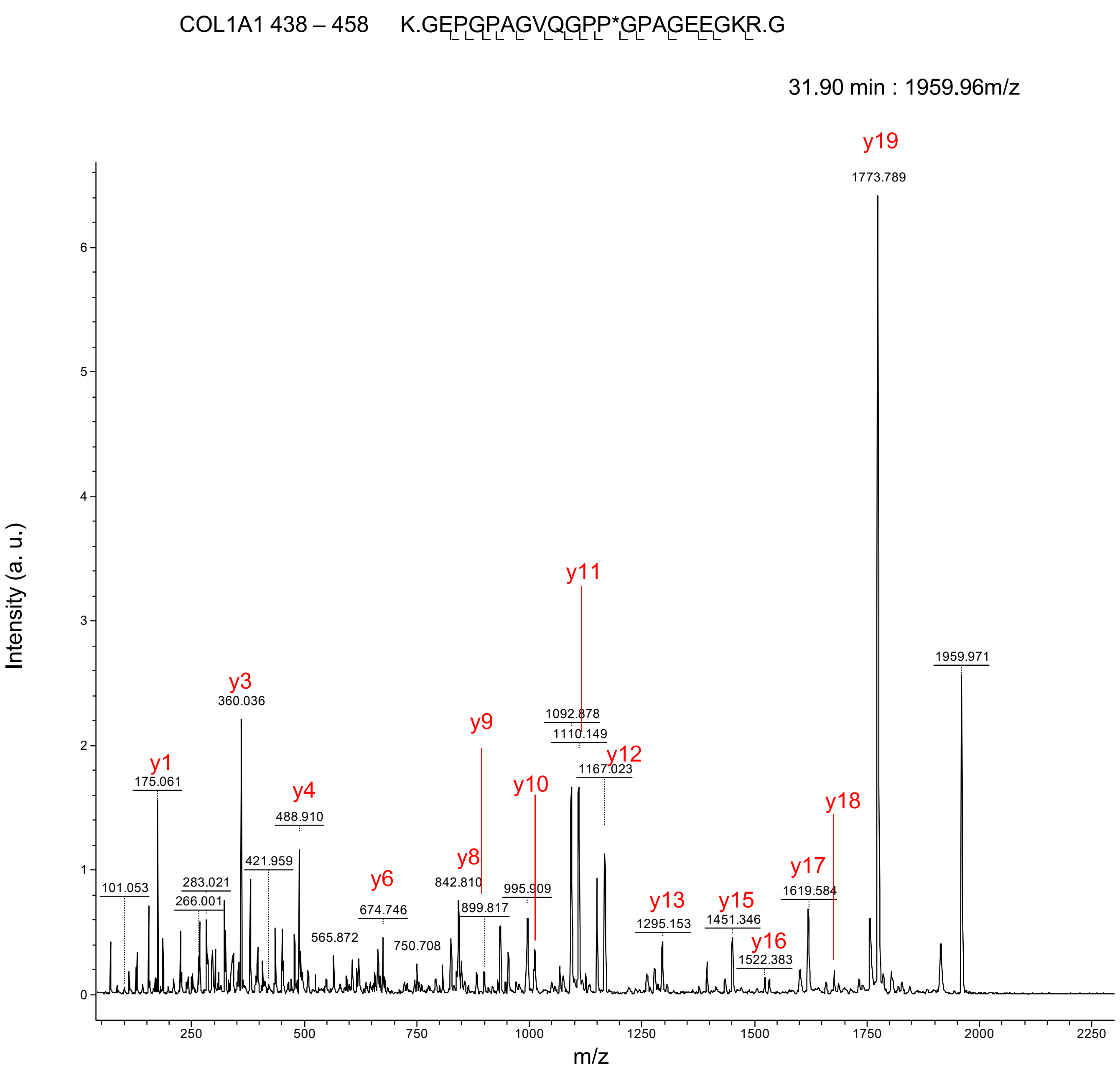
COL1A1 438 – 458 K.GEPGPAGVQGPP*GPAGEEGKR.G
31.90 min : 1959.96m/z
Intensity (a. u.)
m/z
y1
175.061
y3
360.036
y4
488.910
y6
674.746
y8
842.810
y9
899.817
y10
995.909
y11
1092.878
1110.149
y12
1167.023
y13
1295.153
y15
1451.346
y16
1522.383
y17
1619.584
y18
y19
1773.789
1959.971
101.053
283.021
266.001
421.959
565.872
750.708

A (continued)

COL1A1 438 – 458 K.GEP*GPAGVQGPP*GPAGEEGKR.G

31.90 min : 1959.96m/z

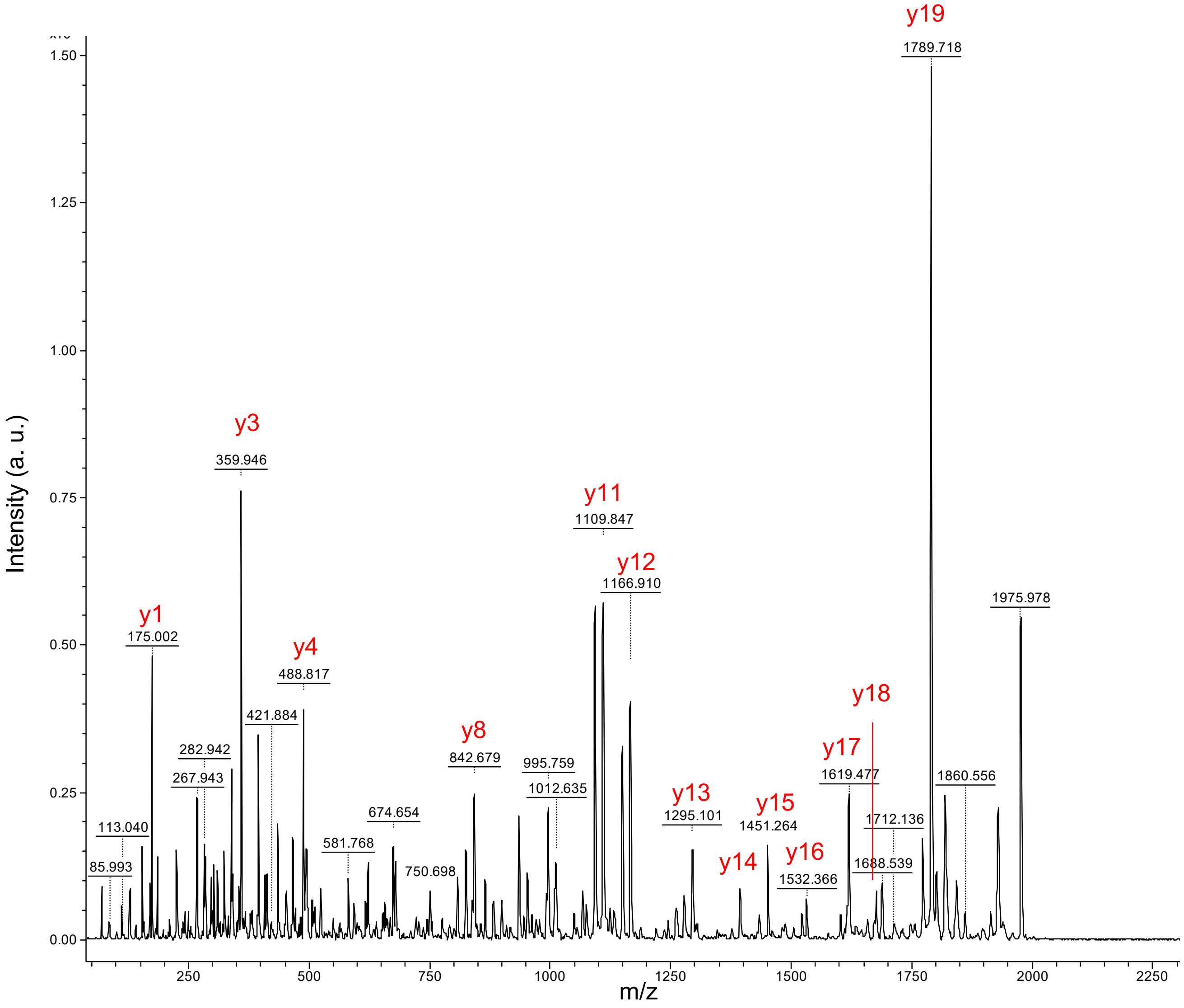

A (continued)

COL1A1 462 – 476 R.GEPGPAGLP*GPAGER.G

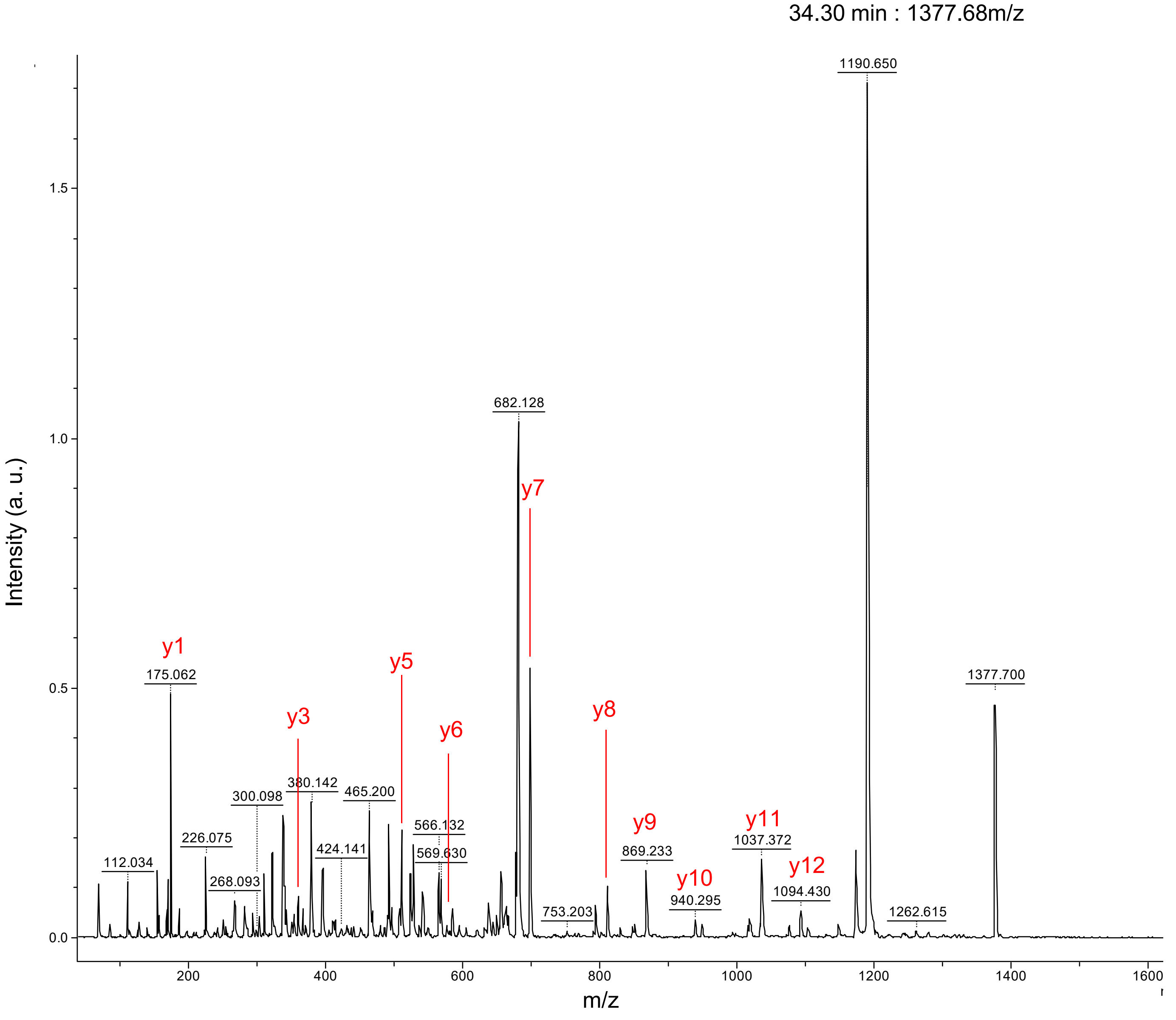

A (continued)

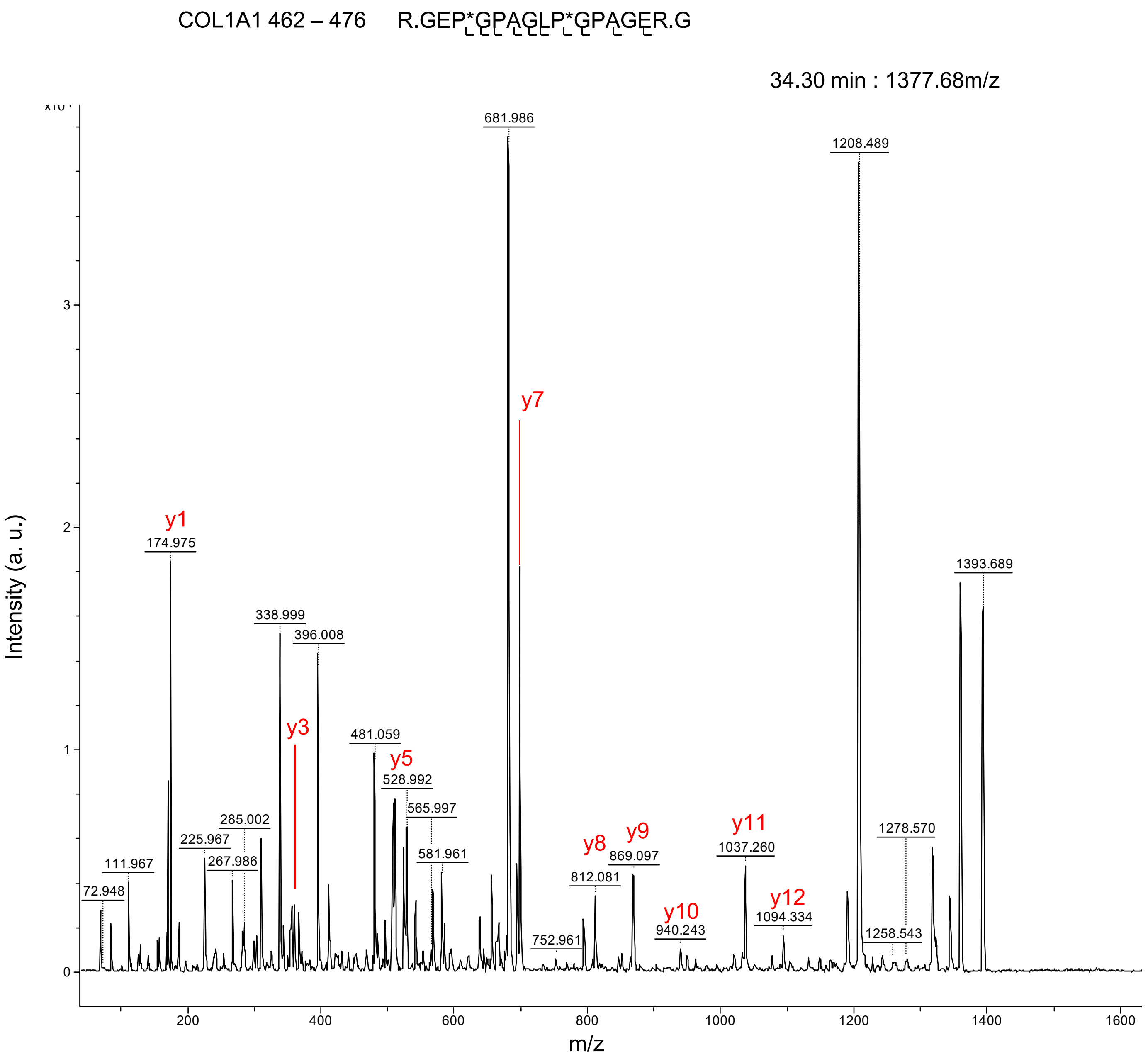

COL1A1 462 – 476 R.GEP*GPAGLP*GPAGER.G
34.30 min : 1377.68m/z
x10⁴
Intensity (a. u.)
m/z
681.986
1208.489
y7
y1
174.975
1393.689
338.999
396.008
y3
481.059
y5
528.992
565.997
285.002
225.967
267.986
111.967
72.948
581.961
y8
y9
812.081
869.097
y11
1037.260
1278.570
y10
940.243
y12
1094.334
752.961
1258.543

A (continued)

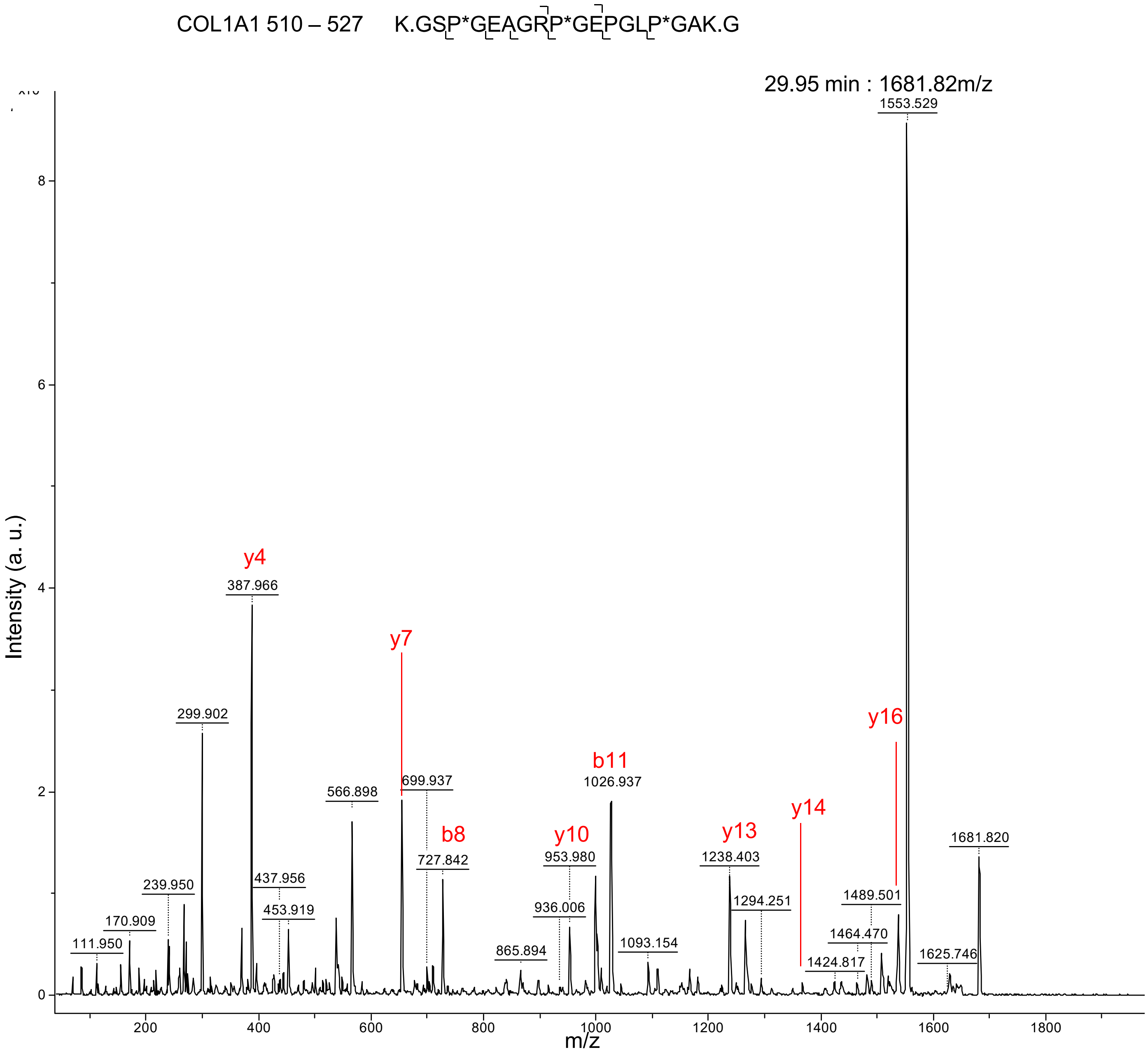

COL1A1 510 – 527 K.GSP*GEAGRP*GEPGLP*GAK.G
29.95 min : 1681.82m/z
Intensity (a. u.)
m/z
1553.529
y4
387.966
299.902
y7
699.937
566.898
b8
727.842
b11
1026.937
y10
953.980
936.006
y13
1238.403
1294.251
y14
y16
1681.820
1489.501
1464.470
1424.817
1625.746
1093.154
865.894
437.956
453.919
239.950
170.909
111.950
0
2
4
6
8
200
400
600
800
1000
1200
1400
1600
1800

A (continued)

COL1A1 510 – 527 K.GSP*GEAGRP*GEP*GLP*GAK.G

27.85 min : 1697.83m/z

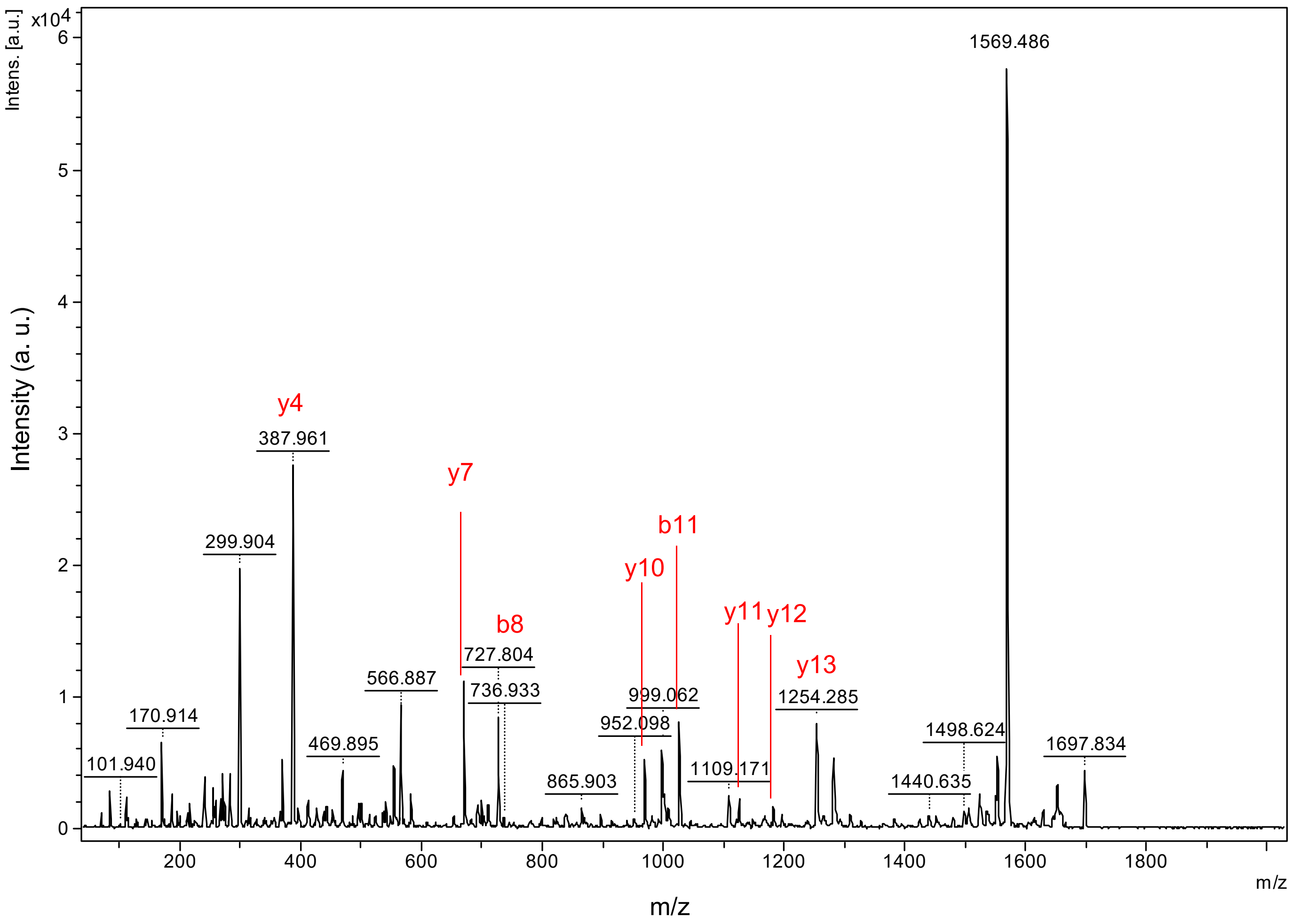

A (continued)

COL1A2 113 – 131   F.QGVPGEPGEP*GQTGPQGPR.G

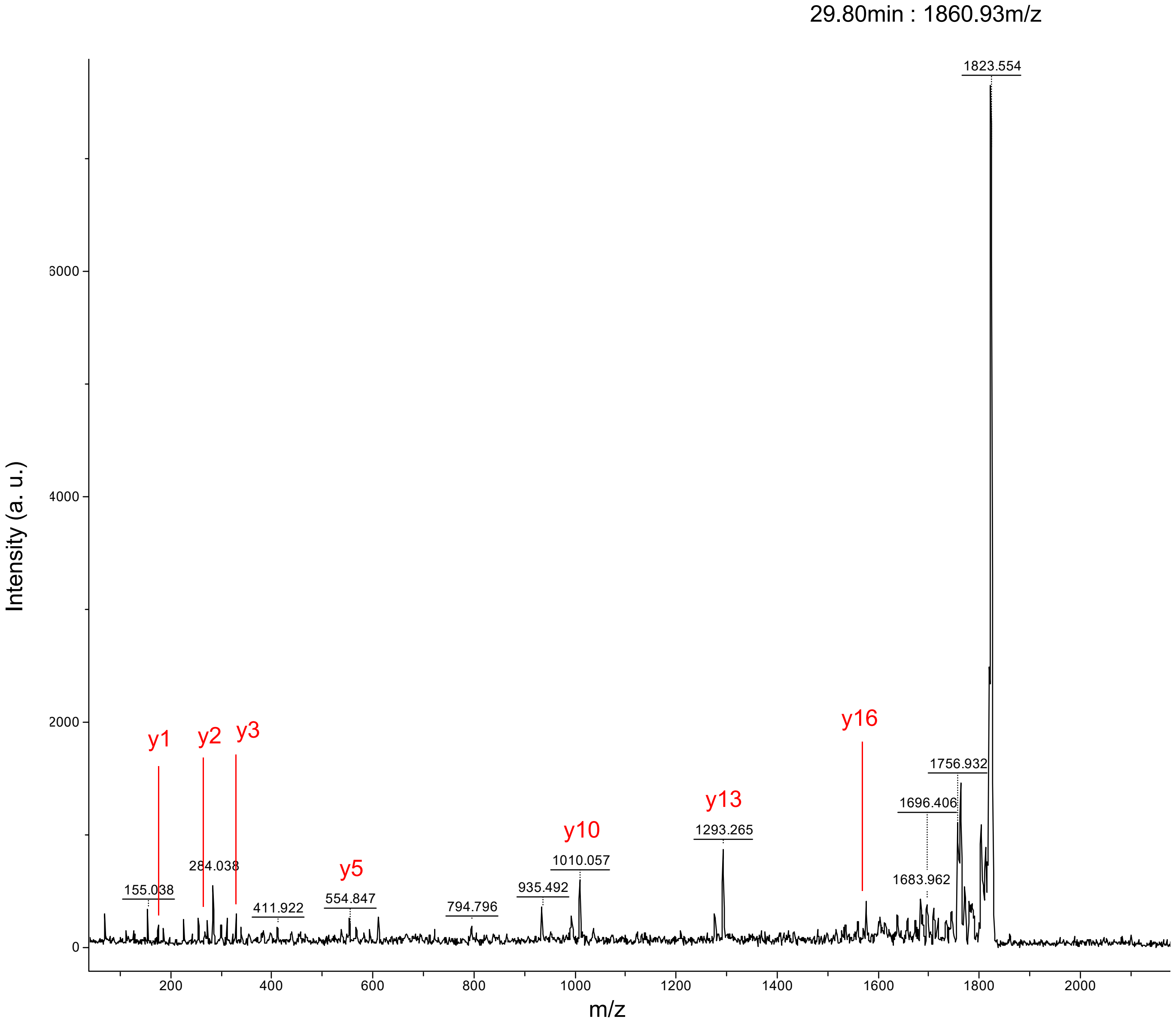

A (continued)

COL1A2 113 – 131 F.QGVPGEP*GEP*GQTGPQGPR.G

28.30min : 1876.93m/z

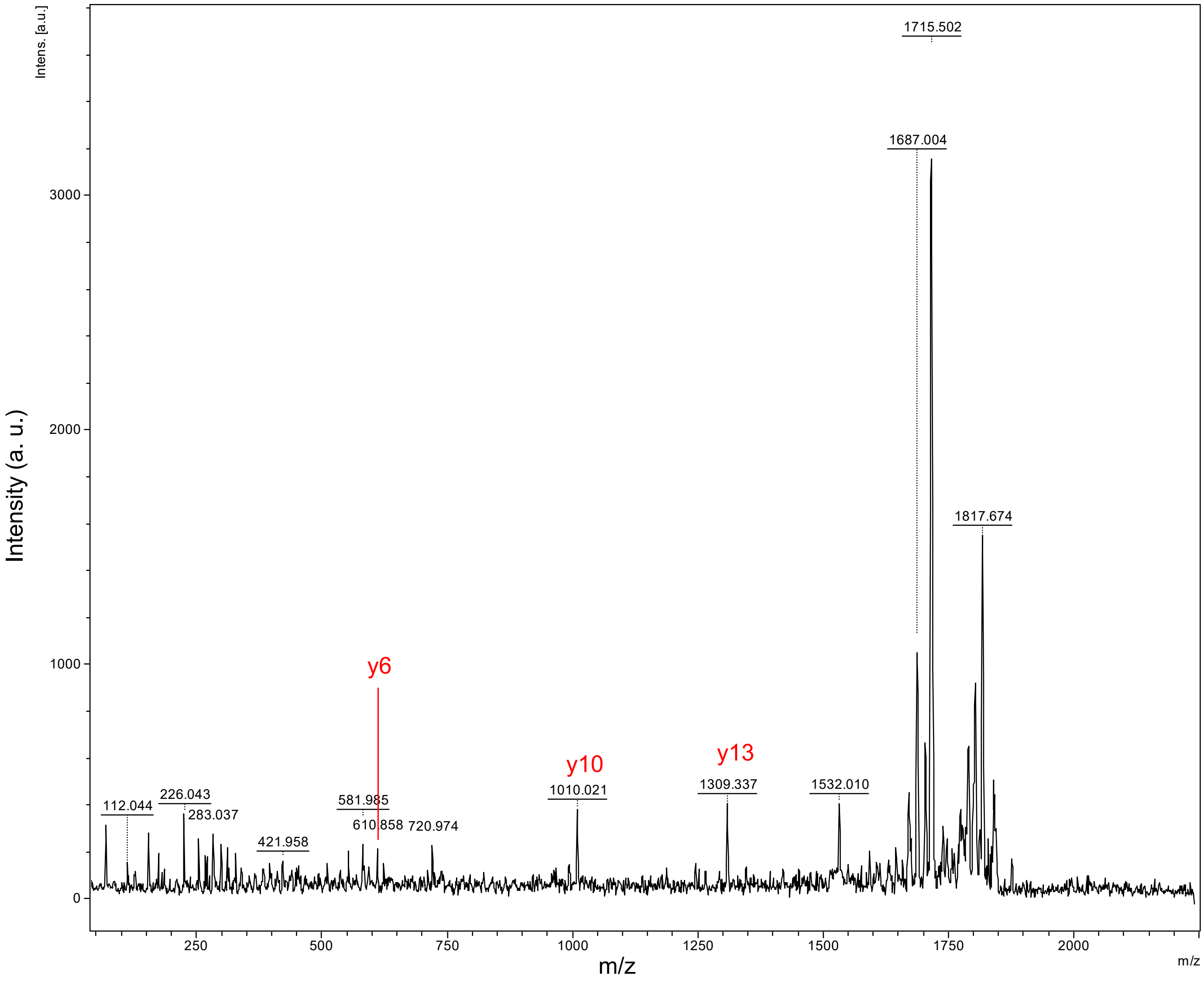

A (continued)

COL1A2 432 – 449 K.GPNGDAGRP*GEPGLMGPR.X

35.65min : 1750.83m/z

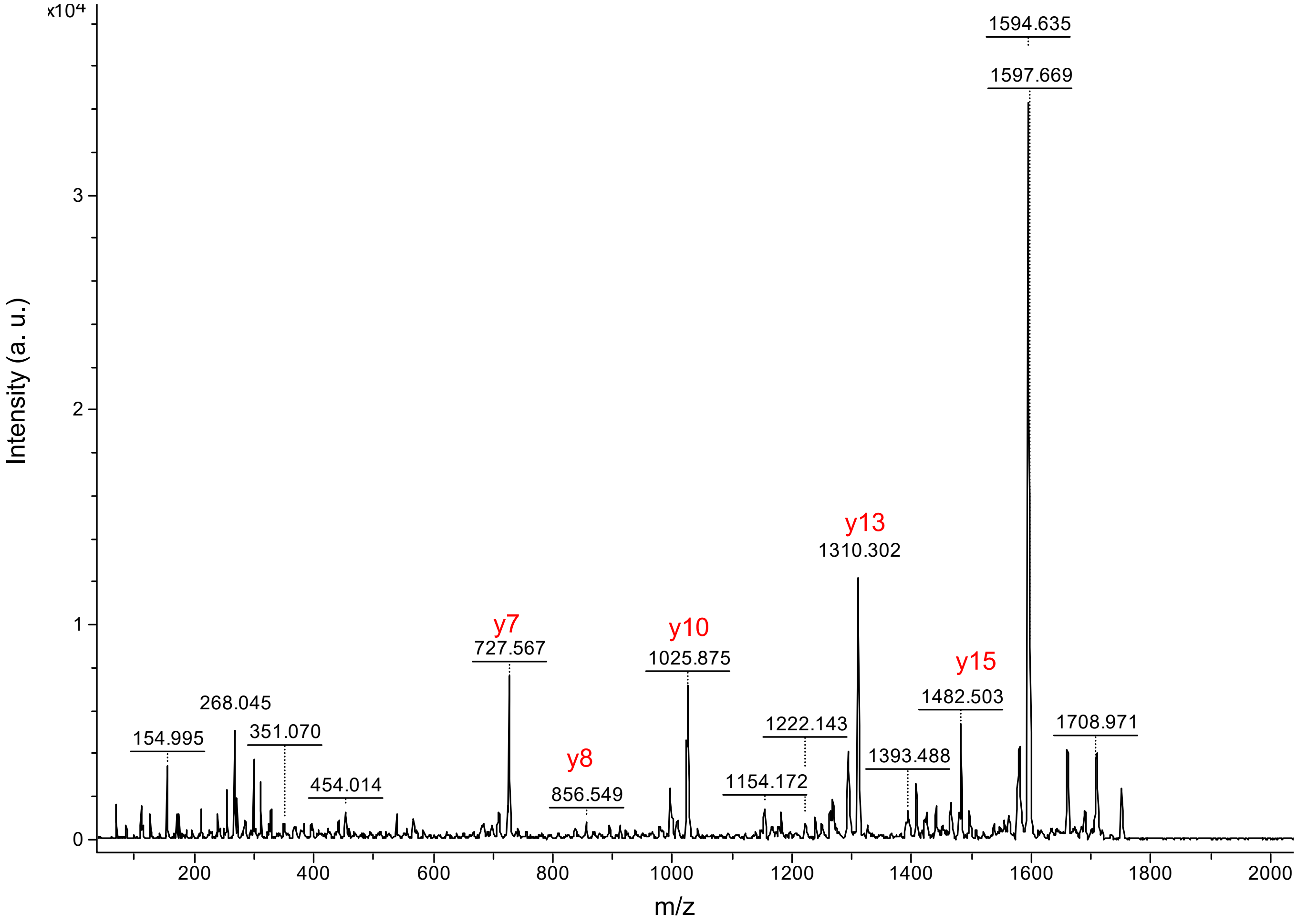

A (continued)

COL1A2 432 – 449    K.GPNGDAGRP*GEP*GLMGPR.X

34.00min : 1766.83m/z

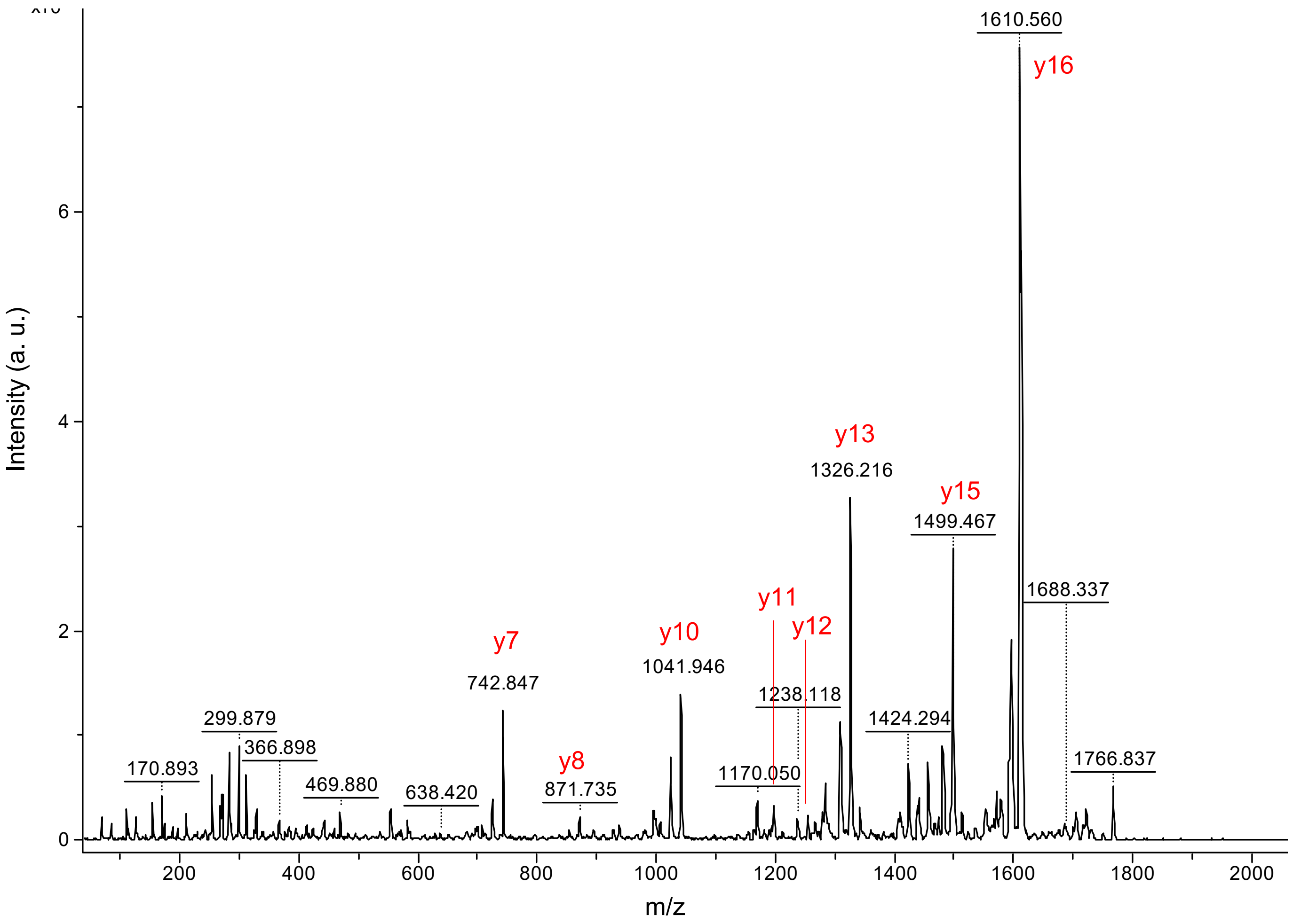

A (continued)

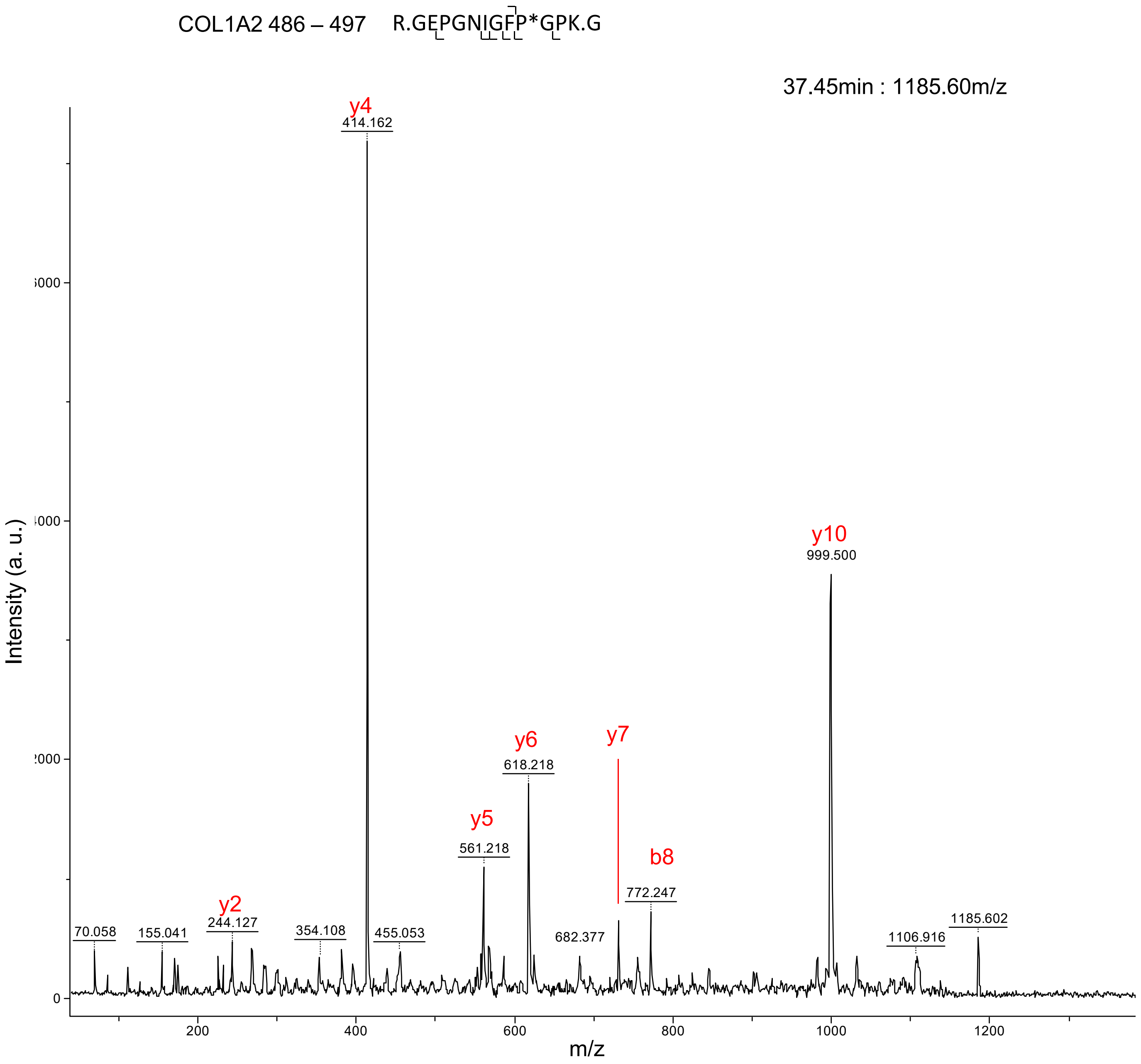
COL1A2 486 – 497 R.GEPGNIGFP*GPK.G
37.45min : 1185.60m/z
Intensity (a. u.)
m/z
y4
414.162
y10
999.500
y6
618.218
y7
y5
561.218
b8
772.247
y2
244.127
70.058
155.041
354.108
455.053
682.377
1106.916
1185.602
0
200
400
600
800
1000
1200

A (continued)

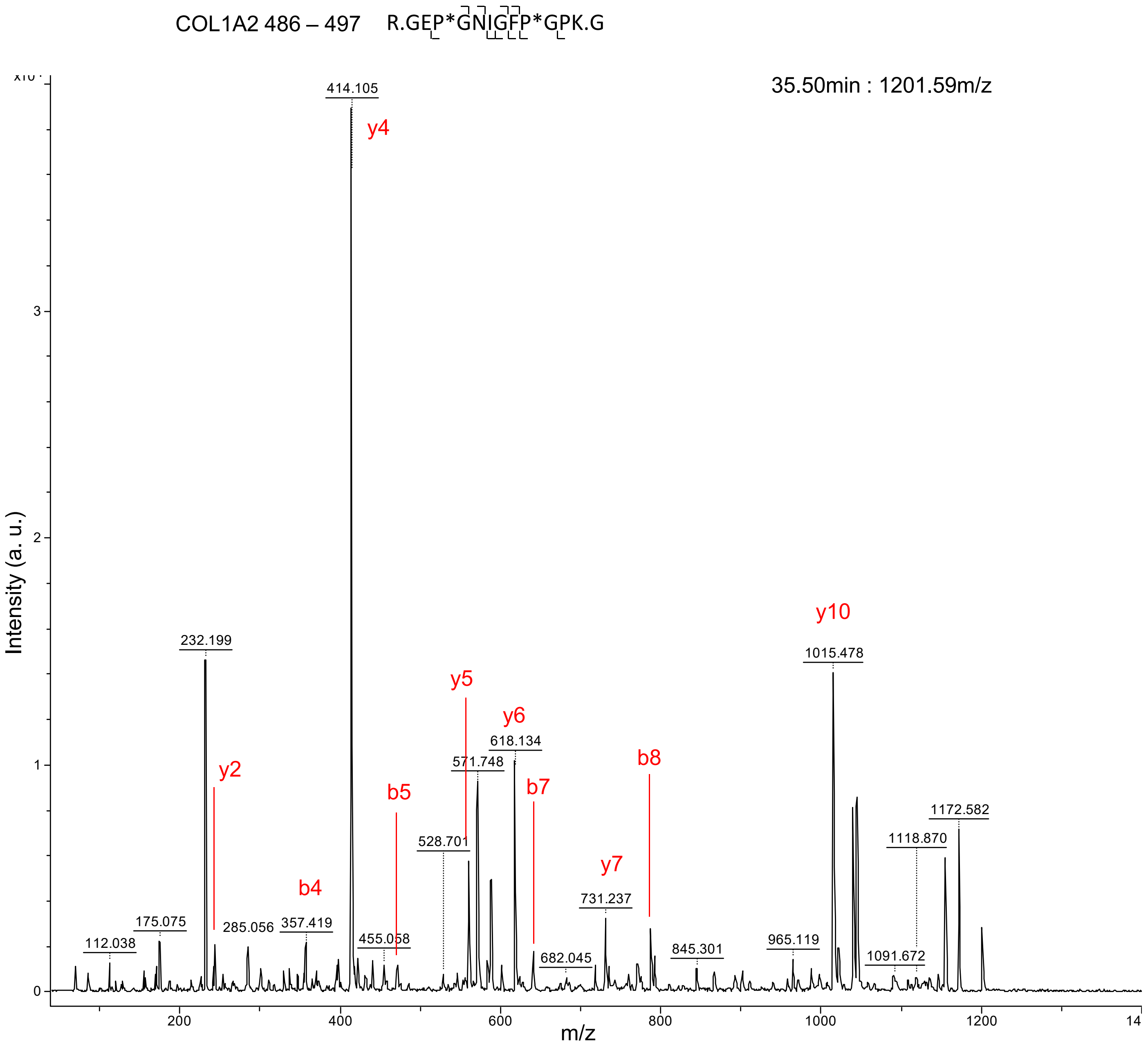

COL1A2 486 – 497 R.GEP*GNIGFP*GPK.G
35.50min : 1201.59m/z
Intensity (a. u.)
m/z
414.105
y4
232.199
y10
1015.478
y5
y6
618.134
571.748
y2
b5
b7
b8
528.701
1172.582
1118.870
b4
y7
731.237
175.075
285.056
357.419
112.038
455.058
682.045
845.301
965.119
1091.672
0
1
2
3
200
400
600
800
1000
1200
14

A (continued)

COL1A2 498 – 518 K.GPTGEPGKP*GEK*GNVGLAPGR.G

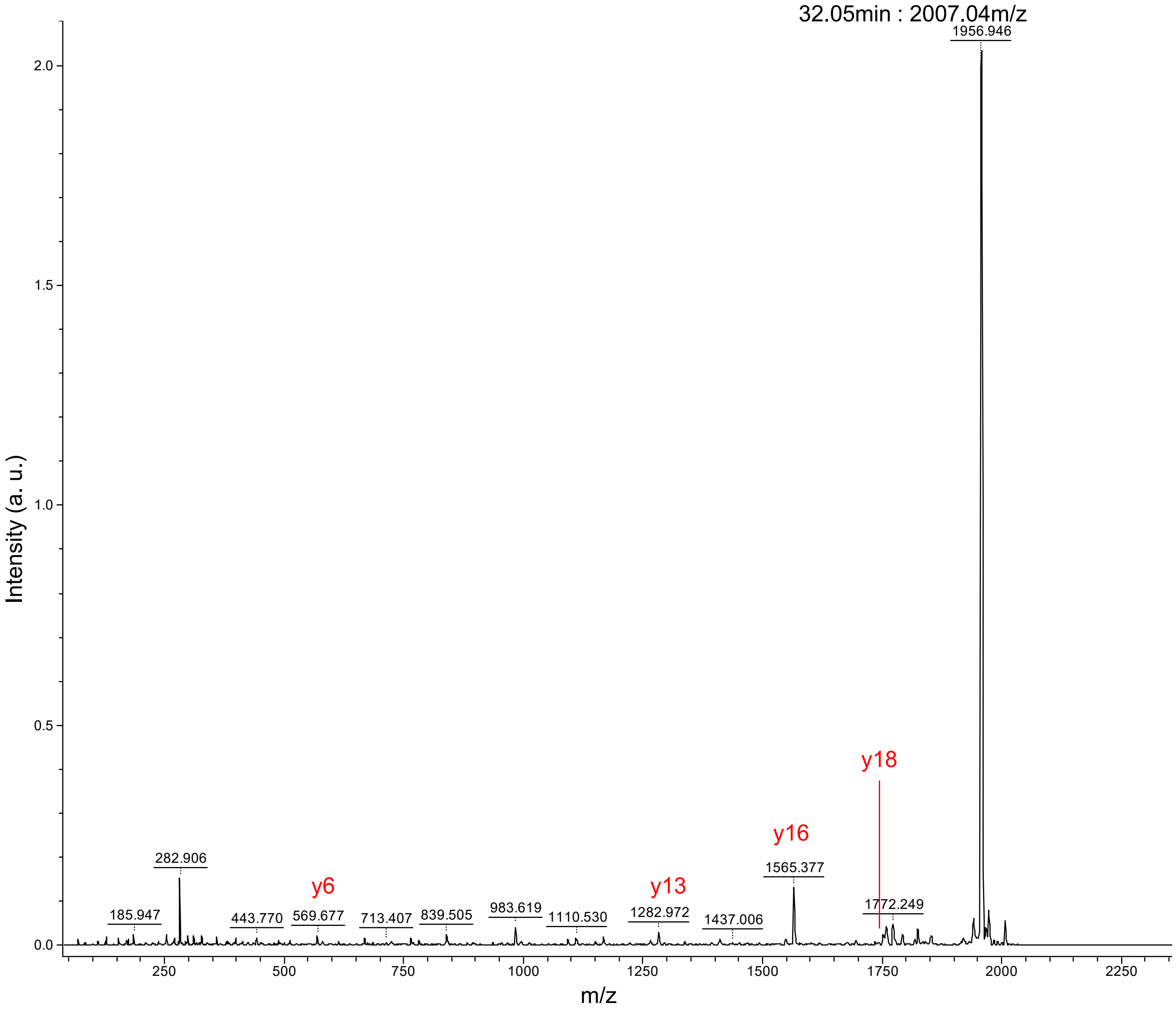

A (continued)

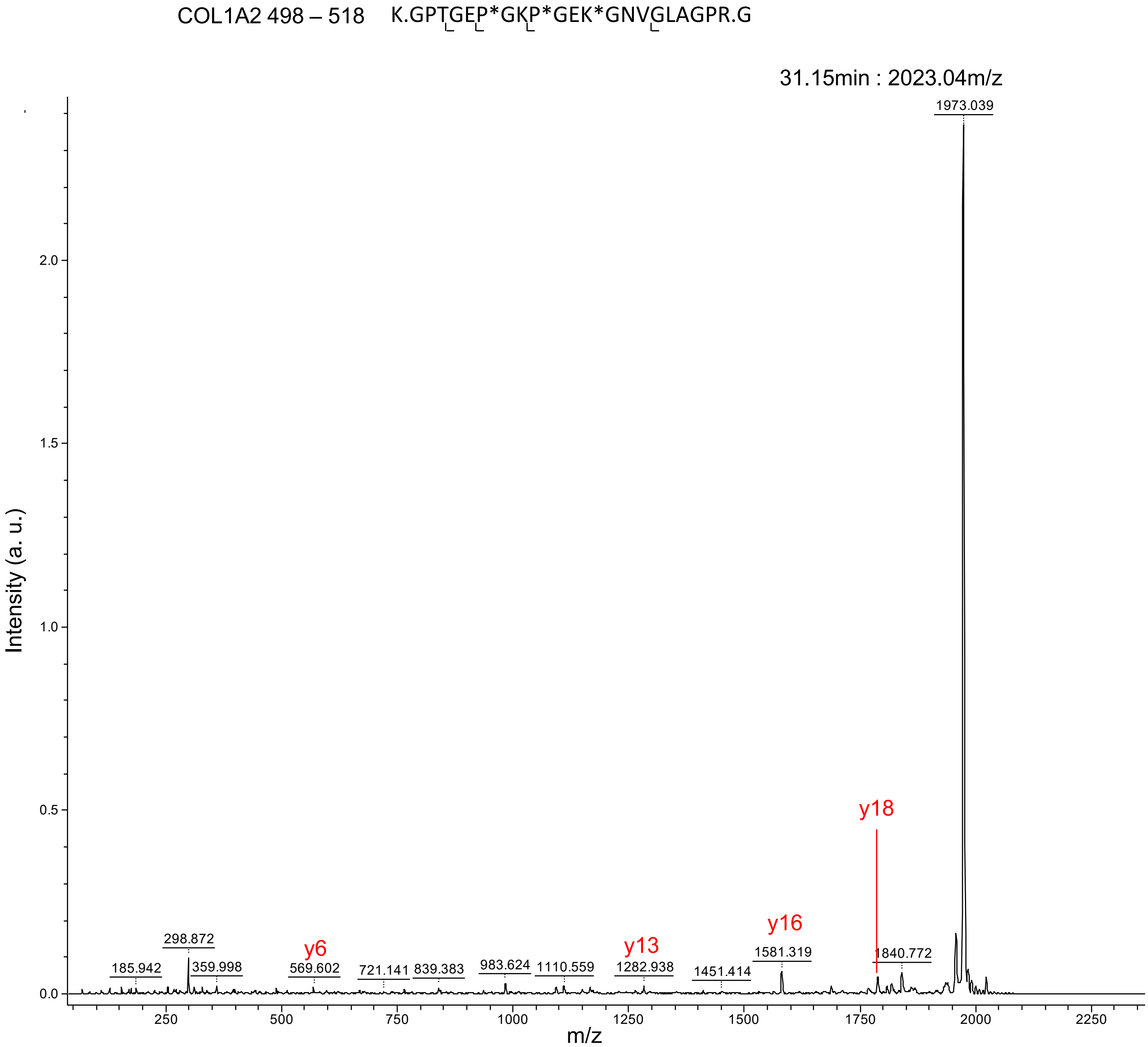
COL1A2 498 – 518 K.GPTGEP*GKP*GEK*GNVGLAGPR.G
31.15min : 2023.04m/z
1973.039
298.872
185.942
359.998
569.602
721.141
839.383
983.624
1110.559
1282.938
1451.414
1581.319
1840.772
y6
y13
y16
y18
Intensity (a. u.)
m/z
0.0
0.5
1.0
1.5
2.0
250
500
750
1000
1250
1500
1750
2000
2250

A (continued)

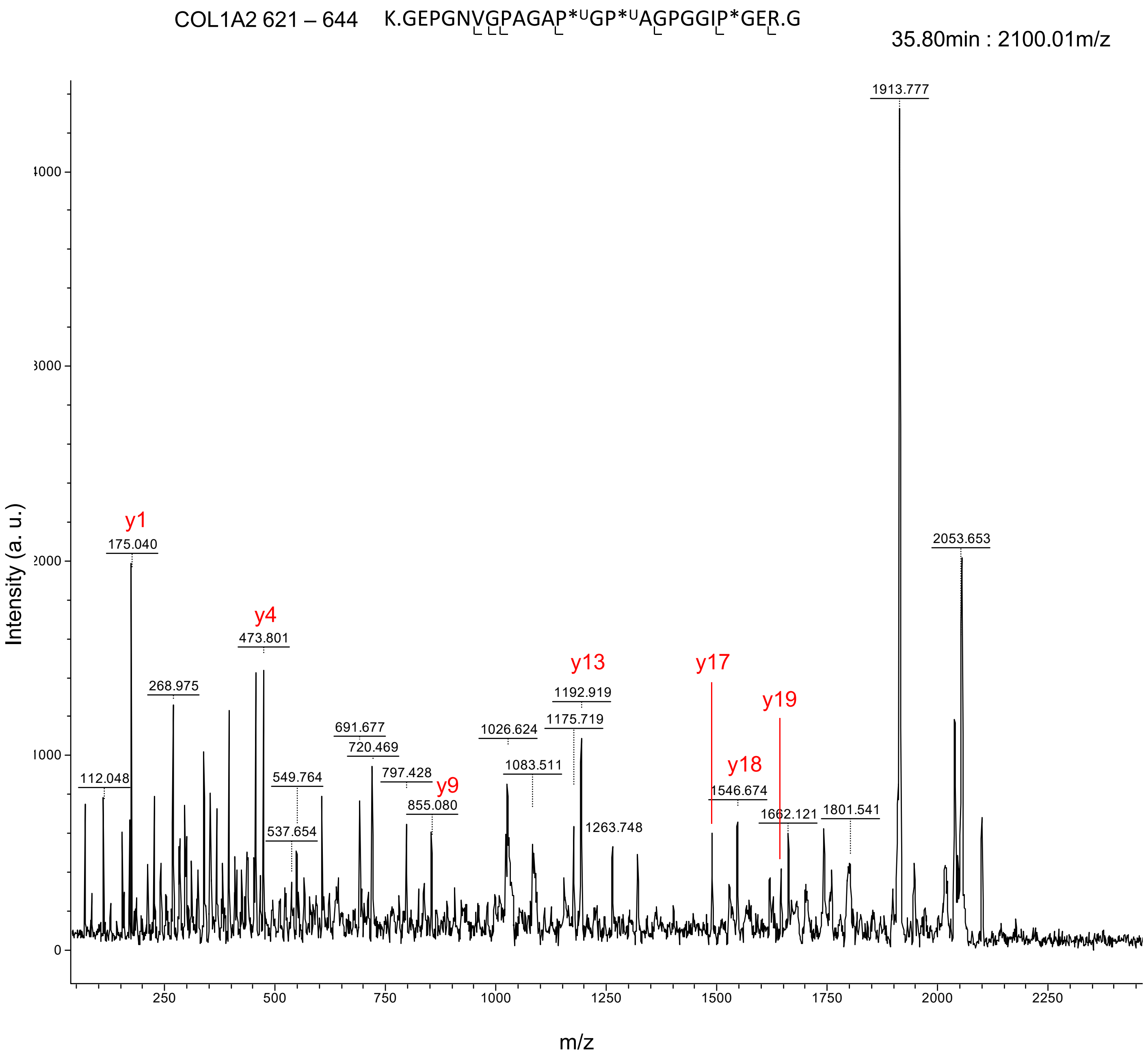

COL1A2 621 – 644 K.GEPGNVGPAGAP*UGP*UAGPGGIP*GER.G
35.80min : 2100.01m/z
Intensity (a. u.)
m/z
y1
175.040
y4
473.801
268.975
112.048
549.764
537.654
691.677
720.469
797.428
y9
855.080
1026.624
1083.511
y13
1192.919
1175.719
1263.748
y17
y18
1546.674
y19
1662.121
1801.541
1913.777
2053.653

A (continued)

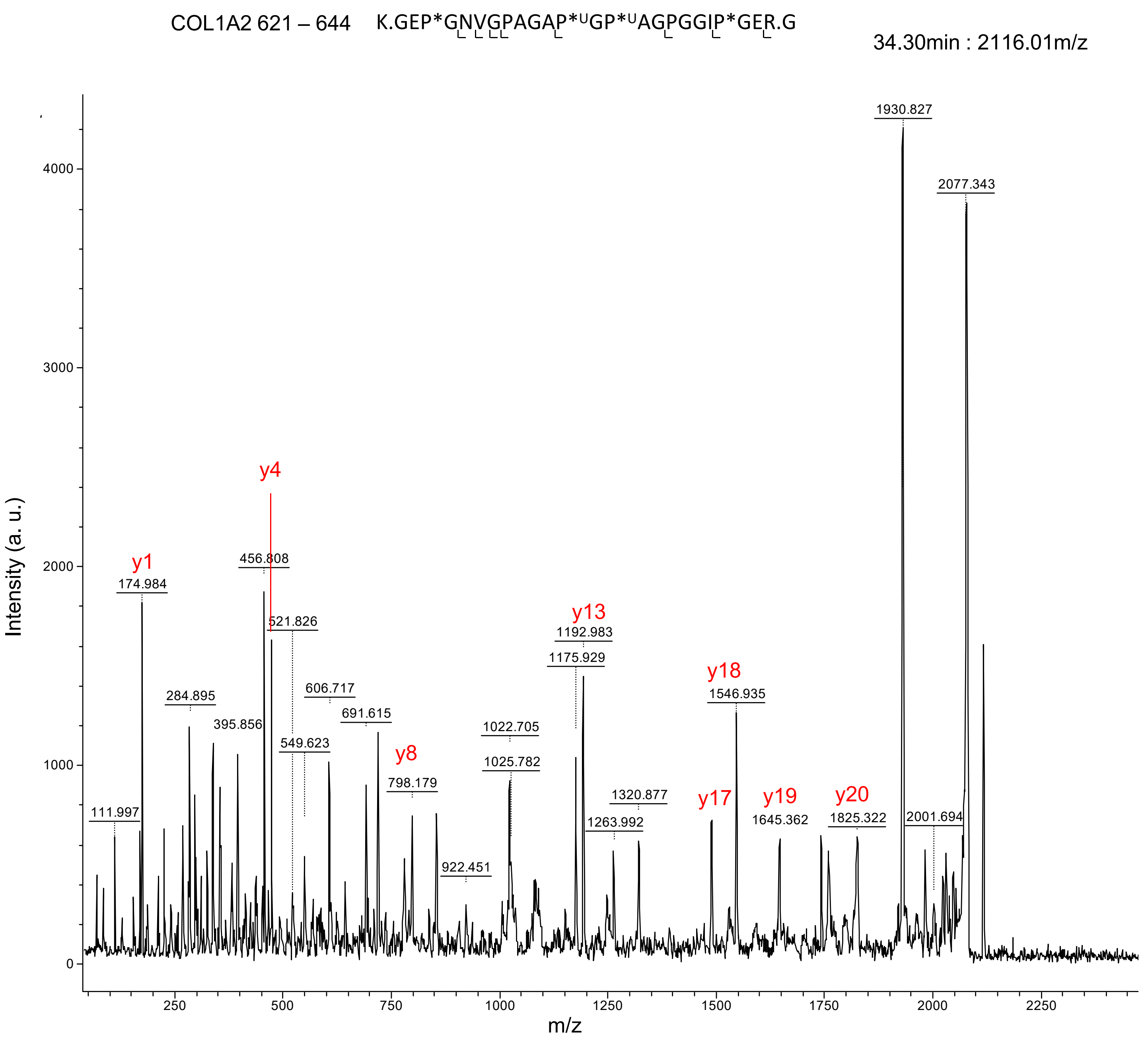

COL1A2 621 – 644 K.GEP*GNVGPAGAP*UGP*UAGPGGIP*GER.G
34.30min : 2116.01m/z
Intensity (a. u.)
m/z
y1
y4
y8
y13
y17
y18
y19
y20
111.997
174.984
284.895
395.856
456.808
521.826
549.623
606.717
691.615
798.179
922.451
1022.705
1025.782
1175.929
1192.983
1263.992
1320.877
1546.935
1645.362
1825.322
1930.827
2001.694
2077.343

A (continued)

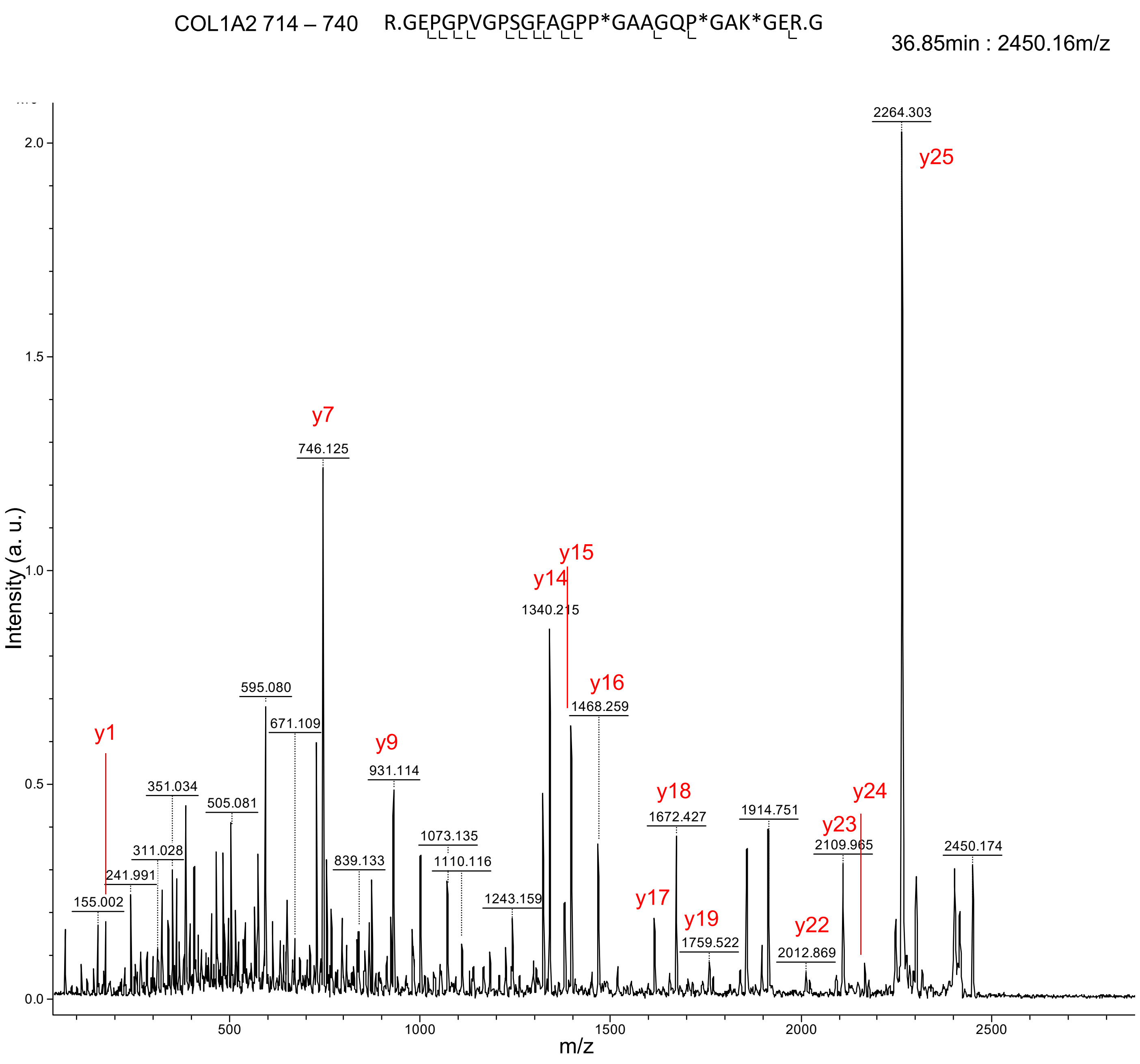
COL1A2 714 – 740
R.GEPGPVGPSGFAGPP*GAAGQP*GAK*GER.G
36.85min : 2450.16m/z
Intensity (a. u.)
m/z
2264.303
y25
y7
746.125
y14
y15
1340.215
y16
1468.259
595.080
671.109
y9
931.114
y1
351.034
505.081
311.028
241.991
155.002
839.133
1073.135
1110.116
1243.159
y18
1672.427
y17
y19
1759.522
1914.751
y24
y23
2109.965
y22
2012.869
2450.174
0.0
0.5
1.0
1.5
2.0
500
1000
1500
2000
2500

A (continued)

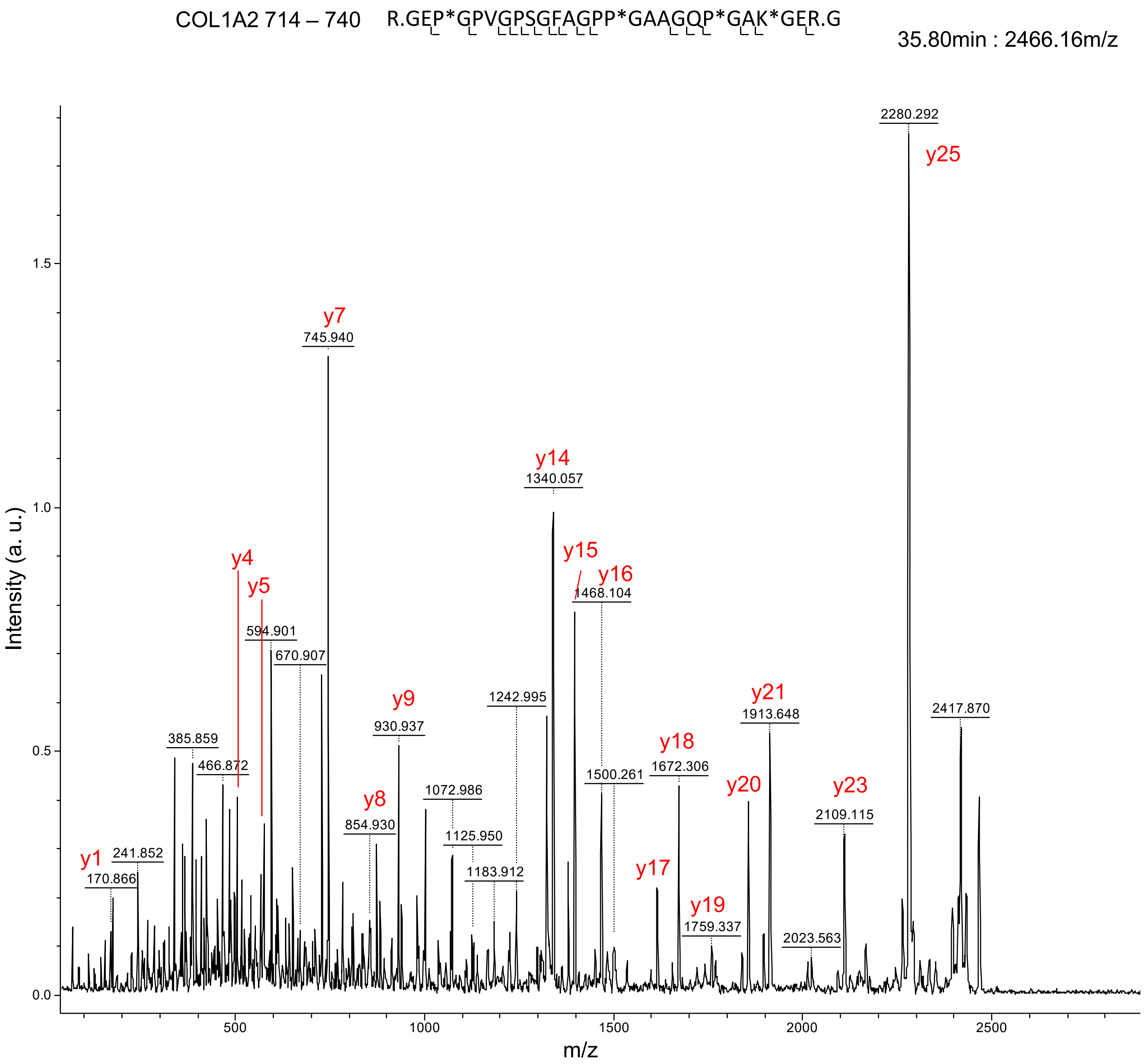

COL1A2 714 – 740 R.GEP*GPVGPSGFAGPP*GAAGQP*GAK*GER.G
35.80min : 2466.16m/z
Intensity (a. u.)
m/z
2280.292
y25
y7
745.940
y14
1340.057
y15
y16
1468.104
y4
y5
594.901
670.907
y9
930.937
1242.995
y21
1913.648
2417.870
385.859
466.872
y18
1672.306
1500.261
y20
y23
2109.115
y8
854.930
1072.986
1125.950
y1
241.852
170.866
1183.912
y17
y19
1759.337
2023.563
1.5
1.0
0.5
0.0
500
1000
1500
2000
2500

A (continued)

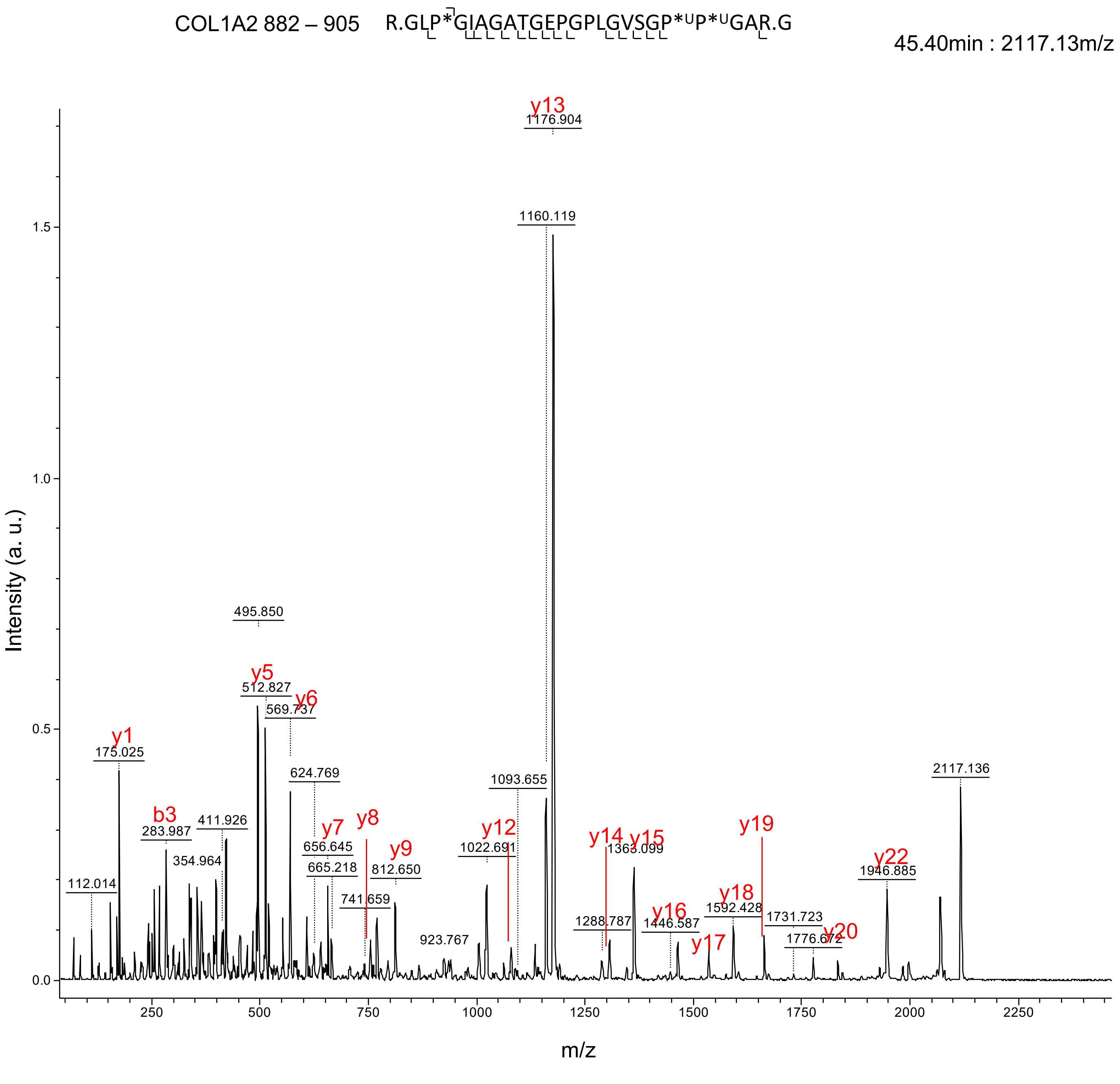
COL1A2 882 – 905 R.GLP*GIAGATGEPGPLGVSGP*ᵁP*ᵁGAR.G
45.40min : 2117.13m/z
y13
1176.904
1160.119
495.850
y5
512.827
y6
569.737
y1
175.025
624.769
1093.655
2117.136
b3
283.987
411.926
y7
y8
656.645
y9
y12
1022.691
y14
y15
1363.099
y19
y22
1946.885
354.964
665.218
812.650
112.014
741.659
923.767
1288.787
y16
1446.587
y18
1592.428
1731.723
y17
1776.672
y20
Intensity (a. u.)
m/z
0.0
0.5
1.0
1.5
250
500
750
1000
1250
1500
1750
2000
2250

A (continued)

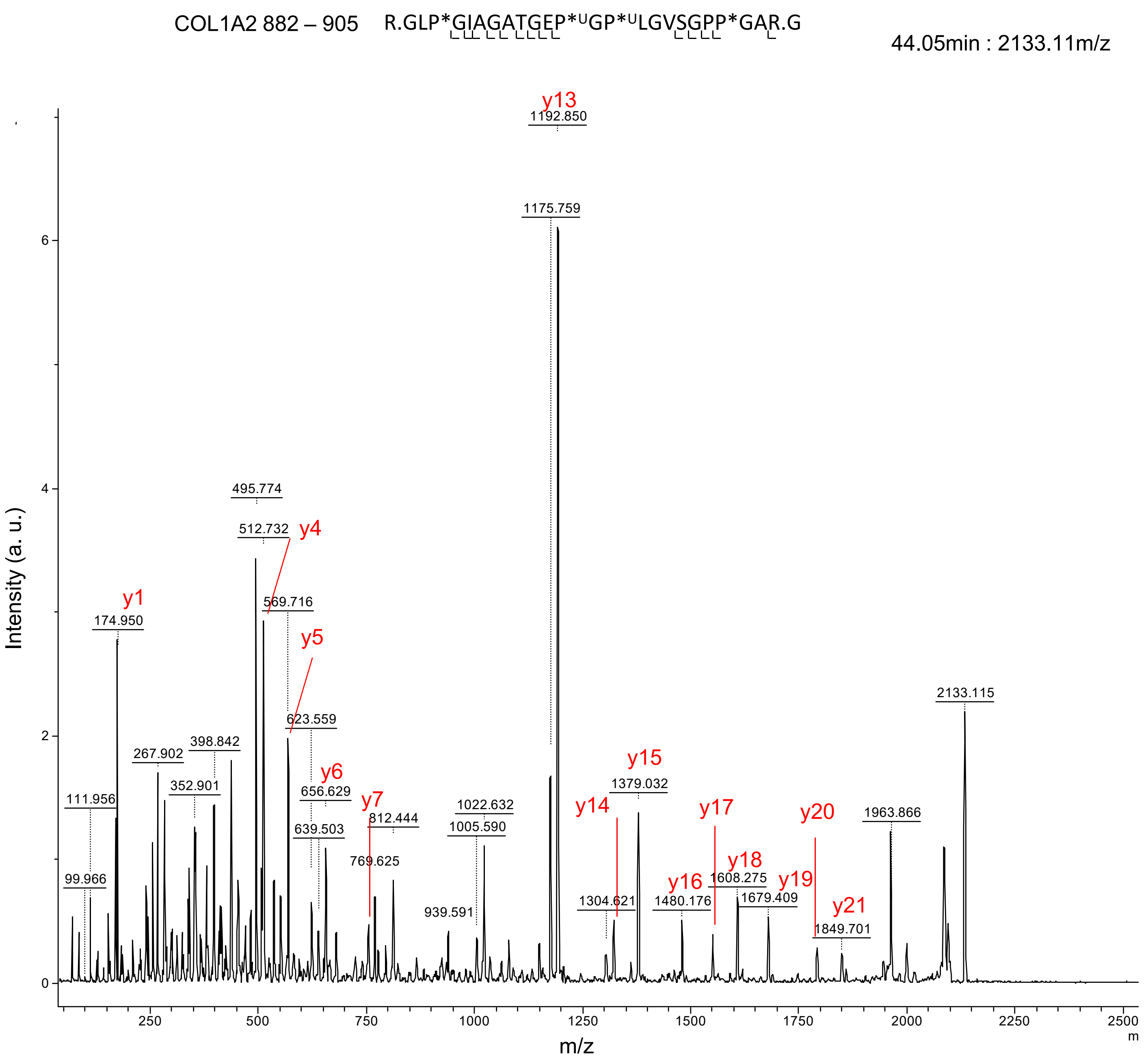

COL1A2 882 – 905 R.GLP*GIAGATGEP*UGP*ULGVSGPP*GAR.G
44.05min : 2133.11m/z
y13
1192.850
1175.759
495.774
512.732
y4
y1
174.950
569.716
y5
623.559
2133.115
267.902
398.842
352.901
y6
656.629
y15
1379.032
111.956
y7
812.444
1022.632
1005.590
y14
y17
y20
1963.866
639.503
769.625
y18
1608.275
y16
y19
1679.409
1480.176
1304.621
99.966
939.591
y21
1849.701
Intensity (a. u.)
m/z

A (continued)

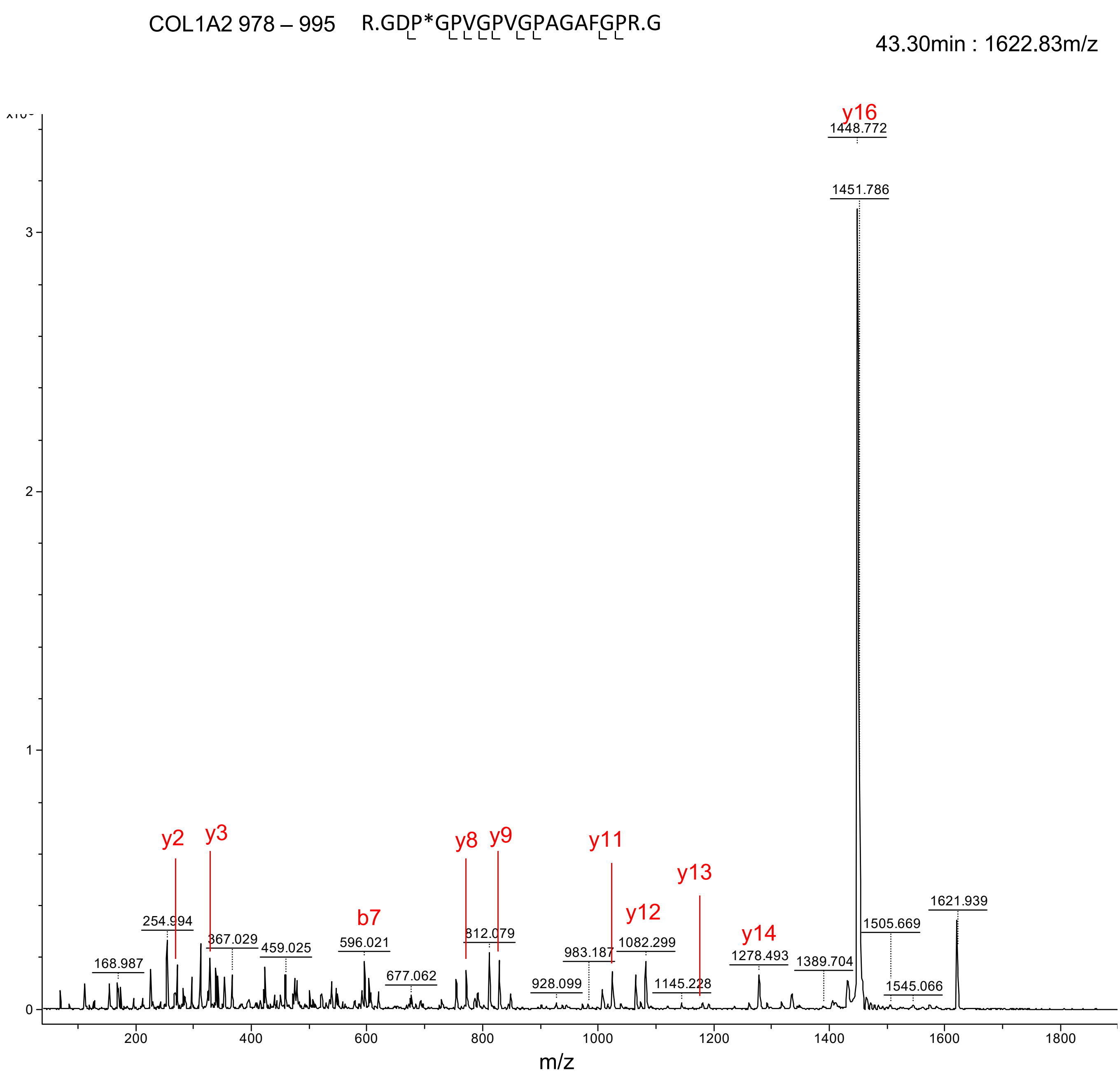

COL1A2 978 – 995 R.GDP*GPVGPVGPAGAFGPR.G
43.30min : 1622.83m/z
y16
1448.772
1451.786
y2
y3
y8
y9
y11
y12
y13
y14
b7
168.987
254.994
367.029
459.025
596.021
677.062
812.079
928.099
983.187
1082.299
1145.228
1278.493
1389.704
1505.669
1545.066
1621.939
0
1
2
3
200
400
600
800
1000
1200
1400
1600
1800
m/z

B

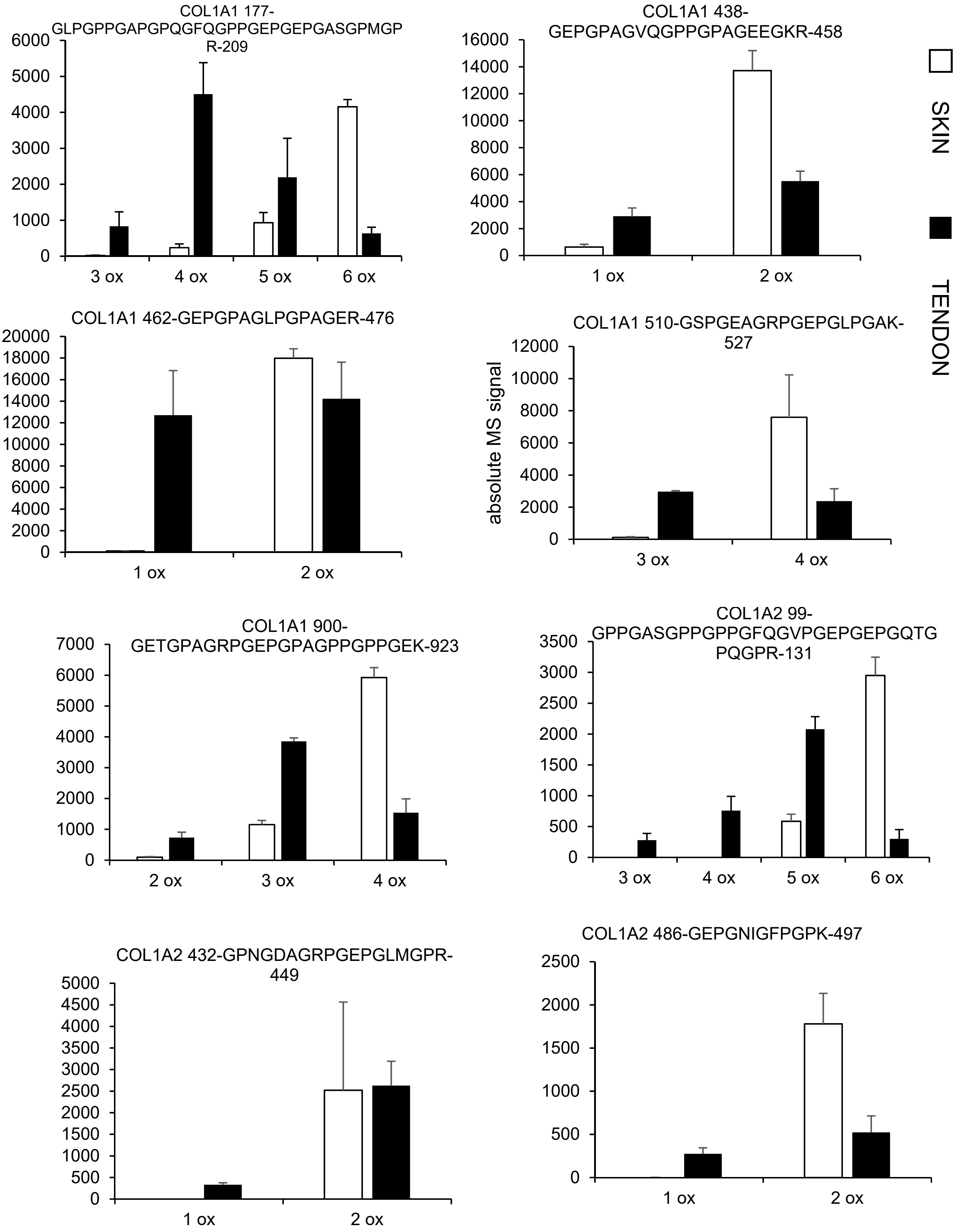

COL1A1 177-GLPGPPGAPGPQGFQGPPGEPGEPGASGPMGPR-209
6000
5000
4000
3000
2000
1000
0
3 ox
4 ox
5 ox
6 ox
COL1A1 438-GEPGPAGVQGPPGPAGEEGKR-458
16000
14000
12000
10000
8000
6000
4000
2000
0
1 ox
2 ox
SKIN
TENDON
COL1A1 462-GEPGPAGLPGPAGER-476
20000
18000
16000
14000
12000
10000
8000
6000
4000
2000
0
1 ox
2 ox
COL1A1 510-GSPGEAGRPGEPGLPGAK-527
absolute MS signal
12000
10000
8000
6000
4000
2000
0
3 ox
4 ox
COL1A1 900-GETGPAGRPGEPGPAGPPGPPGEK-923
7000
6000
5000
4000
3000
2000
1000
0
2 ox
3 ox
4 ox
COL1A2 99-GPPGASGPPGPPGFQGVPGEPGEPGQTGPQGPR-131
3500
3000
2500
2000
1500
1000
500
0
3 ox
4 ox
5 ox
6 ox
COL1A2 432-GPNGDAGRPGEPGLMGPR-449
5000
4500
4000
3500
3000
2500
2000
1500
1000
500
0
1 ox
2 ox
COL1A2 486-GEPGNIGFPGPK-497
2500
2000
1500
1000
500
0
1 ox
2 ox

B (continued)

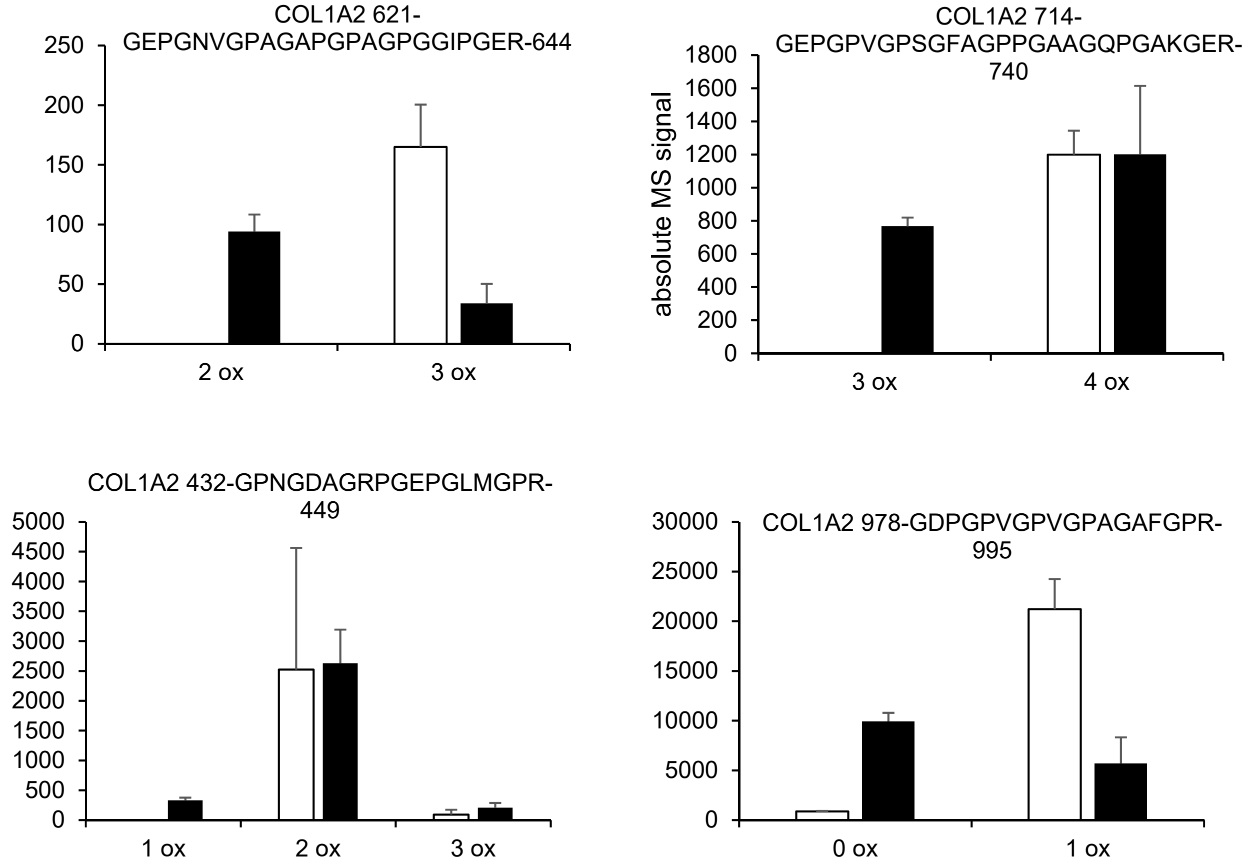
COL1A2 621-GEPGNVGPAGAPGPAGPGGIPGER-644
250
200
150
100
50
0
2 ox
3 ox
COL1A2 714-GEPGPVGPSGFAGPPGAAGQPGAKGER-740
absolute MS signal
1800
1600
1400
1200
1000
800
600
400
200
0
3 ox
4 ox
COL1A2 432-GPNGDAGRPGEPGLMGPR-449
5000
4500
4000
3500
3000
2500
2000
1500
1000
500
0
1 ox
2 ox
3 ox
COL1A2 978-GDPGPVGPVGPAGAFGPR-995
30000
25000
20000
15000
10000
5000
0
0 ox
1 ox

C

| | residue positions | peptide sequence | hydroxylation # skin | tendon | p-value |
|---|---|---|---|---|---|
| chicken COL1A1 | 177 - 209 | GLP$^{1}$GPP$^{2}$GAP$^{3}$GPQGFQGPP$^{5R}$ GEP$^{6R}$GEP$^{4}$GASGPMGPR | 5.73 | 4.32 | 3.95E-4 |
| | 276 - 293 | GEPGSPGENGAPGQMGPR | NI | | |
| | 438 - 458 | GEP$^{2R}$GPAGVQGPP$^{1}$GPAGEEG KR | 1.96 | 1.66 | 4.96E-6 |
| | 462 - 476 | GEP$^{2R}$GPAGLP$^{1}$GPAGER | 1.99 | 1.53 | 4.47E-3 |
| | 510 - 527 | GSP$^{3}$GEAGRP$^{3}$GEP$^{4R}$GLP$^{3}$GAK | 3.98 | 3.44 | 3.66E-4 |
| | 816 - 824 | GETGDAGAK | ND | | |
| | 900 - 923 | GETGPAGRPGEP$^{4R}$GPAGP$^{3U}$P$^{3U}$GP$^{3U}$P$^{3U}$GEK.G | 3.81 | 3.13 | 6.67E-6 |
| | 984 - 1003 | GPPGPMGPPGLAGPPGEAGR | 2.92 | 2.89 | NS |
| chicken COL1A2 | 99 – 131 | GPP$^{3}$GASGP$^{3}$P$^{3}$GPPGFQGVP$^{6R}$ GEP$^{5R}$GEP$^{4}$GQTGPQGPR | 5.84 | 4.7 | 3.88E-4 |
| | 198 - 215 | GEPGAPGENGTPGQPGAR | NI | | |
| | 360 - 380 | GEPGAAGPPGPPGPSGEEGKR | NI | | |
| | 432 - 449 | GP$^{3}$NGDAGRP$^{1}$GEP$^{2R}$GLMGPR | 2* | 1.89* | 1.09E-3 |
| | 486 - 497 | GEP$^{2R}$GNIGFP$^{1}$GPK | 2 | 1.65 | 2.54E-3 |
| | 498 - 509 | GPTGEP$^{3R}$GKP$^{2}$GEK | 1.94 | 1.39 | 1.05E-4 |
| | 573 - 587 | GLHGEFGVPGPAGPR | 1.19 | 1.16 | NS |
| | 621 - 644 | GEP$^{3R}$GNVGPAGAPGP$^{2U}$AGP$^{2U}$ GGIP$^{1}$GER | 3 | 2.26 | 3.17E-4 |
| | 714 - 740 | GEP$^{4R}$GPVGPSGFAGPP$^{3}$GAAG QP$^{3}$GAK$^{3}$GER | 4 | 3.6 | 1.82E-3 |
| | 846 - 878 | GPSGEAGAAGPPGTPGPQGILG APGILGLPGSR | 4 | 4 | NS |
| | 882 - 905 | GLP$^{2}$GIAGATGEP$^{3RU}$GP$^{3RU}$LGVS GP$^{2U}$P$^{2U}$GAR | 2.94 | 2.2 | 2.21E-6 |
| | 978 - 995 | GDP$^{1R}$GPVGPVGPAGAFGPR | 0.96 | 0.36 | 1.27E-3 |
