## supplemental data S3 for "Tissue-specific collagen hydroxylation at GEP/GDP triplets mediated by P4HA2"

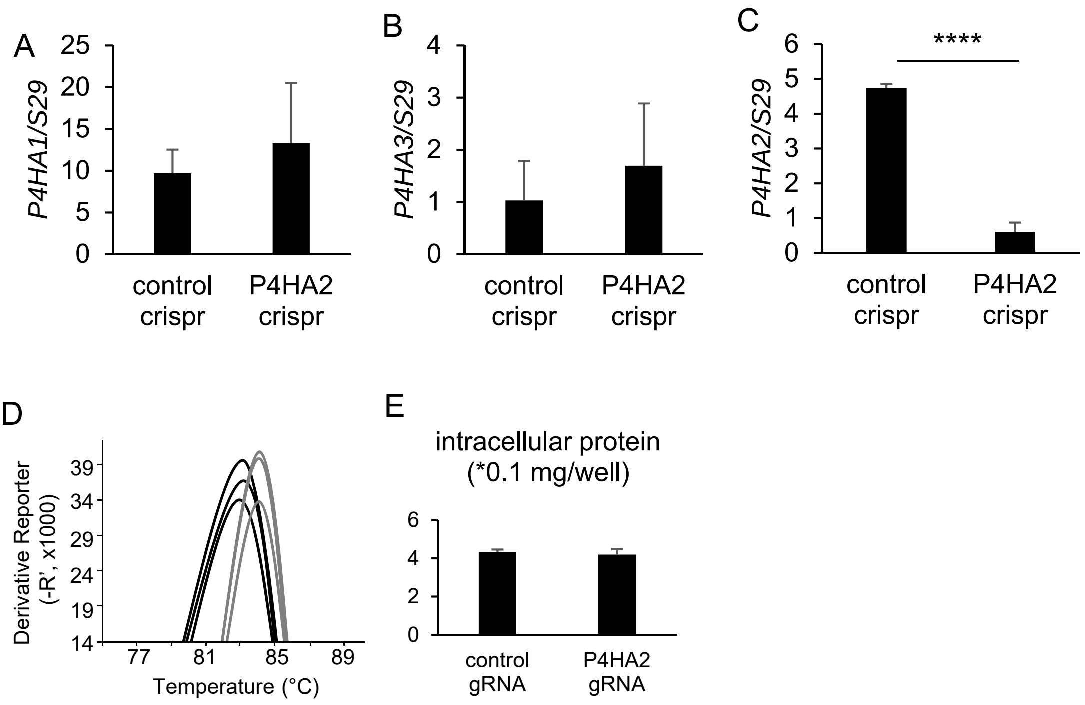


**Supplemental data S3. genetic invalidation of *P4HA2* in the ATDC5 model examined by qRT-PCR.** (A) Expression of *P4HA1* in both experimental groups. (B) Expression of *P4HA3* in in both experimental groups. (C) Expression of *P4HA2* in in both experimental groups. **** indicates t-test p-value < 0.00005. (D) Examination of the melting curve obtained for P4HA2 amplicons, showing that the remaining *P4HA2* mRNA expressed in the P4HA2-targetted group exhibit a lower melting temperature due to in/dels, as originally described in (Samarut *et. al*. BMC Genomics 2016). Clear grey, control gRNA replicates; dark grey, *P4HA2* gRNA replicates. The *P4HA2* amplificon contains the gRNA-targeted region. (E) Bradford assay showing similar total intracellular protein content in both experimental groups at the end of the culture period.(n=3).
