## supplemental data S4 for "Tissue-specific collagen hydroxylation at GEP/GDP triplets mediated by P4HA2"

**Supplemental data S4. Peptide details in ATDC5 cells upon P4HA2 invalidation.** (A-B) relative quantification of the distinct molecular states of each peptide found modulated, either in the CNBr extract ((A) or, when unavailable in the CNBr extract, in the culture supernatent (B). Throughout panels A-B, *, **, ***, **** indicate t-test p-values < 0.05, 0.005, 0.0005 or 0.00005 versus control. (n=3 for panel A, n=4 for panel B). White bars: control cultures; black bars: P4HA2-invalidated cells. (C) Table summarizing quantifications per peptide. Peptides are classified per their amino acid boundaries, referring to their positions in the $\alpha$1 (COL1A1) or $\alpha$2 chain (COL1A2) of type I mouse collagen precursor. All peptides were determined from tryptic digests of either collagen CNBr-extracts (white background) or culture supernantants (grey background). The amino acid sequence is also given, GEP/GDP triplets being highlighted in red. The average hydroxylation number of each peptide in either control or P4HA2-invalidated ATDC5 cells is calculated from total ion chromatograms. ND: no data available; NS: non-significant. t-test p-value is calculated for control versus P4HA2-invalidated cells (n=3 per group for data obtained from the CNBr extract, n=4 for data obtained from culture supernatents).

A

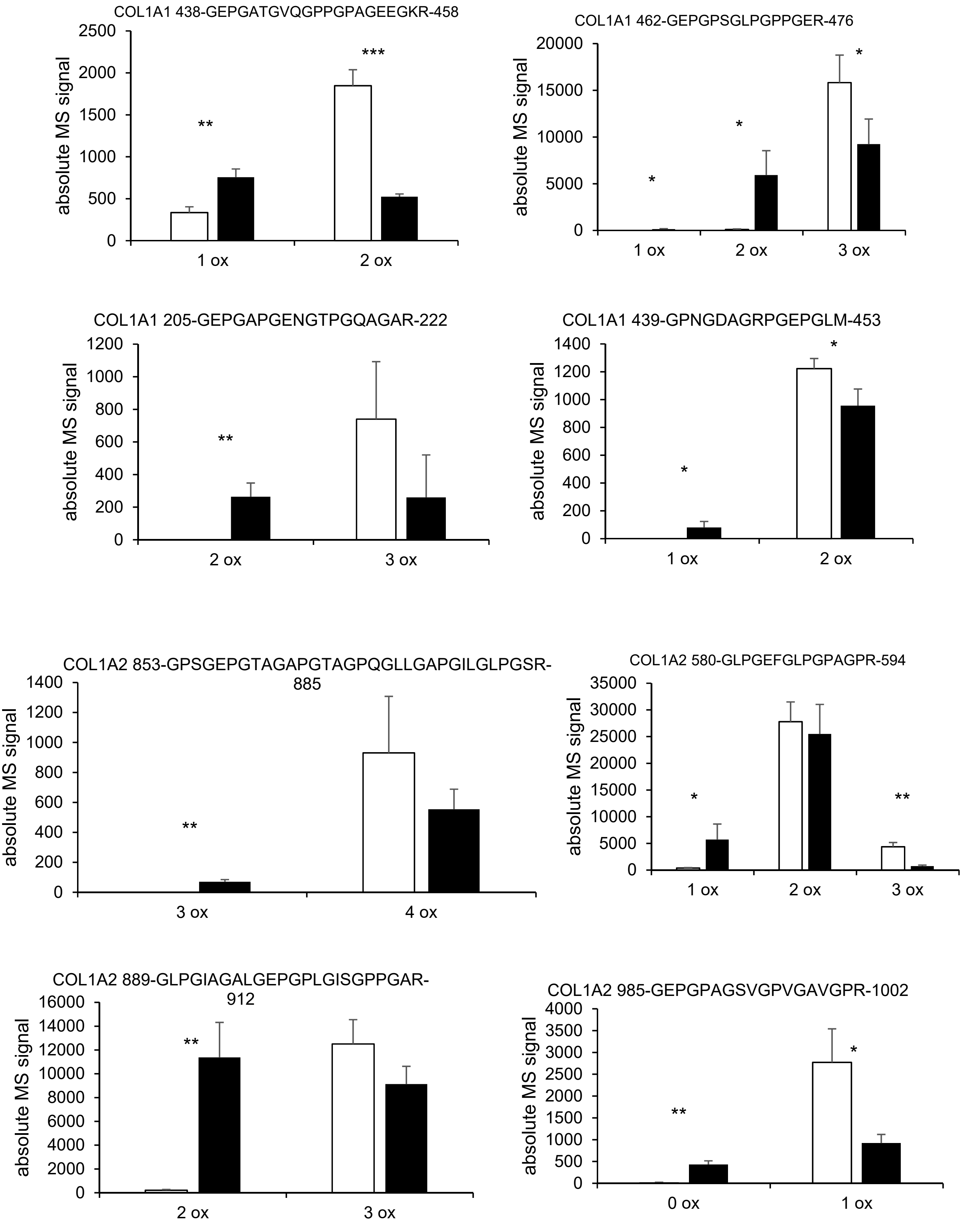
COL1A1 438-GEPGATGVQGPPGPAGEEGKR-458
absolute MS signal
2500
2000
1500
1000
500
0
**
***
1 ox
2 ox
COL1A1 462-GEPGPSGLPGPPGER-476
absolute MS signal
20000
15000
10000
5000
0
*
*
*
1 ox
2 ox
3 ox
COL1A1 205-GEPGAPGENGTPGQAGAR-222
absolute MS signal
1200
1000
800
600
400
200
0
**
2 ox
3 ox
COL1A1 439-GPNGDAGRPGEPGLM-453
absolute MS signal
1400
1200
1000
800
600
400
200
0
*
*
1 ox
2 ox
COL1A2 853-GPSGEPGTAGAPGTAGPQGLLGAPGILGLPGSR-885
absolute MS signal
1400
1200
1000
800
600
400
200
0
**
3 ox
4 ox
COL1A2 580-GLPGEFGLPGPAGPR-594
absolute MS signal
35000
30000
25000
20000
15000
10000
5000
0
*
**
1 ox
2 ox
3 ox
COL1A2 889-GLPGIAGALGEPGPLGISGPPGAR-912
absolute MS signal
16000
14000
12000
10000
8000
6000
4000
2000
0
**
2 ox
3 ox
COL1A2 985-GEPGPAGSVGPVGAVGPR-1002
absolute MS signal
4000
3500
3000
2500
2000
1500
1000
500
0
**
*
0 ox
1 ox

A (continued)

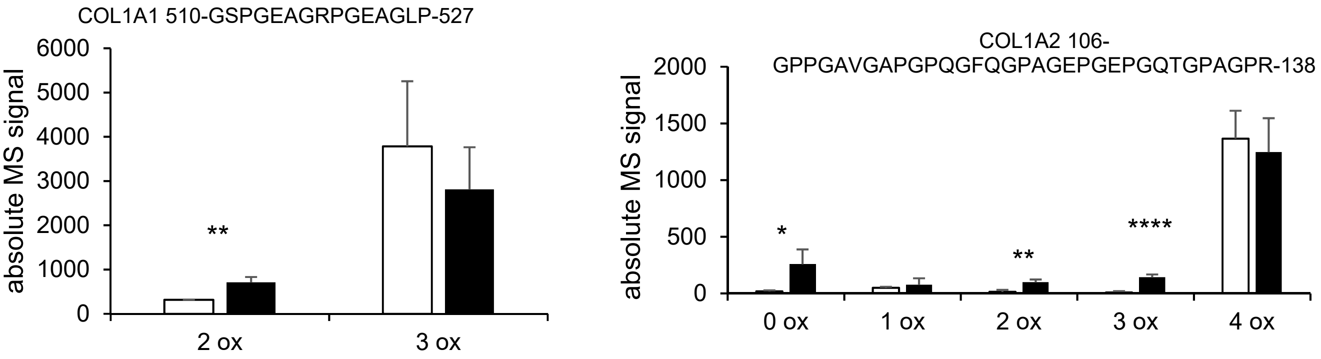


B

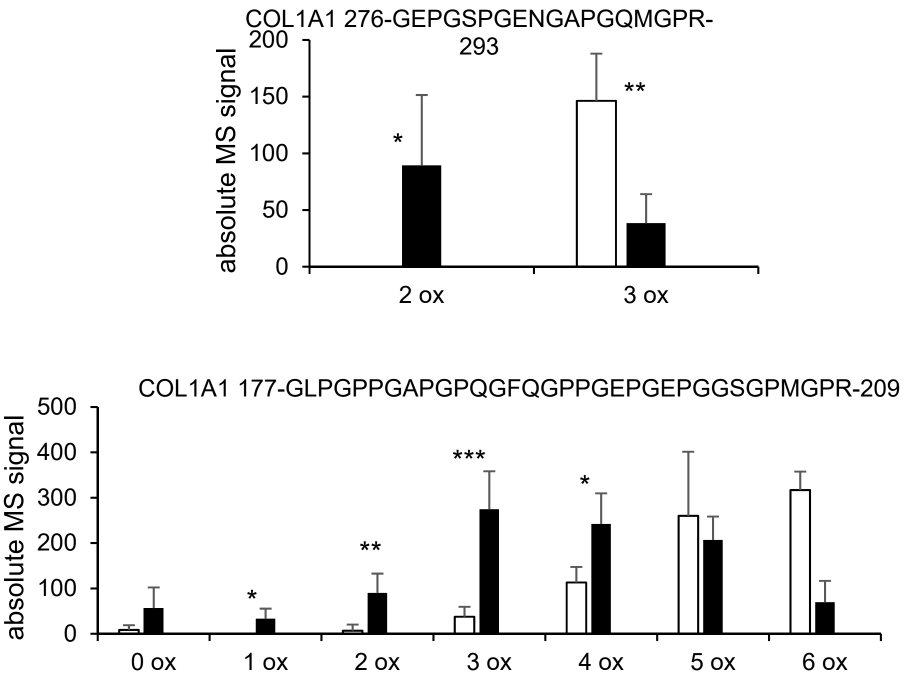

C

| | residue positions | sequence | average hydroxylation # wt | P4HA 2 ko | p-value |
|---|---|---|---|---|---|
| **COL1A1** | 177 - 209 | GLPGPPGAPGPQGFQGPPGEPGEPGGSGPMGPR | 5.11 | 3.58 | 2.24E-5 |
| | 276 - 293 | GEPGSPGENGAPGQMGPR | 3 | 2.41 | 3.14E-2 |
| | 438 - 458 | GEPGATGVQGPPGPAGEEGKR | 1.85 | 1.41 | 1.20E-4 |
| | 462 - 476 | GEPGPSGLPGPPGER | 3 | 2.6 | 8.37E-5 |
| | 510 - 527 | GSPGEAGRPGEAGLPGAK | 2.92 | 2.79 | 2.71E-2 |
| | 816 - 824 | GEPGDTGVK | ND | | |
| | 984 - 1,003 | GPPGPMGPPGLAGPPGESGR | 2.62 | 2.27 | NS |
| **COL1A2** | 106 - 138 | GPPGAVGAPGPQGFQGPAGEPGEPGQTGPAGPR | 4 | 3.94 | 1.46E-4 |
| | 205 - 222 | GEPGAPGENGTPGQAGAR | 3 | 2.4 | NS |
| | 367 - 387 | GEPGSVGAQGPPGPSGEEGKR | ND | | |
| | 439 - 453 | GPNGDAGRPGEPGLM | 2 | 1.92 | 2.72E-2 |
| | 505 - 516 | GPSGDPGKPGER | 0.55 | 0.43 | NS |
| | 580 - 594 | GLPGEFGLPGPAGPR | 2.12 | 1.85 | 7.69E-4 |
| | 853 - 885 | GPSGEPGTAGAPGTAGPQGLLGAPGILGLPGSR | 4 | 3.88 | 2.88E-6 |
| | 889 - 912 | GLPGIAGALGEPGPLGISGPPGAR | 2.98 | 2.45 | 3.22E-5 |
| | 985 - 1,002 | GEPGPAGSVGPVGAVGPR | 1 | 0.68 | 3.34E-5 |
