## supplemental data S5 for "Tissue-specific collagen hydroxylation at GEP/GDP triplets mediated by P4HA2"

**Supplemental data S5. Hydroxylation quantification regulations found upon nickel treatment of ATDC5 cells.** Measurements were done from the culture supernatants. (A) tendon-related peptides. (B) tendon-unregulated peptides. Throughout this figure, *, **, ***, ****, ***** indicate t-test p-values < 0.05, 0.005, 0.0005, 0.00005 or 0.000005 versus control and n =3.

A

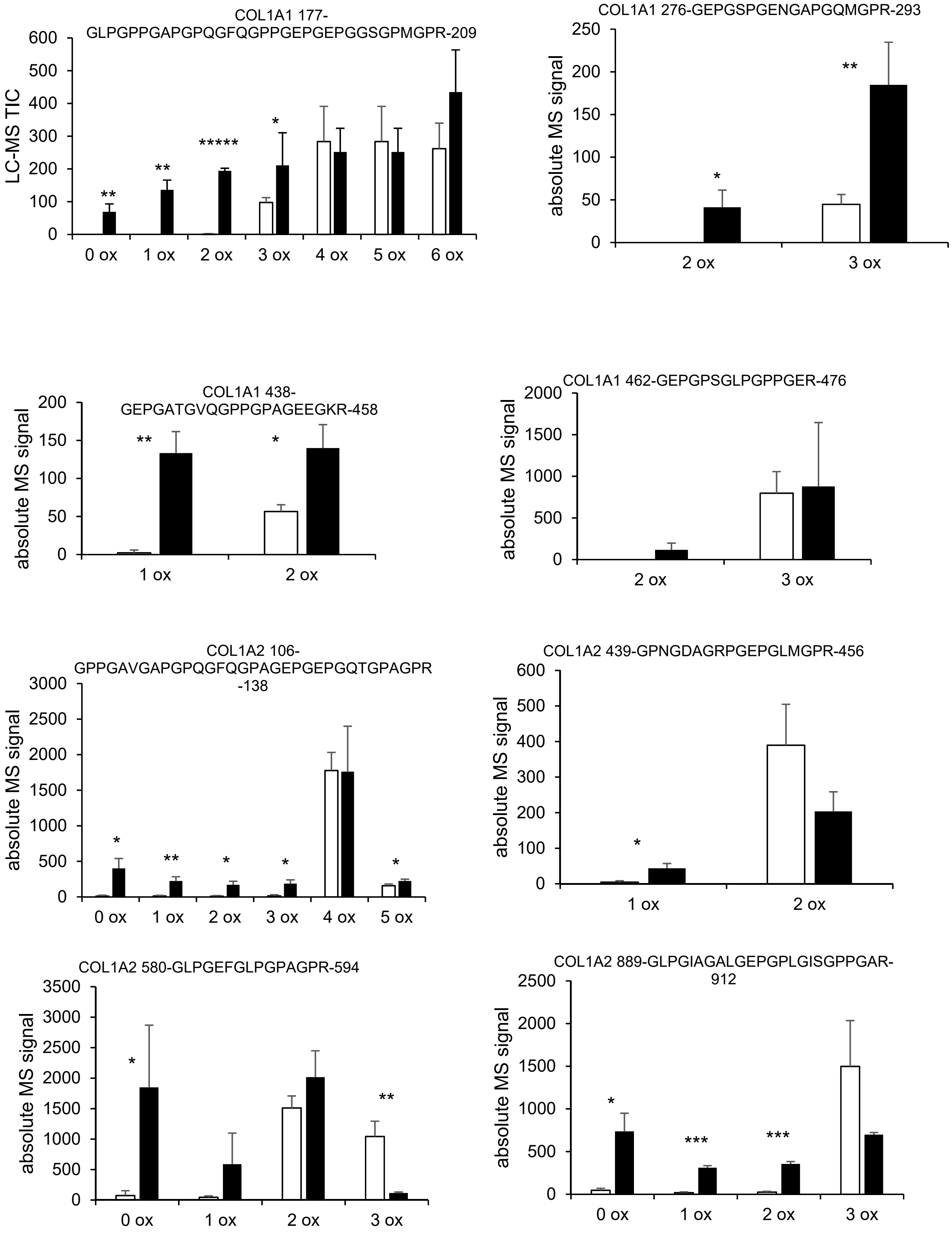

B

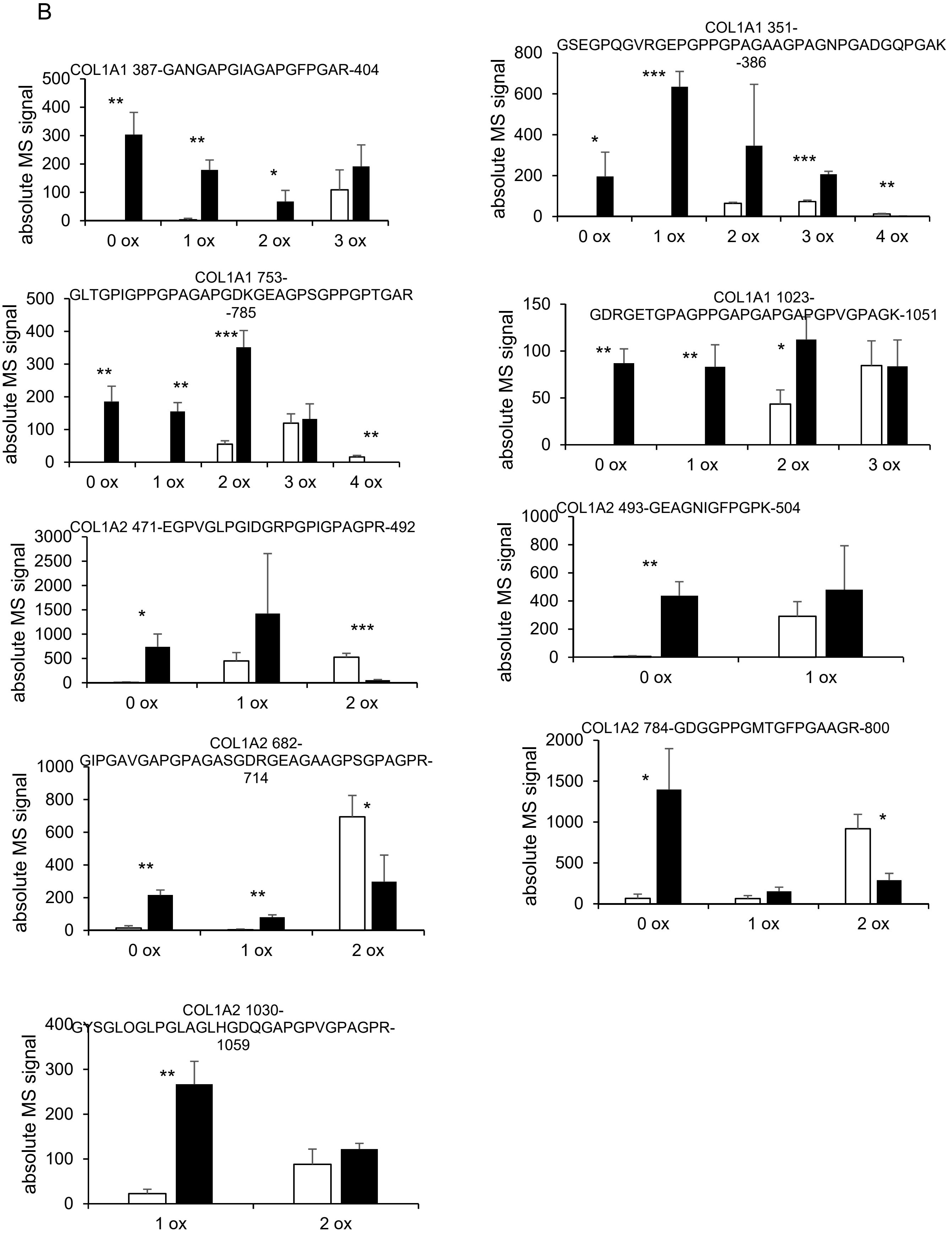
COL1A1 387-GANGAPGIAGAPGFPGAR-404
absolute MS signal
0 ox
1 ox
2 ox
3 ox
COL1A1 351-GSEGPQGVRGEPGPPGPAGAAGPAGNPGADGQPGAK-386
absolute MS signal
0 ox
1 ox
2 ox
3 ox
4 ox
COL1A1 753-GLTGPIGPPGPAGAPGDKGEAGPSGPPGPTGAR-785
absolute MS signal
0 ox
1 ox
2 ox
3 ox
4 ox
COL1A1 1023-GDRGETGPAGPPGAPGAPGAPGPVGPAGK-1051
absolute MS signal
0 ox
1 ox
2 ox
3 ox
COL1A2 471-EGPVGLPGIDGRPGPIGPAGPR-492
absolute MS signal
0 ox
1 ox
2 ox
COL1A2 493-GEAGNIGFPGPK-504
absolute MS signal
0 ox
1 ox
COL1A2 682-GIPGAVGAPGPAGASGDRGEAGAAGPSGPAGPR-714
absolute MS signal
0 ox
1 ox
2 ox
COL1A2 784-GDGGPPGMTGFPGAAGR-800
absolute MS signal
0 ox
1 ox
2 ox
COL1A2 1030-GYSGLOGLPGLAGLHGDQGAPGPVGPAGPR-1059
absolute MS signal
1 ox
2 ox
