## supplemental data S6 for "Tissue-specific collagen hydroxylation at GEP/GDP triplets mediated by P4HA2"

**Supplemental data S6. Sequence alinement of mouse α1 (col1a1) and α2 (col1a2) chains of type I collagen.** Residues are annotated according to their position in mature chains. Strictly conserved residues are boxed in black. Functionally similar residues are boxed in grey. GEP/GDP triplets found under-hydroxylated in this study are annotated in red. The one found strictly hydroxylated in all tissues is annotated in green. The ones for which the current data do not allow conclusions or suggest no regulation are annotated in orange. Non-GEP positions found significantly under-hydroxylated in tendon are annotated in light blue.

```
col1a1    1 QMSYGYDEKSAGVSVPGPMGPSGPRGLPGPPGAPGPQGFQGPPGEPGEPGGSGPMGPRGP
col1a2    1 -----QYSDKGVSSGPGPMGLMGPRGPPGAVGAPGPQGFQGPAGEPGEPGQTGPAGPRGP
                                                                       P
col1a1   61 PGPPGKNGDDGEAGKPGRPGERGPPGPQGARGLPGTAGLPGMKGHRGFSGLDGAKGDAGP
col1a2   56 AGSPGKAGEDGHPGKPGRPGERGVVGPQGARGFPGTPGLPGFKGVKGHSGMDGLKGQPGA
            XGP                                                  PXGP
col1a1  121 AGPKGEPGSPGENGAPGQMGPRGLPGERGRPGPPGTAGARGNDGAVGAAGPPGPTGPTGP
col1a2  116 QGVKGEPGAPGENGTPGQAGARGLPGERGRVGAPGPAGARGSDGSVGPVGPAGPIGSAGP

col1a1  181 PGFPGAVGAKGEAGPQGARGSEGPQGVRGEPGPPGPAGAAGPAGNPGADGQPGAKGANGA
col1a2  176 PGFPGAPGPKGELGPVGNPGPAGPAGPRGEVGLPGLSGPVGPPGNPGTNGLTGAKGATGL
                             PXGP
col1a1  241 PGIAGAPGFPGARGPSGPQGPSGPPGPKGNSGEPGAPGNKGDTGAKGEPGATGVQGPPGP
col1a2  236 PGVAGAPGLPGPRGIPGPAGAAGATGARGLVGEPGPAGSKGESGNKGEPGSVGAQGPPGP

col1a1  301 AGEEGKRGARGEPGPSGLPGPPGERGGPGSRGFPGADGVAGPKGPSGERGAPGPAGPKGS
col1a2  296 SGEEGKRGSPGEAGSAGPAGPPGLRGSPGSRGLPGADGRAGVMGPPGNRGSTGPAGIRGP

col1a1  361 PGEAGRPGEAGLPGAKGLTGSPGSPGPDGKTGPPGPAGQDGRPGPAGPPGARGQAGVMGF
col1a2  356 NGDAGRPGEPGLMGPRGLPGSPGNVGPSGKEGPVGLPGIDGRPGPIGPAGPRGEAGNIGF

col1a1  421 PGPKGTAGEPGKAGERGLPGPPGAVGPAGKDGEAGAQGAPGPAGPAGERGEQGPAGSPGF
col1a2  416 PGPKGPSGDPGKPGERGHPGLAGARGAPGPDGNNGAQGPPGPQGVQGGKGEQGPAGPPGF

col1a1  481 QGLPGPAGPPGEAGKPGEQGVPGDLGAPGPSGARGERGFPGERGVQGPPGPAGPRGNNGA
col1a2  476 QGLPGPSGTTGEVGKPGERGLPGEFGLPGPAGPRGERGTPGESGAAGPSGPIGSRGPSGA

col1a1  541 PGNDGAKGDTGAPGAPGSQGAPGLQGMPGERGAAGLPGPKGDRGDAGPKGADGSPGKDGA
col1a2  536 PGPDGNKGEAGAVGAPGSAGASGPGGLPGERGAAGIPGGKGEKGETGLRGDTGNTGRDGA

col1a1  601 RGLTGPIGPPGPAGAPGDKGEAGPSGPPGPTGARGAPGDRGEAGPPGPAGFAGPPGADGQ
col1a2  596 RGIPGAVGAPGPAGASGDRGEAGAAGPSGPAGPRGSPGERGEVGPAGPNGFAGPAGAAGQ

col1a1  661 PGAKGEPGDTGVKGDAGPPGPAGPAGPPGPIGNVGAPGPKGPRGAAGPPGATGFPGAAGR
col1a2  656 PGAKGEKGTKGPKGENGIVGPTGSVGAAGPSGPNGPPGPVGSRGDGGPPGMTGFPGAAGR

col1a1  721 VGPPGPSGNAGPPGPPGPVGKEGGKGPRGETGPAGRPGEVGPPGPPGPAGEKGSPGADGP
col1a2  716 TGPPGPSGIAGPPGPPGAAGKEGIRGPRGDQGPVGRTGETGASGPPGFVGEKGPSGEPGT

col1a1  781 AGSPGTPGPQGIAGQRGVVGLPGQRGERGFPGLPGPSGEPGKQGPSGSSGERGPPGPMGP
col1a2  776 AGAPGTAGPQGLLGAPGILGLPGSRGERGLPGIAGALGEPGPLGISGPPGARGPPGAVGS

col1a1  841 PGLAGPPGESGREGSPGAEGSPGRDGAPGAKGDRGETGPAGPPGAPGAPGAPGPVGPAGK
col1a2  836 PGVNGAPGEAGRDGNPGSDGPPGRDGQPGHKGERGYPGSIGPTGAAGAPGPHGSVGPAGK

col1a1  901 NGDRGETGPAGPAGPIGPAGARGPAGPQGPRGDKGETGEQGDRGIKGHRGFSGLQGPPGS
col1a2  896 HGNRGEPGPAGSVGPVGAVGPRGPSGPQGIRGDKGEPGDKGHRGLPGLKGYSGLQGLPGL

col1a1  961 PGSPGEQGPSGASGPAGPRGPPGSAGSPGKDGLNGLPGPIGPPGPRGRTGDSGPAGPPGP
col1a2  956 AGLHGDQGAPGPVGPAGPRGPAGPSGPVGKDGRSGQPGPVGPAGVRGSQGSQGPAGPPGP

col1a1 1021 PGPPGPPGPPSGGYDFSFLPQPPQEKSQDGGRYY
col1a2 1016 PGPPGPPGVSGGGYDFGFEGDFYRA---------
```
